# Cell type-specific dysregulation of gene expression due to *Chd8* haploinsufficiency during mouse cortical development

**DOI:** 10.1101/2024.08.14.608000

**Authors:** Kristina M. Yim, Marybeth Baumgartner, Martina Krenzer, María F. Rosales Larios, Guillermina Hill-Terán, Timothy Nottoli, Rebecca A. Muhle, James P. Noonan

## Abstract

Disruptive variants in the chromodomain helicase *CHD8*, which acts as a transcriptional regulator during neurodevelopment, are strongly associated with risk for autism spectrum disorder (ASD). Loss of CHD8 function is hypothesized to perturb gene regulatory networks in the developing brain, thereby contributing to ASD etiology. However, insight into the cell type-specific transcriptional effects of CHD8 loss of function remains limited. We used single-cell and single-nucleus RNA-sequencing to globally profile gene expression and identify dysregulated genes in the embryonic and juvenile wild type and *Chd8^+/−^* mouse cortex, respectively. *Chd8* and other ASD risk-associated genes showed a convergent expression trajectory that was largely conserved between the mouse and human developing cortex, increasing from the progenitor zones to the cortical plate. Genes associated with risk for neurodevelopmental disorders and genes involved in neuron projection development, chromatin remodeling, signaling, and migration were dysregulated in *Chd8^+/−^* embryonic day (E) 12.5 radial glia. Genes implicated in synaptic organization and activity were dysregulated in *Chd8^+/−^* postnatal day (P) 25 deep- and upper-layer excitatory cortical neurons, suggesting a delay in synaptic maturation or impaired synaptogenesis due to CHD8 loss of function. Our findings reveal a complex pattern of transcriptional dysregulation in *Chd8^+/−^* developing cortex, potentially with distinct biological impacts on progenitors and maturing neurons in the excitatory neuronal lineage.

## Introduction

Whole exome and whole genome studies have identified rare, protein-damaging genetic variants in over 200 genes that contribute to an increased risk of an autism spectrum disorder (ASD) diagnosis^1–8^. The majority of these genes encode proteins involved in transcriptional regulation, such as chromatin modifiers and transcription factors, or proteins involved in neuronal communication^2^. These findings suggest that ASD risk-associated genes converge in transcriptional networks and biological pathways that are disrupted by variants that perturb ASD risk gene function, thereby contributing to ASD etiology.

Gene expression studies in fetal and adult human brain support this hypothesis. Longitudinal analyses of transcription across developing brain regions have identified co-expression networks in the human mid-fetal cortex that are enriched in ASD risk-associated genes^9–11^. Single-cell transcriptome studies of human cortical development further indicate that ASD risk-associated genes show enriched expression in excitatory and inhibitory neuronal lineages^2^. Single-nucleus transcriptome analysis of cortical tissues from people diagnosed with ASD also support that perturbation of gene expression is a feature of ASD^12^. Understanding how disruptions in ASD risk gene function alter gene expression in specific cell types is therefore essential for identifying the molecular and neurodevelopmental mechanisms contributing to ASD.

Efforts to address this question have relied on genetic and experimental perturbation of individual ASD risk-associated genes in model systems. Several of these studies have focused on the chromatin remodeler *CHD8*, which is among the most significant ASD risk-associated genes yet discovered in rare variant screens^2,8,13^. ASD risk-associated protein-truncating variants in *CHD8* are hypothesized to lead to *CHD8* loss of function, resulting in changes in CHD8 target gene expression in the developing brain^14–17^. Maps of CHD8 binding profiles in the human and mouse developing brain showed that other ASD risk-associated genes are significantly overrepresented among CHD8 gene targets^18^. Knockdown of *CHD8* expression in human neural stem cells also resulted in dysregulation of CHD8-bound ASD risk-associated genes^18,19^. These results suggest that disruption of *CHD8* contributes to ASD risk in part through its regulation of other ASD risk-associated genes.

The effects of *Chd8* loss of function on gene expression and neurodevelopment have also been studied using mouse and cortical organoid knockout models. To date, analyses of expression changes in *Chd8^+/−^*mouse models have been limited by a lack of cellular resolution and have yet to reveal a consistent pattern of transcriptional disruption. Bulk transcriptome studies in the developing mouse brain point to mild effects on transcription overall and have identified abnormal activation of the REST pathway, transcriptional changes in RNA processing pathways, and disruption of cell adhesion and axon guidance pathways^14,17,20^.

Mouse and organoid knockout studies have also revealed a variety of cell type-specific effects due to *Chd8* loss of function. Disruption of *Chd8* expression in mouse oligodendrocyte precursor cells alters oligodendrocyte lineage development and leads to myelination defects^21,22^. Disruption of *Chd8* via Perturb-Seq in the mouse embryonic brain also altered gene expression in oligodendrocyte precursor cells^23^. *Chd8* knockdown has been shown to disrupt neural progenitor proliferation in the mouse embryonic cortex, and CRISPR-Cas9 knockout of *CHD8* reduces proliferation of cultured human neural stem cells, although this may reflect differential effects on cellular viability at very low levels of *CHD8* expression^24,25^. *CHD8^+/−^* cultured excitatory neurons derived from human embryonic stem cells have been reported to exhibit decreased synaptic activity^26^. An increase in the number of *Pax6*-positive cells has also been reported in the *Chd8^+/−^* developing mouse cortex, as has a massive increase in GABAergic interneurons in *CHD8^+/−^* human cortical organoids^17,27^. However, a consistent molecular phenotype has yet to be established across these systems.

These results, coupled with the overall cellular heterogeneity of the developing cortex, suggest there may be cell type-specific variation in the effects of *Chd8* loss of function. Such changes could be obscured by bulk transcriptome analyses. To address this question, we used single-cell RNA-sequencing (scRNA-seq) and single-nucleus RNA-sequencing (snRNA-seq) to characterize global gene expression at single-cell resolution during embryonic and juvenile cortical development in wild type and *Chd8^+/−^* mice. We found that CHD8 exhibits a clear expression gradient in the developing mouse cortex, increasing from the germinal zone to cortical plate. *Chd8* and other genes associated with risk for ASD and other neurodevelopmental disorders show convergent expression trajectories across the developing mouse cortex, and this convergent pattern is conserved between mouse and human cortical development. Our results also support that loss of *Chd8* expression results in dysregulation of genes associated with risk for neurodevelopmental disorders and genes potentially regulated by CHD8 in the embryonic cortex, particularly in embryonic day (E) 12.5 radial glia. We also found that genes involved in synaptic organization, synaptic signaling, and synaptic activity are specifically dysregulated in both deep- and upper-layer excitatory neurons in the postnatal day (P) 25 *Chd8^+/−^* cortex, including multiple glutamate receptor subunits and their regulators. This signature may indicate a delay in synaptic maturation, impaired synaptogenesis, or both. Collectively, our results point to a complex pattern of transcriptional disruption in the excitatory neuronal lineage, impacting both early progenitors and maturing neurons, due to loss of *Chd8* expression.

## Results

### Generation and initial characterization of a Chd8^+/−^ mouse model

To investigate how CHD8 loss of function may affect both CHD8 expression and gene expression overall within individual cortical cell types, we generated a *Chd8* loss of function mouse model. We used CRISPR editing to generate a *Chd8^+/−^* mutant mouse carrying a constitutive deletion of *Chd8* exon 3 (Fig. 1A; Methods). This mutation results in a frameshift at Ala408 predicted to result in premature termination of the CHD8 protein. We found that CHD8 protein levels were significantly reduced in the E16.0 *Chd8^+/−^* cortex (Fig. 1B, Table S1). *Chd8^−/−^* null embryos were not obtained from *Chd8^+/−^* in-crosses, recapitulating the finding that homozygous loss-of-function of *Chd8* is incompatible with life^14,17,28,29^.

**Figure 1.**
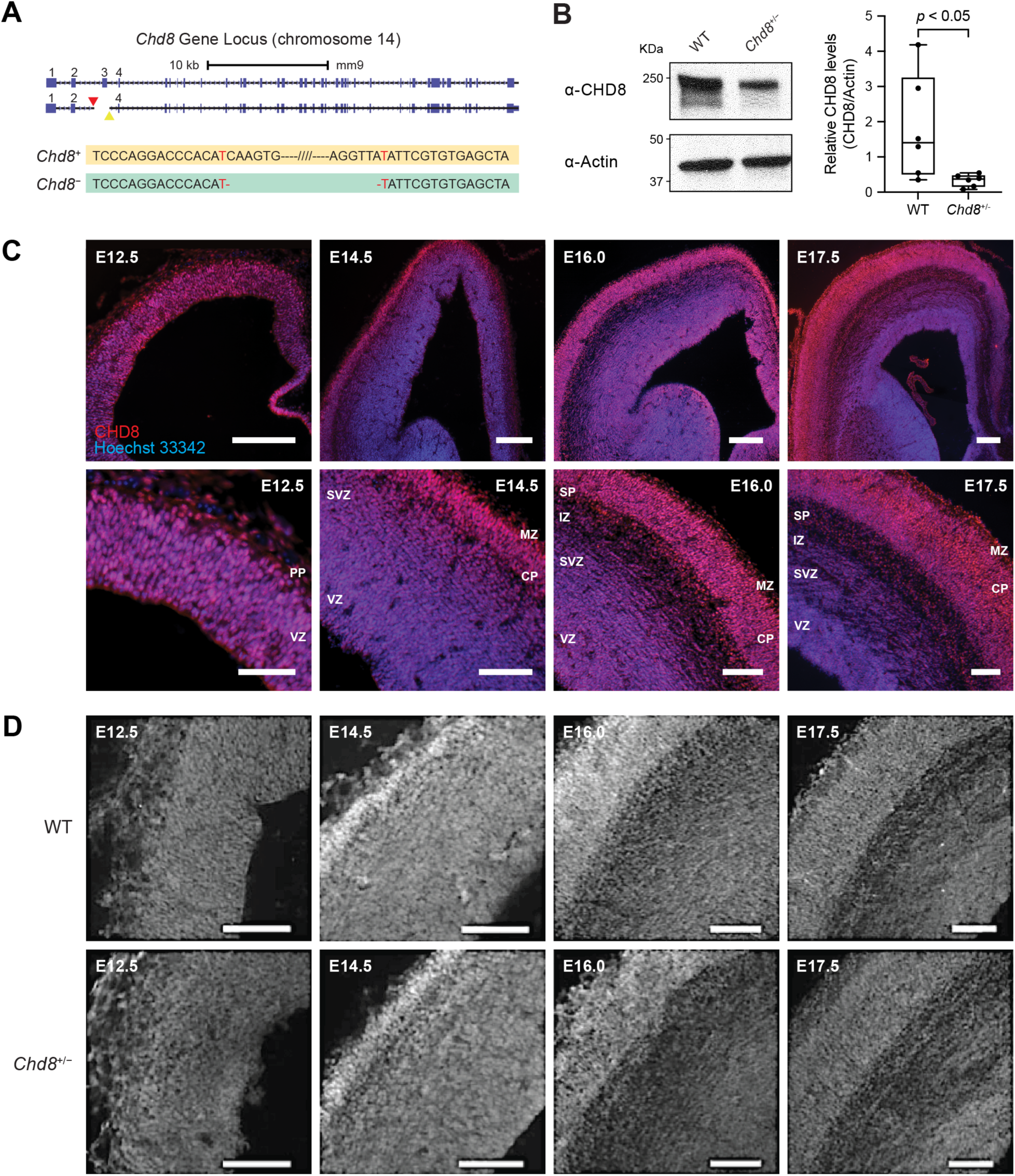
Generation and characterization of a *Chd8^+/−^* mouse model. (**A**) Schematic showing our strategy for constitutive *Chd8^+/−^* mouse generation, using CRISPR-Cas9 targeted mutagenesis to delete 1,061 nucleotides (chr14:52,852,109-52,853,169, mm9) that include exon 3, causing a frameshift at alanine 408. The *Chd8* gene models (*top*) and comparison of the targeted sequence (*bottom*) in the wild type (*Chd8*^+^) and disrupted (*Chd8*^-^) alleles are shown; nucleotides directly adjacent to the deletion site are colored red. Triangles denote expected guide RNA (gRNA)-directed cleavage sites of the upstream (red) and downstream (yellow) gRNAs (Methods). The dashed line with slashes indicates sequence omitted for clarity. (**B**) Western blot showing CHD8 and actin expression in male wild type (WT) and *Chd8^+/−^* embryonic day (E) 16.0 cortices (*left*), with quantification of CHD8 expression in the wild type and *Chd8^+/−^* E16.0 cortex (*right*). CHD8 levels were normalized to actin in each replicate (Methods). Whiskers span the minimum and maximum data points, boxes span the interquartile interval, and lines indicate the median. Significance was determined by one-tailed Welch’s *t*-test (Methods). (**C**) Coronal sections of wild type embryonic mouse cortex at the indicated stages, stained with anti-CHD8 antibody (red) and Hoechst 33342 (nuclei; blue). Scale bars: 200μm (*top*), 50μm (*bottom*). (**D**) Spatiotemporal expression of CHD8 (grayscale) in coronal sections of the wild type (*top*) and *Chd8^+/−^* (*bottom*) embryonic mouse cortex collected from littermates at the indicated stages. Scale bar: 100µm. VZ = ventricular zone; SVZ = subventricular zone; IZ = intermediate zone; PP = preplate; SP = subplate; CP = cortical plate; MZ = marginal zone. See also Figures S1-S2 and Table S1.

We first assessed the spatiotemporal distribution of CHD8 expression in wild type and *Chd8^+/−^* mice at four time points during embryonic cortical development (E12.5, E14.5, E16.0, E17.5) (Fig. 1C-D; Methods). At E12.5, CHD8 exhibited a uniform nuclear expression pattern across the cortex, from neural progenitor cells in the ventricular zone (VZ) to neurons in the preplate (PP). At E14.5, CHD8 expression showed a clear apical-to-basal expression gradient, with increased expression in the cortical plate (CP) compared to progenitor cells of the VZ and subventricular zone (SVZ). This expression gradient persisted at E16.0 and E17.5, with increased expression in neurons of the subplate (SP) and CP compared to the germinal zones. We validated these CHD8 expression patterns using secondary antibody-only negative controls (Fig. S1A). CHD8 expression was consistent between male and female embryos at each developmental stage (Fig. S1B) as well as along the rostral-caudal axis of the cortex (Fig. S2). The distribution of CHD8 expression in the developing CP was also similar in wild type and *Chd8^+/−^* littermates at each of the four embryonic time points we examined (Fig. 1D).

### Single-cell transcriptome profiling of wild type and Chd8^+/−^ mouse embryonic cortex

We next used scRNA-seq to quantify gene expression in wild type and *Chd8*^+/−^ mouse cortices at the same embryonic time points examined in the IHC studies described above (Fig. 1C-D). We generated 32 scRNA-seq datasets from sex- and litter-matched pairs of wild type and *Chd8^+/−^* cortices (eight embryos per time point), capturing single-cell expression profiles for 145,082 total cells (Fig. 2A, Table S2; Methods). Genotyping results for each sample are presented in Figure S3. We removed low quality cells and potential doublets by thresholding on the total feature counts per cell, the number of genes detected per cell, and the percentage of counts originating from mitochondrial RNA, resulting in 135,926 cells after filtering (Fig. S4, Table S2; Methods).

**Figure 2.**
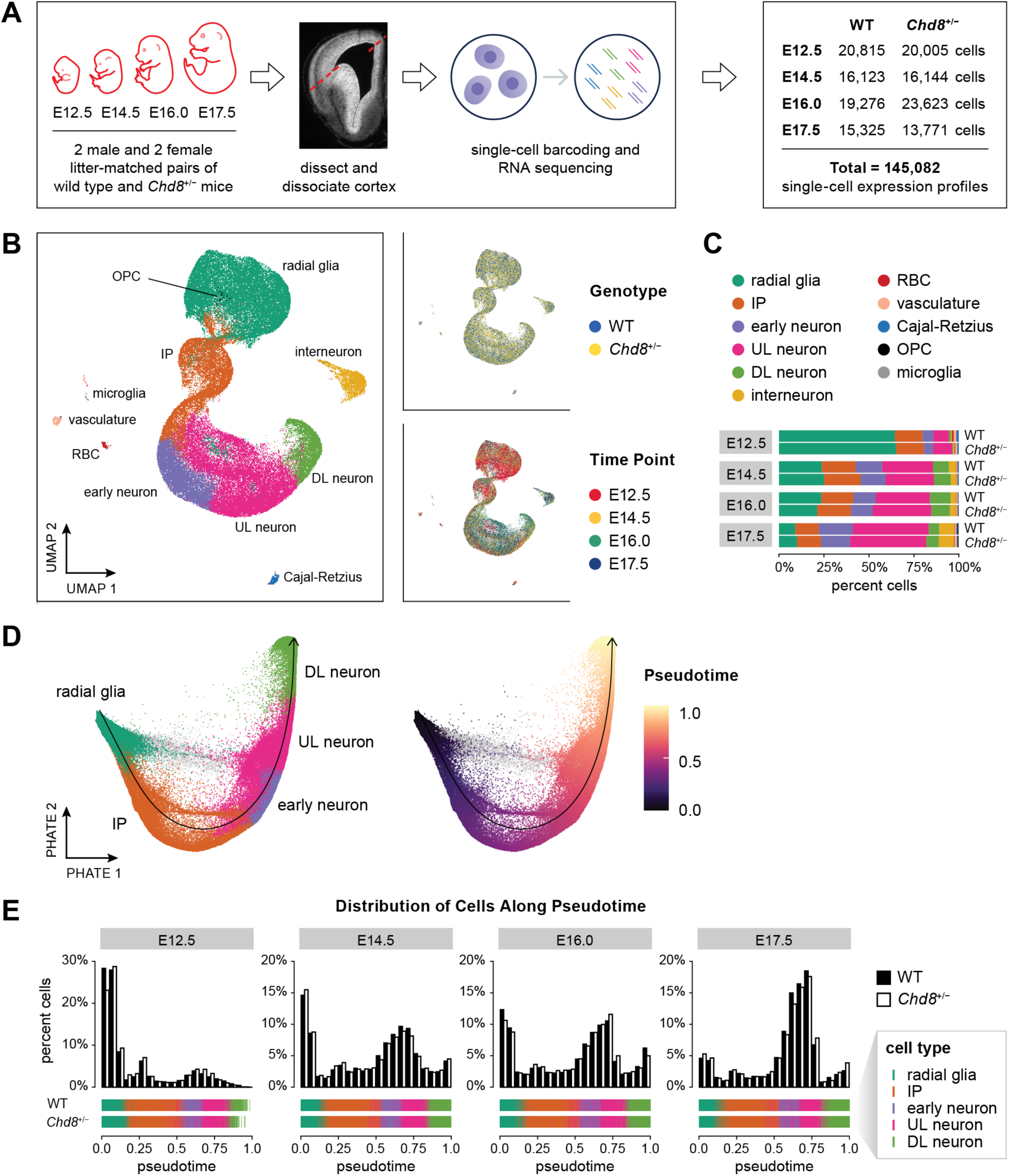
Analysis of embryonic cortical development using single-cell RNA-sequencing in wild type (WT) and *Chd8*^+/−^ mice. (**A**) Study design, including dissection schema and number of cells collected at each time point in each genotype. (**B**) UMAP embedding of 135,926 cells colored by cell type clusters (*left*), genotype (*top right)*, and time point (*bottom right*). Cell type clusters were identified using a graph-based clustering approach, implemented in Seurat, and then annotated based on the expression of known cell type marker genes (Methods). (**C**) Cell type composition at each time point in each genotype. (**D**) PHATE embedding of 128,112 cells in the excitatory neuronal lineage, colored by cell type (*left*) and pseudotime (*right)*. The principal curve through the primary trajectory of the PHATE embedding is shown as a black arrow. A set of 4,457 radial glia-like cells (shown in gray), mostly present at embryonic day (E) 12.5, diverged from the primary trajectory and were excluded from further analyses. (**E**) Cell type representation along pseudotime. Male and female samples were aggregated for each time point and genotype; the pseudotime trajectory was divided into 20 equally-spaced bins and the percent of cells within each bin is shown, color-coded by genotype (WT: black; *Chd8*^+/−^: white). Below each distribution, the position of each cell along pseudotime is plotted as a vertical line, colored according to cell type label. See also Figures S3-S9 and Tables S2-S5. IP = intermediate progenitors; UL = upper-layer; DL = deep-layer; RBC = red blood cells; OPC = oligodendrocyte precursor cells.

To mitigate differences in cell cycle phase among dividing cells, we calculated a cell cycle score for each cell and then regressed out the difference between G2M and S phase scores (Methods). We then normalized feature counts using SCTransform, integrated all 32 datasets using Seurat, and visualized the integrated data using uniform manifold approximation and projection (UMAP) (Fig. 2B; Methods)^30–35^. Labeling cells by cell cycle phase, we observed a distinct boundary between proliferating and post-mitotic cells (Fig. S5A). Cells within the embedding did not separate by sex, genotype, or sample batch (Fig. 2B, Fig. S5A-B). The distribution of cells within the UMAP embedding was largely determined by the expression of highly variable genes, particularly markers of specific cell types such as radial glia (*Sox2, Pax6*)^36,37^, intermediate progenitors (*Eomes, Neurog2*)^38^, and neurons (*Tbr1*, *Fezf2*, *Neurod1*)^38,39^ (Fig. S6A-B & S7, Table S3).

We identified cell clusters using Seurat and then annotated clusters based on the expression of known cell type marker genes (Fig. 2B, Fig. S7-8, Table S3; Methods). The vast majority of cells (>94%) belonged to the excitatory glutamatergic neuronal lineage (Fig. 2C, Table S4). These included radial glia (expressing *Sox2*, *Pax6*), intermediate progenitors (*Eomes, Neurog2*), newborn/early neurons (*Neurod1*, *Unc5d*)^40^, upper-layer neurons (*Satb2*)^39^, and deep-layer neurons (*Fezf2*, *Tbr1*). We also identified GABAergic interneurons (∼4% of total cells, expressing *Dlx2, Gad2*)^41^, and low percentages of red blood cells, vasculature cells^42,43^, Cajal-Retzius cells^44^, oligodendrocyte precursor cells^45^, and microglia^46^ (Fig. S7-8, Table S4). Using a two-tailed Welch’s *t*-test, we compared the number of cells per cell type in the wild type and *Chd8^+/−^* cortices (Fig. S9, Table S5; Methods). In contrast to previous studies, we did not observe significant differences in cell type representation due to loss of *Chd8*^17,27^.

### Inferring gene expression trajectories in the developing mouse cortex

Our IHC results demonstrated that CHD8 exhibits a clear apical-to-basal expression gradient in the developing mouse cortex (Fig. 1C). To identify genes, including other ASD risk-associated genes, that show similar patterns of expression, we used potential of heat-diffusion affinity-based transition embedding (PHATE) to infer gene expression trajectories in our scRNA-seq datasets (Methods)^47^. For this analysis, we focused on cells in the excitatory neuronal lineage (i.e., radial glia, intermediate progenitors, newborn/early excitatory neurons, upper-layer excitatory neurons, and deep-layer excitatory neurons); other cell types were excluded (Fig. 2D). PHATE showed that the expression profiles of cells in the excitatory neuronal lineage are distributed along a continuum. Given the continuous structure of the data, we fit a principal curve to the PHATE embedding and then projected cells onto the curve to assign each cell a value on a 0 to 1 pseudotime scale (Fig. 2D-E)^48,49^. We found that cell types were ordered along the principal curve corresponding to the apical-to-basal cortical excitatory neurogenic axis (Fig. 2D-E, Fig. S6C). These findings support that the pseudotime scale we inferred from PHATE recapitulates the process of differentiation and maturation in the excitatory neuronal lineage, i.e., the primary trajectory of differentiation captured in the data.

To determine whether our approach generally identified spatial and developmental gene expression gradients in the developing mouse cortex, we inferred expression trajectories from scRNA-seq data for *Chd8*, the ASD risk-associated gene *Pogz*, and the cortical developmental markers *Pax6* and *Tbr1* (Methods). In parallel, we performed IHC in wild type mouse cortices at E14.5 and E17.5 for CHD8, POGZ, TBR1, and PAX6 to compare the transcriptional trajectories inferred from scRNA-seq data with the expression of each protein (Fig. 3; Methods). *Chd8* showed lower transcriptional expression at the beginning of the differentiation trajectory (i.e., in radial glia) and peaked at the end of the trajectory (i.e., in post-mitotic neurons in the cortical plate). These results were consistent with the spatiotemporal distribution of CHD8 we detected using IHC, namely increased expression in mature neurons in the cortical plate and subplate compared to expression in progenitor cells in the ventricular and subventricular zones (Fig. 1C, Fig. 3A). *Pogz* exhibited a similar expression gradient to *Chd8* across the excitatory neuronal lineage in the E14.5 and E17.5 wild type cortex; this finding was also supported by IHC (Fig. 3B). The trajectories we inferred for *Tbr1* and *Pax6* were also consistent with their known expression patterns and with protein expression visualized using IHC (Fig. 3C-D)^50,51^. These findings establish that our inferred transcriptional trajectories correspond to *bona fide* developmental and cell type-specific expression patterns.

**Figure 3.**
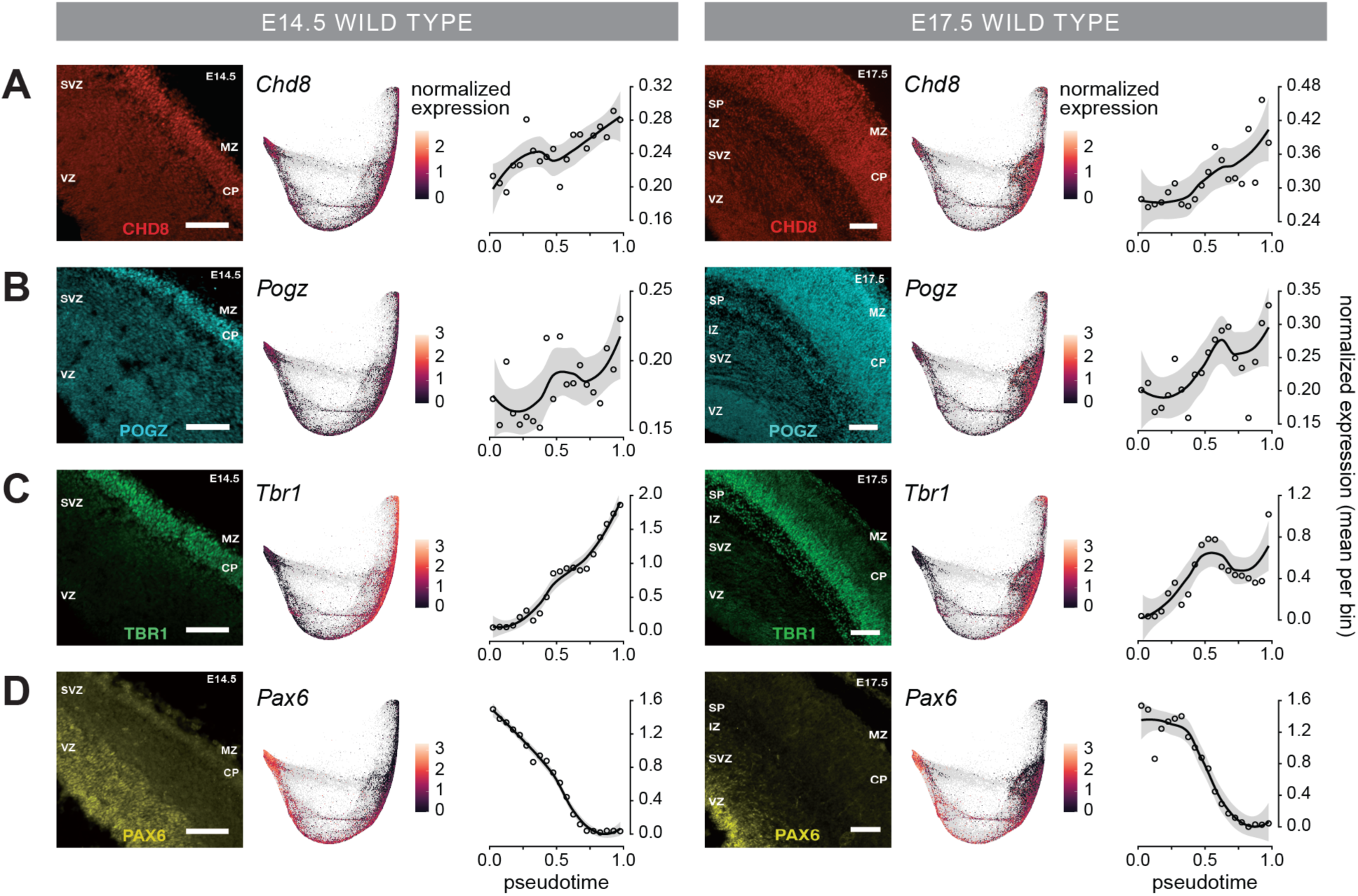
Inferring spatial and developmental gene expression gradients in the developing mouse cortex using single cell RNA-sequencing. Expression gradients for (**A**) *Chd8*, (**B**) *Pogz*, (**C**) *Tbr1*, and (**D**) *Pax6* inferred at E14.5 (*left*) and E17.5 (*right*). For each time point, the expression of each protein visualized using immunohistochemistry is shown at the left (CHD8 = red; POGZ = cyan; TBR1 = green; PAX6 = yellow; Methods). The inferred expression trajectories for each gene are shown at the right. Male and female samples were aggregated for each time point and genotype, then cells were divided into 20 equally-spaced bins along the pseudotime scale and mean expression per bin was computed for each gene. Mean expression per bin (open circles) is plotted against pseudotime. Smoothed lines are drawn with loess (span = 0.75) and a confidence interval of 0.95. See also Figure 2 and Figure S6. Scale bars: 100µm. VZ = ventricular zone; SVZ = subventricular zone; IZ = intermediate zone; SP = subplate; CP = cortical plate; MZ = marginal zone.

### Defining sets of genes in wild type and Chd8^+/−^ mouse embryonic cortex exhibiting shared transcriptional trajectories

To identify genes with similar developmental and cell type-specific expression patterns, we classified gene expression trajectories into metagenes—groups of genes with convergent expression along the pseudotime scale. We inferred gene expression trajectories from scRNA-seq data in wild type and *Chd8^+/−^* mouse cortex at E14.5, E16.0, and E17.5 (Fig. S10A-B; Methods). E12.5 expression data were excluded given the low representation of intermediate progenitors and neurons along the differentiation trajectory at this time point. We defined 10 metagene centers (labeled A through J) by *k*-means clustering wild type trajectories (*k* = 25), followed by hierarchical clustering using correlation distance (Fig. S10C-D; Methods). We then assigned each expression trajectory to a metagene based on highest Pearson correlation between the trajectory and the metagene center (Fig. S10E, Table S6).

The majority of gene expression trajectories were assigned to metagenes C, D, H, or I (Fig. 4A). These four metagenes represented the most common expression patterns we observed in the developing mouse cortex and included known cell type-specific marker genes (e.g., *Sox2* in metagene C, *Eomes* in metagene D; Fig. 4A). Metagenes H and I included genes whose expression peaks in post-mitotic neurons and revealed two distinct developmental expression patterns: genes whose expression steadily increases along the differentiation trajectory (including *Chd8*) and genes that are spatially restricted to post-mitotic neurons with no or very low expression in other cell types (e.g., *Scn2a*; Fig. 4A). The presence of shared transcriptional trajectories during development suggests that genes assigned to the same metagene may participate in common biological functions. To examine this question, we performed Gene Ontology (GO) enrichment analysis to identify biological processes that were significantly enriched in each metagene (Table S7; Methods). Metagenes C and D, i.e., genes with maximal expression in progenitor cells, were enriched for GO Biological Process (GO:BP) terms associated with metabolic and biosynthetic processes. Metagene C, which contains radial glia-specific genes, was specifically enriched for the terms “cell cycle” and “cell division” (Table S7). Metagenes H and I were both enriched for terms such as “vesicle-mediated transport” and “neuron projection development” (Fig. 4B, Table S7). Metagene H was specifically enriched for the term “autophagy,” while metagene I was specifically enriched for terms such as “chemical synaptic transmission,” “synapse organization,” and “cell-cell signaling” (Fig. 4B, Table S7).

**Figure 4.**
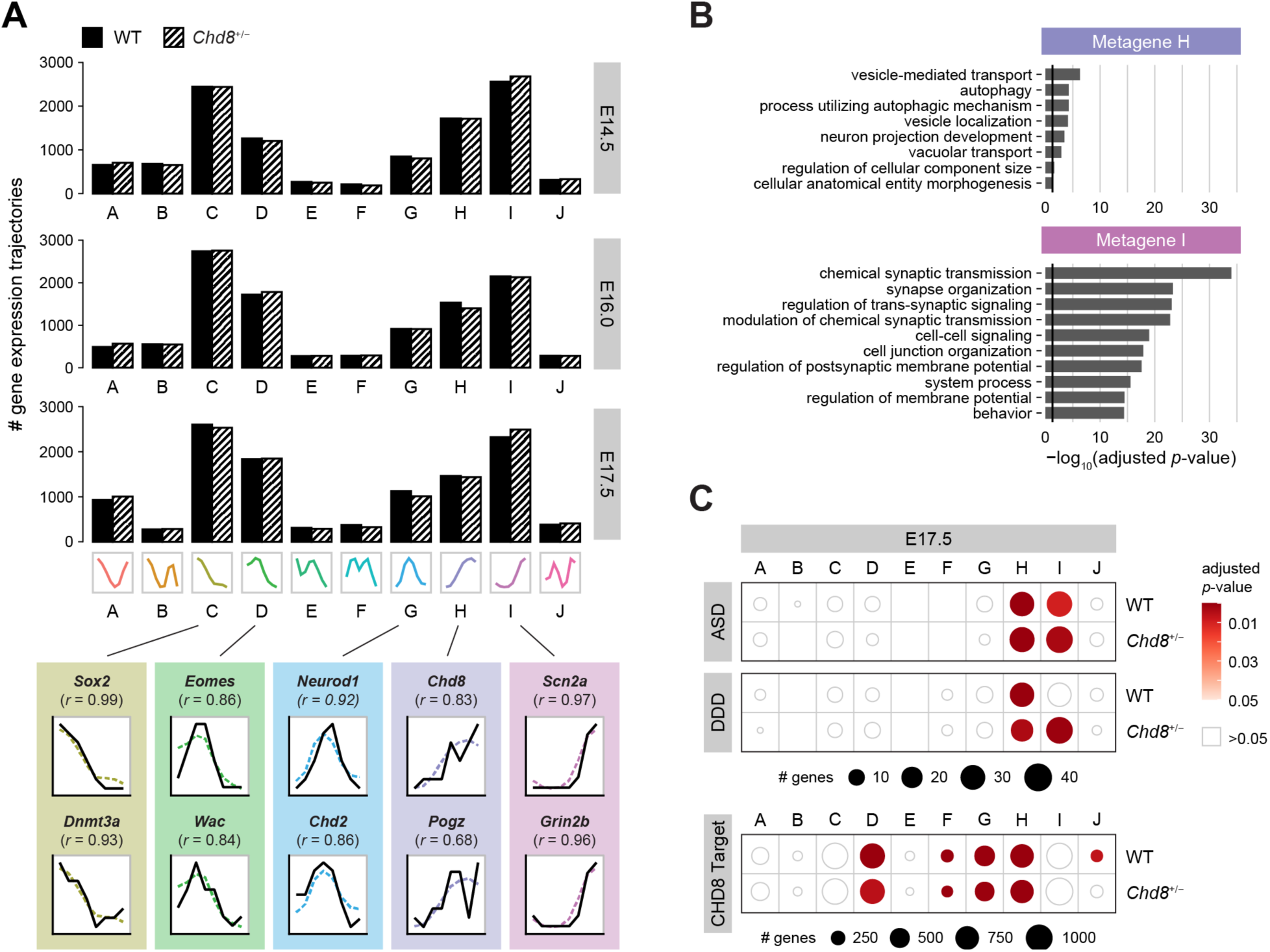
Metagenes reveal neurodevelopmental, ASD risk-associated genes, and developmental disorder risk-associated genes with common transcriptional trajectories. (**A**) Number of genes assigned to each metagene for each time point and genotype (*top*), with examples of symbolic aggregate approximation (SAX)-transformed gene expression trajectories assigned to each metagene (*bottom*). The metagene center is plotted as dashed line, and the SAX-transformed gene expression trajectory is plotted as a solid line. The Pearson correlation (*r*) between the trajectory and the metagene center is shown for each example. Example trajectories were calculated from embryonic day (E) 17.5 wild type (WT) cortex scRNA-seq data. (**B**) Gene Ontology Biological Process (GO:BP) terms significantly enriched in metagenes H and I (g:SCS-adjusted *p*-value < 0.05; black line = g:SCS-adjusted *p*-value = 0.05; Methods). GO enrichment analysis was performed using g:Profiler, and the resulting list of GO:BP terms was summarized using Revigo. Up to ten of the most significant representative GO:BP terms are shown for each metagene (Methods). (**C**) Enrichment of autism spectrum disorder risk-associated genes (ASD), the Deciphering Developmental Disorder gene set (DDD), and CHD8 binding targets in the E17.5 wild type mouse cortex for each metagene. The circle size corresponds to the number of genes within each metagene that intersect the given gene set at each time point and genotype. The circle color corresponds to the Benjamini Hochberg-adjusted *p*-value (one-tailed Fisher exact test; Methods). See also Figures S10-S12 and Tables S6-S9.

### ASD and developmental disorder risk-associated genes converge in specific metagenes in the mouse embryonic cortex

Previous studies indicate that ASD risk-associated genes are co-expressed in specific cortical regions and cell types^2,10^. To determine if ASD risk-associated genes showed convergent expression trajectories in the developing mouse cortex, we tested metagenes for enrichment for ASD risk-associated genes using a one-tailed Fisher exact test (Fig. 4C, Fig. S11A, Tables S8-S9)^2^. We also tested for enrichment of genes associated with developmental and intellectual disability identified by the Deciphering Developmental Disorders (DDD) consortium^6^. Metagene H, which includes *Chd8*, was significantly enriched for ASD risk-associated genes and DDD genes at E16.0 and E17.5 (BH-adjusted *p*-value < 0.05; Fig. 4C, Fig. S11A-B, Tables S8-S9). Metagene I was also consistently enriched for ASD risk-associated genes at E14.5, E16.0, and E17.5, and enriched for DDD genes at E16.0 (BH-adjusted *p*-value < 0.05; Fig. 4C, Fig. S11A-B, Tables S8-S9).

Risk-associated genes in metagene H tend to be involved in the regulation of gene expression and appeared to gradually increase in expression along the differentiation trajectory (e.g., *Adnp*, *Bcl11a*, *Elavl3*, *Kdm5b*, *Kmt5b*, *Kdm6b, Kmt2e*, *Myt1l*, *Pogz, Smarcc2*, *Tbr1*) (Fig. S12)^2^. In contrast, risk-associated genes in metagene I tend to function in neuronal communication and displayed cell type-specific expression in post-mitotic neurons (e.g., *Ank2, Cacna1e, Cacna2d3, Dip2a, Gabrb3, Grin2b, Kcnma1, Lrrc4c, Nrxn1, Scn2a, Shank2, Stxbp1*) (Fig. S12)^2^. Given the established role of CHD8 in gene regulation, including the regulation of other ASD risk-associated genes, we also tested whether metagenes were enriched for genes whose promoters were bound by CHD8 in the wild type E17.5 mouse cortex (Fig. S11C, Tables S8-S9)^18^. Metagene D and metagene H (which includes *Chd8*) were significantly enriched for CHD8 targets at E14.5, E16.0, and E17.5 in both wild type and *Chd8*^+/−^ mice (BH-adjusted *p*-value < 0.05; Fig. 4C, Fig. S11C). These findings suggest CHD8 may contribute to the convergent expression pattern we observed in metagene H via regulation of its target genes.

### ASD risk-associated gene trajectories are conserved between the mouse and human developing cortex

Our analysis of mouse embryonic cortex scRNA-seq data revealed developmental and cell type-specific expression patterns that were enriched for ASD risk-associated genes, DDD genes, and CHD8 binding targets. We next investigated whether similar expression patterns exist in the developing human cortex and whether these patterns are also enriched for ASD risk-associated genes. We performed pseudotime and metagene analysis on a published scRNA-seq dataset generated in the human cortex at gestational weeks 17 and 18 (Methods)^52^. As in mouse, we found that human cortical cells were ordered along the pseudotime scale based on the apical-to-basal axis of neurogenesis (Fig. 5A-B). We then inferred gene expression trajectories along the primary trajectory of differentiation in the developing human cortex and classified these trajectories into 9 metagenes using *k*-means and hierarchical clustering as previously described (Fig. 5C, Tables S10-S11).

**Figure 5.**
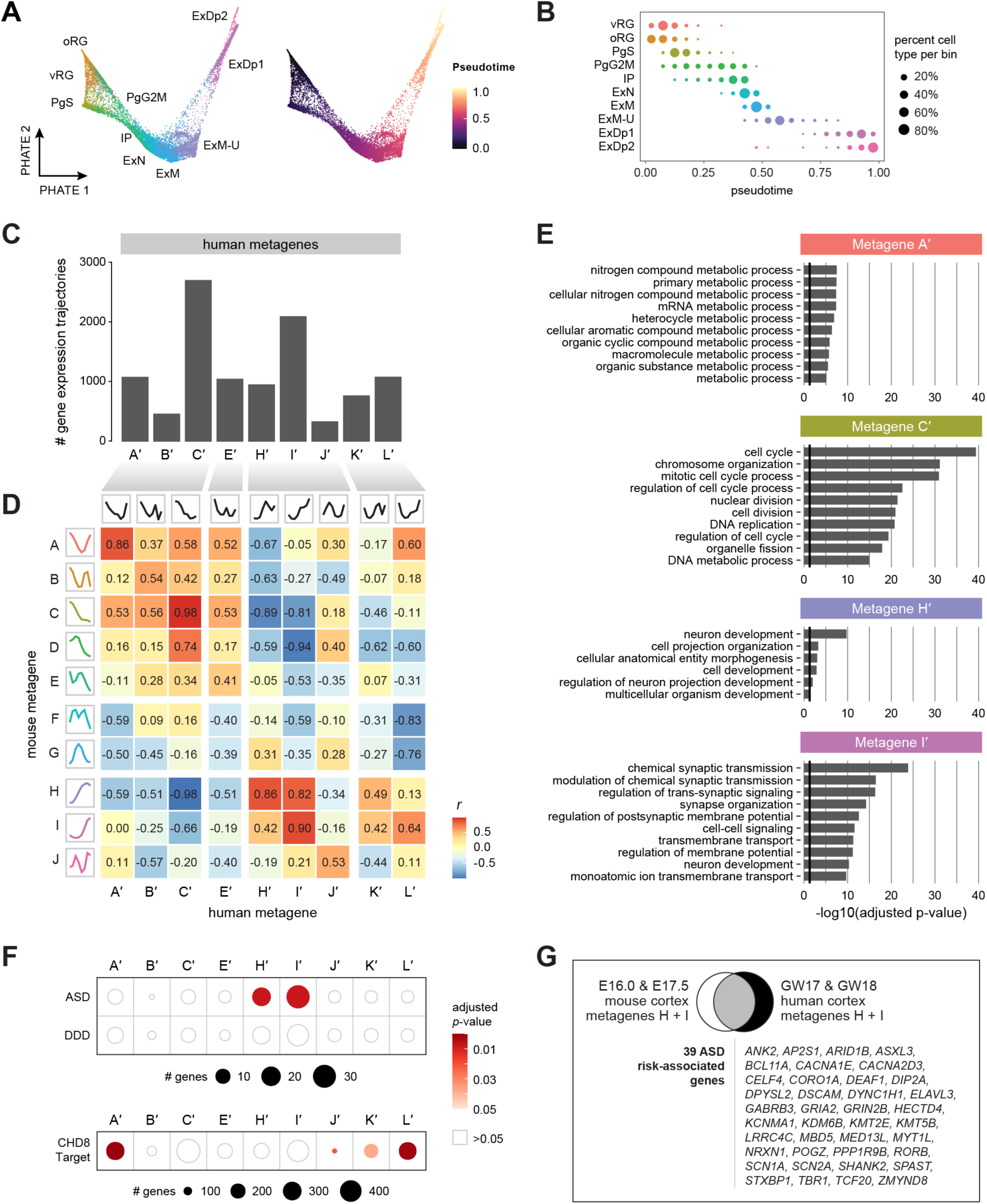
ASD risk-associated genes and developmental disorder risk-associated genes show convergent expression trajectories in mouse and human cortical development. (**A**) PHATE embedding of 30,132 cells from the fetal human cortex at gestational weeks (GW) 17 and 18, colored by cell types as defined in Polioudakis et al., 2019 (*left)* and by pseudotime (*right)*. vRG = ventral radial glia; oRG = outer radial glia; PgS = S phase cycling progenitors; PgG2M = G2M phase cycling progenitors; ExN = newborn excitatory neurons; ExM = maturing excitatory neurons; ExM-U = upper-layer-enriched maturing excitatory neurons; ExDp1 and ExDp2 = deep-layer excitatory neurons (subclusters 1 and 2). (**B**) Cell type representation in 20 bins along pseudotime, labeled as in (A). Circle size corresponds to the percent of cells of the indicated cell type per pseudotime bin. (**C**) Number of genes in each human cortex metagene. (**D**) Heat map of Pearson correlation (*r*) between mouse cortex metagene centers (labeled A through J) and human cortex metagene centers. Human metagenes were labeled based on maximal correlation between human and mouse metagene centers and denoted with a prime (′) symbol. (**E**) Gene Ontology Biological Processes (GO:BP) terms significantly enriched in human metagenes A′, C′, H′, and I′ (g:SCS-adjusted *p*-value < 0.05; black line = g:SCS-adjusted *p*-value = 0.05; Methods). GO enrichment analysis was performed using g:Profiler, and the resulting list of GO:BP terms was summarized using Revigo. Up to ten of the most significant representative GO:BP terms are shown for each metagene. (**F**) Metagene enrichment for autism spectrum disorder risk-associated genes (ASD), the Deciphering Developmental Disorder gene set (DDD), and CHD8 binding targets in the human mid-fetal cortex. Circle size corresponds to the number of genes within each metagene that intersect the given gene set. Circle color corresponds to Benjamini Hochberg-adjusted *p*-value, one-tailed Fisher exact test. (**G**) Overlap of ASD risk-associated genes identified in mouse metagenes H and I at embryonic day (E) 16.0 and E17.5, and ASD risk-associated genes identified in human metagenes H′ and I′. The 39 ASD risk-associated genes that were assigned to these metagenes in both species are shown. See also Figure S12 and Tables S6, S9, and S10-S13.

To compare metagenes identified in the human cortex to those identified in the mouse cortex, we calculated the Pearson correlation (*r*) between human and mouse metagene centers, and then we labeled human metagenes based on maximal correlation with mouse metagene centers (Fig. 5D, Table S11; Methods). The human metagenes with the highest correlation to their mouse counterparts were metagene A′ (*r* = 0.86), metagene C′ (*r* = 0.98), metagene H′ (*r* = 0.86), and metagene I′ (*r* = 0.90). Metagenes D, F, and G in mouse were not matched to a corresponding metagene in human; likewise, metagenes K′ and L′ in human did not have a corresponding metagene in mouse (Fig. 5D, Table S11).

To further assess human metagenes and compare them to the metagenes we identified in mouse, we performed GO enrichment analysis. We observed the same GO:BP terms enriched in human metagenes as their mouse counterparts: human metagene C′ was enriched for the term “cell cycle”, metagenes H′ and I′ were enriched for “neuron projection development,” and metagene I′ was enriched for “chemical synaptic transmission,” “synapse organization,” and “cell-cell signaling” (Fig. 5E, Table S12). These results suggest that highly correlated metagenes identified in the developing mouse and human cortex represent conserved developmental and cell type-specific expression patterns associated with specific biological processes.

Next, we tested human metagenes for enrichment for ASD risk-associated genes, DDD genes, and genes whose promoters were bound by CHD8 in the human mid-fetal cortex (Tables S9 & S13)^18^. We found that human metagenes H′ and I′ were both significantly enriched for ASD risk-associated genes (one-tailed Fisher exact test, Benjamini Hochberg (BH)-adjusted *p*-value < 0.05; Methods) (Fig. 5F, Tables S9 & S13). Four human metagenes were significantly enriched for CHD8 binding targets (A′, J′, K′, and L′; Fig. 5F, Tables S9 & S13). We did not observe significant enrichment for DDD genes in any human metagenes (Fig. 5F, Tables S9 & S13).

In order to identify ASD risk-associated genes with conserved expression trajectories, we compared ASD risk-associated genes in human metagenes H′ and I′ (49 genes) with ASD risk-associated genes that were consistently classified into mouse metagenes H or I at E16.0 and E17.5 (58 genes). Of these, 39 ASD risk-associated genes were present in these metagenes in both species (Fig. 5G). Similar to mouse, human cortex metagene H′ included risk-associated genes that are involved in the regulation of gene expression (e.g., *ASXL3*, *ELAVL3*, *KDM6B*, *KMT2E*, *KMT5B*, *MED13L*, *MYT1L*, *POGZ*, *RORB*, and *TCF20*) and appeared to be more broadly expressed across the developing cortex, peaking in post-mitotic neurons^2^. Human metagene I′ included risk-associated genes that are involved in neuronal communication (e.g., *AP2S1*, *CACNA1E*, *CACNA2D3*, *DIP2A*, *DSCAM*, *GABRB3*, *GRIA2*, *GRIN2B*, *KCNMA1*, *LRRC4C*, *NRXN1*, *SCN1A*, *SCN2A*, *STXBP1*) and whose expression was more spatially restricted to post-mitotic neurons^2^. Collectively, these findings indicate that ASD risk-associated gene expression trajectories are conserved between human and mouse cortical development at the time points we examined, and they converge in correlated developmental and cell type-specific expression gradients.

### Genes associated with risk for neurodevelopmental disorders are dysregulated in the Chd8^+/−^ embryonic mouse cortex

We next sought to characterize cell type-specific changes in gene expression associated with *Chd8* loss of function. For each embryonic time point, we identified differentially expressed genes (DEGs) between wild type and *Chd8^+/−^*cells in the excitatory neuronal lineage using Monocle 3 (Methods)^49,53^. We first identified DEGs across all cells in the primary trajectory as defined above (Fig. 2D, Tables S14-S15). The most DEGs were identified at E12.5 and E17.5 (2,972 and 1,743 respectively; BH-adjusted *p*-value < 0.05; Methods) (Fig. S13A). Nearly all DEGs showed only small changes in expression, i.e., less than 1.5-fold change (Fig. S13B, Table S15). We then used Monocle 3 to identify DEGs within cell type partitions along the primary trajectory (Tables S14 & S16). We partitioned cells along the pseudotime scale and labeled each partition based on the maximal expression of cell type marker genes (Fig. S13C). We saw the strongest signal of dysregulation in partitions containing radial glia and cells expressing upper-layer neuronal markers. In particular, we identified 2,702 DEGs in radial glia at E12.5 and 1,181 DEGs in upper-layer neurons at E17.5, which are respectively the most abundant cell types recovered at these time points (Fig. 6A, Table S16). There was a large overlap in radial glia and primary trajectory DEGs at E12.5 and a large overlap in upper-layer neurons and primary trajectory DEGs at E17.5 (Fig. S13D), suggesting that the dysregulation signal we detected in primary trajectory cells was being driven by the predominant cell types. As in our analysis of the primary trajectory, nearly all DEGs showed only small changes in expression (Fig. 6B, Table S16).

**Figure 6.**
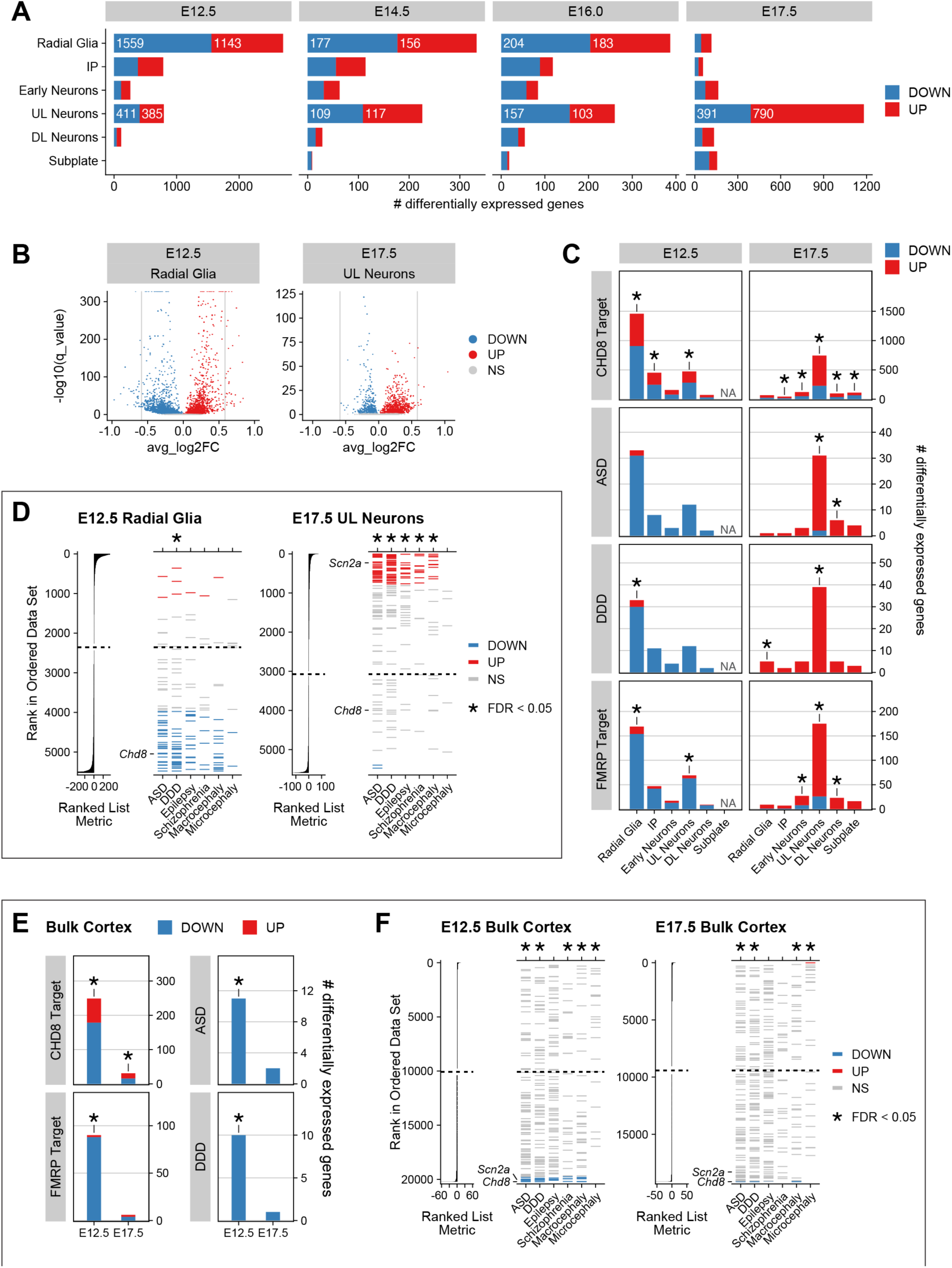
Gene sets associated with neurodevelopmental disorders are enriched among dysregulated genes in embryonic day (E) 12.5 radial glia and E17.5 upper-layer neurons. (**A**) Number of downregulated (DOWN; blue) and upregulated (UP; red) differentially expressed genes identified in cell types of the primary trajectory at each time point, determined by Monocle 3 (Methods). (**B**) Volcano plots of Monocle 3 differential expression results for radial glia at embryonic day (E) 12.5 (*left*) and upper-layer (UL) neurons at E17.5 (*right*) in the embryonic *Chd8*^+/−^ cortex, with genes color-coded by differential expression call (downregulated: DOWN, blue; upregulated: UP, red; not significantly different: NS, gray; Methods). Vertical gray lines = ±log2(1.5 fold-change). (**C**) Intersection between differentially expressed genes in each cell type at E12.5 (*left*) or E17.5 (*right*) and CHD8 target genes in the E17.5 wild type mouse cortex, autism spectrum disorder risk-associated genes (ASD), the Deciphering Developmental Disorders gene set (DDD), and FMRP target genes. Significance was determined by one-tailed Fisher exact test, with adjustment for multiple testing (Methods); * = Benjamini Hochberg (BH)-adjusted *p*-value < 0.05. (**D**) Gene set enrichment analysis (GSEA) results of differential gene expression in radial glia at E12.5 (*left*) or upper-layer neurons at E17.5 (*right*) in the embryonic *Chd8*^+/−^ cortex, assessing enrichment of neurodevelopmental disorder (NDD)-associated gene sets, including ASD genes, DDD genes, and genes associated with risk for epilepsy, schizophrenia, macrocephaly, or microcephaly, among UP or DOWN genes. For each cell type, input genes are ordered by a ranked list metric calculated from the Monocle 3 output equal to sign(avg_log2FC) * -log10(*p*-value). Genes in the ranked list that overlap each NDD-associated gene set are shown as horizontal ticks and color-coded by differential expression calls determined by Monocle 3 (red = UP; blue = DOWN; gray = NS; Methods). Dotted line indicates the zero-cross rank separating positive and negative values. * = FDR < 0.05. (**E**) Intersection between differentially expressed genes in the bulk RNA-seq embryonic mouse cortex dataset at E12.5 or E17.5 and CHD8 target genes, ASD risk-associated genes, DDD genes, and FMRP target genes. Significance was determined by one-tailed Fisher exact test, with adjustment for multiple testing (Methods); * = BH-adjusted *p*-value < 0.05. (**F**) GSEA results of differential gene expression in the bulk E12.5 (*left*) or bulk E17.5 (*right*) *Chd8*^+/−^ cortex, assessing enrichment of NDD-associated gene sets. For each time point, input genes are ordered by a ranked list metric calculated from the DESeq2 output equal to sign(avg_log2FC) * -log10(*p*-value). Genes in the ranked list that overlap each NDD gene set as horizontal ticks and color-coded by differential expression calls determined by DESeq2 (red = UP; blue = DOWN; gray = NS; Methods). Dotted line indicates the zero-cross rank separating positive and negative values. * = FDR < 0.05. See also Figures S13-S15, Table S9, and Tables S16-23. IP = intermediate progenitor; DL = deep-layer.

We then tested whether DEGs were enriched for genes whose promoters were bound by CHD8 in the E17.5 wild type mouse cortex (Tables S9 & S17). Out of 4,349 CHD8 target genes captured in our dataset, 1,621 and 1,070 genes were respectively called as differentially expressed in E12.5 and E17.5 primary trajectory cells (Fig. S14A, Table S17). We found that CHD8 target genes were significantly enriched among DEGs at E12.5 and E17.5 in primary trajectory cells and in E12.5 radial glia and E17.5 upper-layer neurons (one-tailed Fisher exact test; BH-adjusted *p*-value < 0.05) (Fig. 6C, Fig. S14A, Table S17). CHD8 target genes were also enriched among DEGs in E14.5 primary trajectory cells and other cell types including intermediate progenitors, E17.5 early neurons, deep-layer neurons, and subplate neurons, but not enriched in any cell types at E16.0 (Fig. S14A-B, Table S17).

We also sought to determine whether DEGs were enriched for genes associated with neurodevelopmental disorders. In addition to ASD risk-associated genes, we tested whether DEGs were enriched for the DDD gene set and genes associated with risk for epilepsy, schizophrenia, macrocephaly, and microcephaly (Tables S9 & S17)^2,6,54–56^. ASD risk-associated genes and DDD genes were significantly enriched among DEGs in both E12.5 and E17.5 primary trajectory cells (one-tailed Fisher exact test; BH-adjusted *p*-value < 0.05) (Fig. S14C, Table S17). DDD genes were also enriched among DEGs in E12.5 radial glia and E17.5 upper-layer neurons (Fig. 6C, Table S17). Macrocephaly risk-associated genes were enriched among DEGs in E12.5 radial glia, while ASD and schizophrenia risk-associated genes were enriched among DEGs in E17.5 upper-layer neurons (Fig. S14D, Table S17). No gene sets associated with neurodevelopmental disorders were enriched among DEGs at E14.5 or E16.0 (Fig. S14C-D, Table S17).

Finally, we considered a set of genes whose transcripts are bound by FMRP (Table S9)^57^. Loss of FMRP expression causes fragile X syndrome, which is associated with an increased risk of ASD^58,59^. FMRP target genes were enriched among E12.5 and E17.5 primary trajectory DEGs (Fig. S14C, Table S17), as well as among DEGs identified in E12.5 radial glia and E17.5 early neurons, upper-layer neurons, and deep-layer neurons (Fig. 6C, Table S17). FMRP targets were not enriched among DEGs at E14.5 or E16.0 (Fig. S14C-D, Table S17). Collectively, these results suggest that within certain cell types, loss of *Chd8* expression leads to dysregulation of CHD8 target genes, genes associated with risk for neurodevelopmental disorders, and genes potentially subject to post-transcriptional regulation by FMRP.

We then used Gene Set Enrichment Analysis (GSEA) to determine if genes associated with neurodevelopmental disorders were specifically overrepresented among genes upregulated or downregulated in the *Chd8^+/−^*background (Methods)^60,61^. GSEA considers all genes that were tested for differential expression and takes into account both significance (i.e., the Monocle *p*-value) and expression change directionality. Using GSEA, we found that ASD risk-associated genes, DDD genes, and macrocephaly risk-associated genes were enriched among downregulated genes in E12.5 primary trajectory cells (Table S18). DDD genes were also enriched among downregulated genes in radial glia at E12.5 (Fig. 6D, Table S18), while ASD and macrocephaly risk-associated genes were enriched among downregulated genes in E12.5 deep-layer neurons (Table S18). Conversely, ASD risk-associated genes, DDD genes, and epilepsy, schizophrenia, and macrocephaly risk-associated genes were all significantly enriched among upregulated genes in E17.5 primary trajectory cells, radial glia, intermediate progenitors, and neuronal subtypes (i.e., early, upper-layer, deep-layer, and subplate neurons) (Fig. 6D, Table S18). Microcephaly risk-associated genes were enriched among upregulated genes in E14.5 deep-layer neurons and E17.5 radial glia (Table S18).

To validate the trends that emerged from our single-cell differential expression and gene set enrichment analyses, we performed bulk RNA-sequencing of E12.5 and E17.5 wild type and *Chd8^+/−^* cortex (Table S19; Methods). Using DESeq2, we identified 682 DEGs in the bulk cortex at E12.5 (462 down- and 220 up-regulated) and 80 DEGs in the bulk cortex at E17.5 (37 down- and 43 up-regulated) (Table S19). We then used a one-tailed Fisher exact test and GSEA as described above to identify gene sets enriched in these DEGs (Methods). Consistent with the results obtained from the E12.5 single-cell dataset, CHD8 targets and FMRP targets were both enriched among DEGs identified in the E12.5 bulk RNA-seq dataset (Fig. 6E, Table S20). Genes downregulated in the E12.5 *Chd8^+/−^* bulk cortex were also enriched for DDD genes and ASD, schizophrenia, and macrocephaly risk-associated genes, while upregulated genes were enriched for microcephaly risk-associated genes (Fig. 6F, Table S21). In contrast, the enrichment patterns we observed in the E17.5 single-cell dataset were not supported by the E17.5 bulk RNA-seq analysis. We did not detect consistent enrichment of risk-associated gene sets among upregulated genes in the E17.5 *Chd8^+/−^* bulk cortex (Table S21). Using GSEA, only microcephaly risk-associated genes were enriched among upregulated genes at E17.5, while DDD, ASD and macrocephaly risk-associated genes were enriched among downregulated genes (Fig. 6F, Table S21), similar to GSEA results in the E12.5 bulk RNA-seq dataset.

To determine if genes dysregulated in the *Chd8^+/−^* cortex converge on particular biological functions, we tested for enrichment of GO:BP terms among DEGs identified in our scRNA-seq and bulk RNA-seq datasets (Tables S22-S23; Methods). Genes downregulated in the E12.5 primary trajectory and genes downregulated in E12.5 radial glia were both enriched for GO:BP terms such as “cell communication,” “cell migration,” “chromatin remodeling,” and “neuron projection development” (Fig. S15A-B, Table S22). Terms specifically enriched among genes downregulated in E12.5 radial glia, but not among DEGs identified in the E12.5 primary trajectory, included “cell motility” and “cell population proliferation” (Fig. S15A, Table S22). Genes upregulated across multiple cell types at E12.5 were enriched for processes associated with mitochondrial gene expression and oxidative phosphorylation (Fig. S15C, Table S22). Many of the GO:BP terms enriched among downregulated genes identified in E12.5 single-cell datasets were also enriched among downregulated genes identified in E12.5 bulk cortex (Fig. S15A-B, Tables S22-S23). In contrast, we did not detect a similar enrichment of GO:BP terms between upregulated genes identified in E12.5 single-cell datasets and DEGs identified in E12.5 bulk cortex (Fig. S15C). Although upregulated genes detected in the E17.5 primary trajectory or upper-layer neurons were enriched for GO:BP terms including “chromatin remodeling” and “regulation of cell development”, no GO:BP terms were found to be enriched in the 80 DEGs identified in the E17.5 bulk cortex (Fig. 15B-C, Tables S22-S23).

### Synaptic genes are dysregulated across excitatory neuron subtypes in the juvenile Chd8^+/−^ cortex

In light of the findings that NDD risk-associated genes, CHD8 target genes, and FMRP target genes were potentially dysregulated in upper-layer neurons at E17.5, we examined the impact of *Chd8* loss of function later in cortical development by snRNA-seq of the P25 wild type and *Chd8^+/−^* cortex (Fig. 7A, Methods). Summary data for each of the eight litter- and sex-matched samples, including total nuclei captured, mean reads per nucleus, and the number of nuclei that passed our filtering criteria (Methods, Fig. S16-S19) are presented in Table S24. Marker gene-based cell type annotation revealed that the majority of nuclei (59.1%) were from cortical glutamatergic neurons (Fig. 7B, Fig. S18A & S20, Tables S25-S27). These excitatory neurons spanned all cortical layers (L2 through L6) and projection subtypes, such as intratelencephalic projecting neurons (IT), pyramidal tract neurons (PT), near-projecting neurons (NP), and corticothalamic projecting neurons (CT) (Fig. 7B, Fig. S18B & S20, Tables S26-S27)^62–70^. Cortical inhibitory neuron (iN) subtypes were also well-represented in this dataset, as were the major glial populations of the juvenile cortex, including astroglia (AG), oligodendrocyte precursor cells (OPCs), mature and immature oligodendrocytes (OL_1 and OL_2, respectively), and microglia (MG) (Fig. 7B, Fig. S18 & S20A, Tables S26-S27)^62–66,71–73^. As observed in the embryonic scRNA-seq cortical data, we also identified a cluster of vasculature-associated (Vasc) cells in the juvenile snRNA-seq dataset (Fig. 7B, Fig. 18 & S20A, Tables S26-S27)^64,66,74^. Similar to our findings in the embryonic *Chd8^+/−^* cortex, we did not observe a significant difference in cell type representation between the juvenile wild type and *Chd8*^+/-^ cortex (two-tailed Welch’s *t*-test, Methods; Fig. S20B, Tables S27-S28).

**Figure 7.**
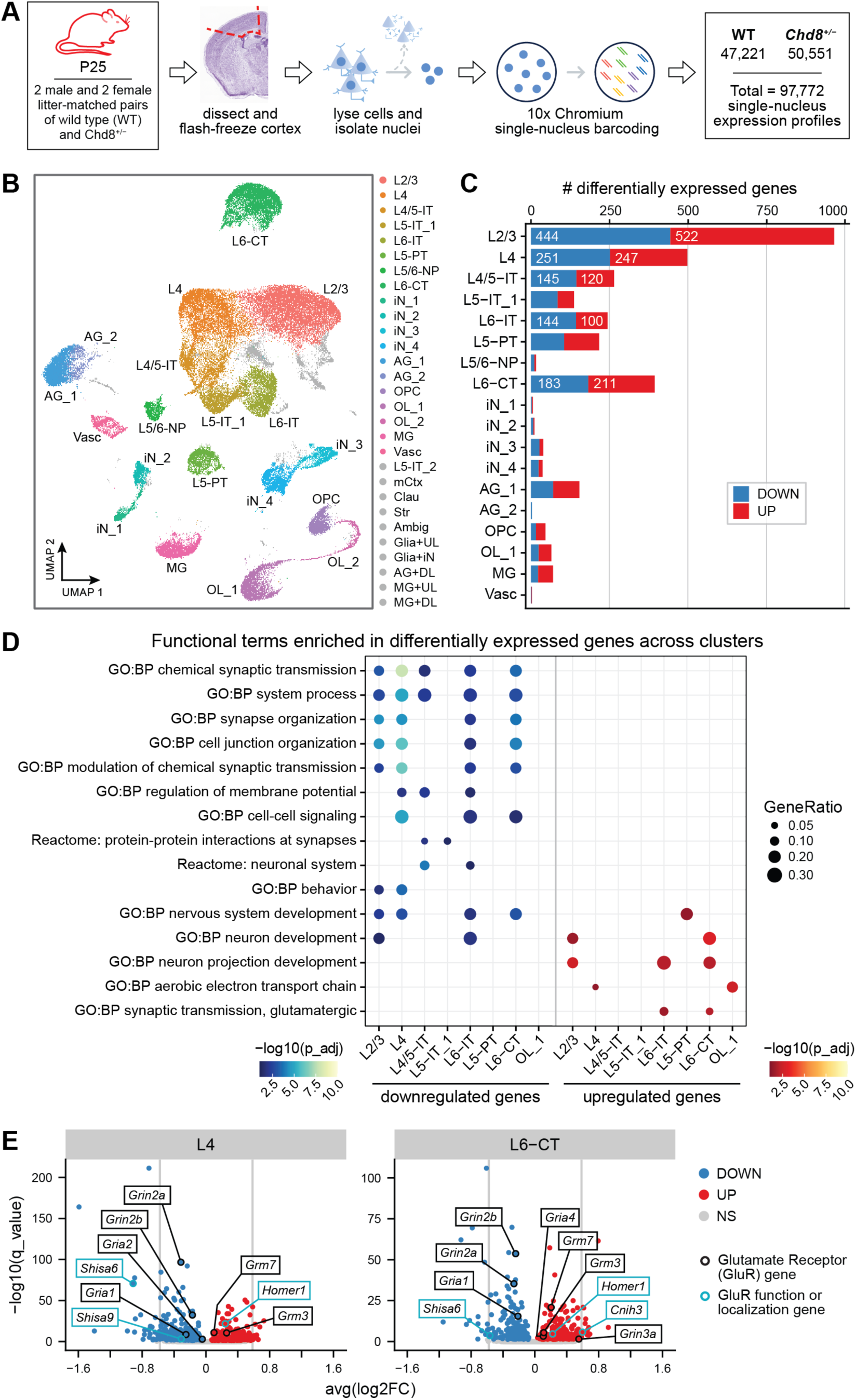
Single-nucleus RNA-sequencing (snRNA-seq) of the juvenile wild type and *Chd8*^+/−^ cortex reveals dysregulation of synaptic genes across excitatory neuron subtypes. (**A**) Schematic of experimental design, showing the dissection schema and number of nuclei collected for each genotype. The representative coronal section of the juvenile mouse brain is from the Allen Mouse Brain Atlas, Nissl-stained (postnatal day 28, position 295; Allen Institute for Brain Science (2004), developingmouse.brain-map.org). (**B**) UMAP embedding of 66,840 singlet nuclei colored by cell type assignment, with clusters excluded from downstream analyses colored in gray. Nuclei clusters were identified by a graph-based clustering approach that utilizes the Louvain algorithm (Methods). (**C**) Number of downregulated (DOWN; blue) and upregulated (UP; red) differentially expressed genes per cluster, identified by Monocle 3 (Methods). Only clusters with at least 1 differentially expressed gene are represented. (**D**) Dot plot showing enrichment of representative functional terms among downregulated and upregulated differentially expressed genes per cluster, restricted to terms enriched across multiple clusters and identified by g:Profiler and Revigo (Methods). Only clusters enriching for functional terms shared across multiple clusters are shown. Dot size corresponds to the ratio of DOWN/UP genes intersecting with the denoted functional term out of all cluster-specific DOWN/UP genes submitted as a query to g:Profiler (GeneRatio; Methods). p_adj = gSCS-adjusted *p*-value (Methods). (**E**) Volcano plots of the Monocle 3 differential expression results for L4 (*left*) and L6-CT (*right*) clusters, with a subset of genes encoding glutamate receptors (GluRs) or regulators of GluR localization and/or function outlined in black or light blue, respectively (Methods). DOWN = significantly downregulated (blue fill); UP = significantly upregulated (red fill); NS = not significantly different (gray fill); vertical gray lines = ±log2(1.5 fold-change). See also Fig. S16-S21 & S23-S25 and Tables S25-S30 & S33-S34. L = layer of excitatory cortical neuron; IT = intratelencephalic; PT = pyramidal tract; NP = near-projecting; CT = corticothalamic; iN = inhibitory neuron; AG = astroglia; OL = oligodendrocyte; OPC = oligodendrocyte precursor cell; MG = microglia; Vasc = vasculature; mCtx = medial cortex; Clau = claustrum; Str = striatum; Ambig = ambiguous cluster; UL = upper-layer cortical neuron; DL = deep-layer cortical neuron.

We then identified DEGs in all cell types using Monocle 3 (BH-adjusted *p*-value < 0.05; Methods). We detected the largest number of DEGs in upper-layer neurons (L2/3 and L4; Fig. 7C). We also observed substantial numbers of DEGs in deep-layer neuronal clusters (L5 and L6), particularly in L6-CT neurons (Fig. 7C). Comparatively, we identified fewer DEGs in the inhibitory neuron and glial populations (Fig. 7C, Fig. S21A, Tables S29-S30). Using the same approach as described above, we found that ASD risk-associated genes were enriched among DEGs in L4/5-IT neurons and L5-PT neurons (one-tailed Fisher exact test, BH-corrected *p*-value < 0.05; Figure S21B and Table S31; Methods). However, we did not observe significant enrichments in upper-layer neurons or any other cell type, in contrast to the results obtained in the single-cell E17.5 dataset (Fig. S21B, Table S31). No other NDD risk-associated gene sets were enriched in any cell type. FMRP target genes were enriched among DEGs in multiple deep- and upper-layer neuronal cell types and in a subset of interneurons (Fig. S21B, Table S31). GSEA revealed enrichment of ASD risk-associated genes in the downregulated DEGs from L4/5-IT and L6-IT neurons, but not in upregulated DEGs or in DEGs detected in other cell types (Fig. S22A-B, Table S32). DDD genes were also enriched among downregulated DEGs in L6-IT neurons (Fig. S22B, Table S32).

To further understand the biological impact of these differential expression signatures, we performed GO:BP term and KEGG and Reactome pathway enrichment analyses on the downregulated and upregulated DEGs for each cell type (Methods). Downregulated DEG sets in multiple excitatory neuronal subtypes were consistently enriched for GO:BP terms relating to synaptic transmission, synaptic signaling, and synapse organization (Fig. 7D, Fig. S23, Tables S33-S34). In comparison, fewer functional terms were consistently enriched in upregulated DEGs, including the GO:BP terms “neuron projection development” and “GO:BP synaptic transmission, glutamatergic” (Fig. 7D, Fig. S23, Tables S33-S34). We did not observe any functional term enrichment from inhibitory neuron subtypes or from L5/6-NP excitatory neurons (Fig. 7C, Table S33).

We next sought to determine whether the DEGs underlying the enrichment of these synaptic functional terms were themselves consistently downregulated or upregulated across multiple (≥2) excitatory neuronal clusters. We first extracted the downregulated genes underpinning the enrichment of the terms “GO:BP synaptic signaling,” the synaptic functional term with the largest gene set, and “GO:BP synapse organization” across neuronal clusters, yielding 114 downregulated synaptic term genes, 73 of which (64%) were downregulated in two or more neuronal subtypes (Fig. S23-S24, Table S33). We similarly extracted all upregulated genes from the neuronal clusters that were enriched for the term “GO:BP synaptic transmission, glutamatergic,” yielding 15 upregulated synaptic term genes, 11 of which (73.3%) were upregulated in two or more neuronal subtypes (Fig. S23-S24, Table S33).

Many of the consistently downregulated or upregulated synaptic term genes encoded for glutamate receptors (GluRs) (Fig. 7E, Fig. S24-S25 ). Consistently downregulated GluR genes included multiple NMDA and AMPA GluR subunit genes, two of which were downregulated in all excitatory neuronal subtypes (*Grin2a*, *Grin2b*; Fig. 7E, Fig. S24-S25, Tables S30 & S33)^75,76^. In contrast, we observed increased expression of a distinct set of ionotropic GluR subunit and metabotropic GluR genes across multiple neuronal subtypes (Fig. 7E, Fig. S24-S25, Tables S30 & S33)^75–78^. Ionotropic GluR adaptor and trafficking genes, such as *Shisa6*, *Shisa9*, *Cacng2* (TARPγ-2), *Neto1*, and *Neto2*, were also consistently downregulated, whereas we observed consistent upregulation of *Homer1*, a regulator of metabotropic GluR localization and function (Fig. 7E, Fig. S24-S25, Tables S30 & S33)^79,80^. The consistently downregulated gene set also included synaptogenesis genes and genes encoding postsynaptic scaffolding proteins, such as *Dlgap2*, *Dlg2* (PSD-93), and *Shank2* (Fig. 7E, Fig. S24-S25, Tables S30 & S33)^80–85^. In all, these consistent trends of down- and up-regulation of these genes across excitatory neuronal subtypes are suggestive of shifts in ionotropic GluR subunit composition, function, and localization; enhanced metabotropic GluR signaling; and impaired synaptogenesis in the *Chd8*^+/−^ postnatal cortex.

To assess the robustness of these trends that emerged from our single-nucleus differential expression analysis using an orthogonal approach, we performed bulk RNA-seq of the P25 wild type and *Chd8^+/−^* cortex (Methods; Table S19). We identified only 39 DEGs in the P25 bulk RNA-seq data, potentially due to the high degree of cell type heterogeneity at this time point (Methods; Fig. 7B, Table S35). These DEGs were not enriched in NDD risk-associated genes, CHD8 target genes, FMRP target genes, GO:BP functional terms, or KEGG/Reactome pathway terms (Tables S35-S36). However, GSEA revealed enrichment of ASD risk-associated genes and DDD genes among downregulated genes in these data, consistent with the snRNA-seq findings (Fig. S22C, Table S37). We also observed enrichment of macrocephaly risk-associated genes among downregulated genes, alongside enrichment of schizophrenia risk-associated genes among upregulated genes, in the P25 bulk RNA-seq dataset (Fig. S22C, Table S37).

## Discussion

In this study, we conducted a longitudinal analysis of gene expression in wild type and *Chd8^+/−^* embryonic and juvenile cortex at single-cell resolution. Our results provide insight both into the spatial and cell type-specific transcriptional trajectories of genes associated with risk for ASD and other neurodevelopmental disorders, as well as the impact of *Chd8* loss of function in particular cell types and developmental stages. We found that *Chd8* exhibits a gradient of expression in the developing mouse cortex, with both transcript and protein expression increasing from the ventricular zone to the cortical plate. This suggests that loss of *Chd8* dosage may perturb both neural progenitors and excitatory neurons, potentially with distinct biological effects. Using a metagene analysis, we then identified sets of genes associated with risk for neurodevelopmental disorders that exhibited similar expression gradients across the developing cortex, including genes that showed convergent expression with *Chd8*. Consistent with previous studies, these findings provide further evidence that ASD risk-associated genes participate in common regulatory networks and biological pathways that are disrupted in ASD^2,9,10,18,19^. Our observation that CHD8 target genes are also significantly enriched in metagenes that include *Chd8*, and our differential expression results, support that *Chd8* itself contributes to the convergent regulation of these gene sets.

The metagene approach we employed here differs from other approaches used to identify sets of genes that show convergent gene expression patterns in specific cell types, such as clustering, and is suited for analysis of single-cell data generated across developmental time series, which remains a challenge in the field. After integrating scRNA-seq datasets and identifying cells across all embryonic time points that shared a specific developmental lineage (i.e., the generation of excitatory cortical neurons, or the primary trajectory of differentiation), we defined metagenes explicitly based on the ordering of these cells along pseudotime—a useful proxy for marking the progression of cells through a developmental process^47,49,53^. Compared to clustering, binning cells along pseudotime better reflects the continuous structure of the data, provides more control over the granularity of cell type and subpopulation assignment used for downstream analyses, and enables the comparison of transient cell states, while also addressing the inherent problem of sparsity in single-cell data by grouping together transcriptionally similar cells. For comparison of gene expression patterns, calculating average expression along pseudotime is an intuitive and straightforward way to both characterize and visualize gene expression along a developmental lineage, facilitating interpretations that are contextualized within a developmentally relevant trajectory inferred from the data. Moreover, this approach enables the use of data mining tools originally designed for “true” time series data in our single-cell analysis pipeline. Metagenes thus capture spatial, temporal, cell type-specific, and cell state-specific gene expression patterns in single-cell datasets spanning developmental time points.

The ASD risk-associated gene trajectories we identified were conserved between the embryonic mouse and fetal human cortex. This finding supports that the etiology of ASD involves the disruption of deeply conserved gene regulatory networks. Metagenes enriched in ASD risk-associated genes in both species were also enriched for genes involved in axonogenesis, synaptic transmission, and synapse organization, pointing to conserved biological processes that may be disrupted due to damaging mutations in ASD risk-associated genes in humans and their orthologs in mouse. Analysis of synaptic and circuit function in mouse models may thus provide insight into how the corresponding processes are altered in human. As has been suggested by previous studies that combined gene co-expression data with genetic variation to enhance risk gene identification, the conserved metagenes enriched for ASD risk-associated genes we describe here may also include additional risk genes yet to be discovered in genetic surveys^3,86^.

We did not observe major differences in cell type representation between the wild type and *Chd8*^+/−^ cortex at any time points we assessed. An increase in the number of *Pax6*-expressing radial glia has been reported in a previously generated *Chd8* loss of function model^17^. An increase in the number of GABAergic interneurons has also been found in human *CHD8^+/−^*organoids^27^. Our results indicate that *Chd8* loss of function in the mouse cortex does not result in major expansion or reduction of any of the cell types we identified, supporting that changes in cellular composition may not be a major feature of ASD. This is consistent with a recent study that identified subtle changes in adult brains from persons with ASD compared to controls, primarily involving microglia and astrocytes^87^. However, we note that we may lack power to detect small but biologically significant differences due to the total number of cells and the low representation of some cell types in our dataset.

Our results reveal a complex pattern of gene expression perturbations resulting from *Chd8* loss of function. Consistent with previous studies, we observed modest changes in gene expression in the *Chd8^+/−^* cortex at all time points and cell types we investigated^14,17,20^. The most consistent effects on gene expression we observed in our embryonic time series were at E12.5, the earliest time point represented in our analysis. We found that CHD8 and FMRP target genes, and genes associated with risk for neurodevelopmental disorders, were enriched genes downregulated in radial glial cells in the *Chd8^+/−^* cortex. Overall, these findings were also supported by bulk RNA-seq analyses, which further suggested that genes associated with risk for ASD, schizophrenia, microcephaly, or macrocephaly were enriched among genes downregulated in the *Chd8^+/−^* cortex. Genes implicated in neuron projection development and chromatin remodeling were also downregulated in the E12.5 *Chd8^+/−^*cortex. Collectively, these results support that loss of *Chd8* dosage results in dysregulation of genes implicated in multiple neurodevelopmental disorders and may impair the neurogenic potential of radial glia.

The impact of *Chd8* loss of function at later embryonic time points, specifically at E17.5, is ambiguous. We observed upregulation of genes associated with risk for ASD, neurodevelopmental disorders, schizophrenia, macrocephaly, and microcephaly in *Chd8^+/−^* E17.5 upper-layer excitatory neurons. However, GSEA analyses using bulk RNA-seq from the *Chd8^+/−^* cortex found that genes associated with neurodevelopmental disorders were enriched among downregulated genes. This may be due to the cellular heterogeneity of the E17.5 cortex, which may result in cell type-specific effects of *Chd8* loss of function being obscured in analyses using bulk RNA-seq data. We note that we only identified 80 DEGs in the E17.5 bulk RNA-seq analysis, compared to 682 DEGs at E12.5, a stage at which the cortex is predominantly composed of progenitor cells. Alternatively, the apparent discrepancy may be due to a transient effect of *Chd8* loss of function at E17.5, as we did not observe similar enrichments in P25 upper-layer excitatory neurons.

Our results also support that loss of *Chd8* results in dysregulation of gene expression in multiple excitatory neuronal cell types in the juvenile mouse cortex. We found that ASD risk-associated genes were enriched among DEGs detected in L4/5-IT, L5-PT, and L6-IT neurons. This is consistent with both our finding in this study and findings from previous studies that ASD risk-associated genes show convergent expression in the excitatory neuronal lineage and supports a role for CHD8 in the regulation of other ASD risk-associated genes in cortical excitatory neurons^2,9,10^. We also found that genes involved in synaptic organization and signaling were dysregulated in multiple deep- and upper-layer excitatory neuron subtypes in the *Chd8^+/-^* cortex, a substantial number of which were consistently dysregulated across excitatory neuron subtypes. Many of these consistently dysregulated genes encoded glutamatergic receptor subunits and regulators of glutamate receptor function, or were genes implicated in synaptogenesis. Notably, most of the consistently downregulated glutamate receptor subunit genes either encode key glutamate receptor components in both developing and mature synapses or are specifically associated with mature synapses, such as *Grin2a*/GluN2a, while multiple consistently upregulated glutamate receptor genes encode subunits that are prevalent in immature neurons^75,76,88–91^. These findings suggest that loss of *Chd8* may impair glutamatergic neurotransmission and delay synaptic maturation across broad swaths of excitatory neurons in the cortex. Deficits in synaptic activity have been reported in cultured human *CHD8^+/−^* cortical neurons and cortical neurons derived from *Chd8^+/−^* mice^26^. Dysregulation of synaptic gene expression has also been reported in analyses of postmortem brain samples from ASD probands, suggesting impaired synaptic function is a feature of ASD that may be amenable to study using *Chd8^+/−^* mice and other mouse models of ASD risk-associated genes^87^.

Collectively, our results point to a complex pattern of gene expression dysregulation in the developing and juvenile cortex resulting from *Chd8* loss of function, impacting radial glia in the early cortex and excitatory neuronal subtypes postnatally. This developmental shift in the cell type predominantly affected by *Chd8* loss of function supports that *Chd8* has distinct, cell type-specific functions across cortical development and that even within a cell type, such as excitatory neurons, CHD8 function is dynamic across cellular maturation and impacts specific developmental events of cell differentiation, such as synaptogenesis and synapse maturation. Thus, the molecular mechanisms involved in each cell type across development are likely to be distinct. In dividing cells, *Chd8* has been shown to be required for expression of S-phase-specific genes and to suppress p53-mediated apoptosis, and loss of *Chd8* has been linked to defects in progenitor proliferation and differentiation^24,28,92^. Studies in cultured human neurons suggest that *CHD8* regulates target genes of the MAPK/ERK pathway, which informs synaptic plasticity^93^. Damaging mutations in *CHD8* and other ASD risk-associated genes will thus have pleiotropic effects, altering neurodevelopment via the perturbation of multiple cell types and distinct developmental processes, potentially throughout the lifespan. Understanding how loss of *CHD8* contributes to ASD etiology will therefore require a broad approach that incorporates analysis of progenitor proliferation, neurogenesis, synaptic maturation, and circuit development and function across all stages of brain development in multiple experimental models.

## Methods

### Generation of *Chd8^+/−^* mice

All animal work was performed in accordance with approved Yale IACUC protocols (#2019–11167, #2022-11167, and #2020–07271). The *Chd8^+/−^*line was generated at the Yale Genome Editing Center. Regions flanking exon 3 of *Chd8* transcript NM_20167.3 were isolated *in silico* and input into the CRISPR Design Tool (http://crispr.mit.edu/, now deprecated) to select guides with minimal off-targeting effects (determined by high Design Tool Score, minimal possible off-target sites genome-wide, and no off-target sites in chromosome 14 exonic regions). Guides with off-target binding sites in the genome with 3 or fewer mismatches were removed, as were those with targets in coding regions of genes on chromosome 14 to prevent the segregation of off-target mutations with the modified locus.

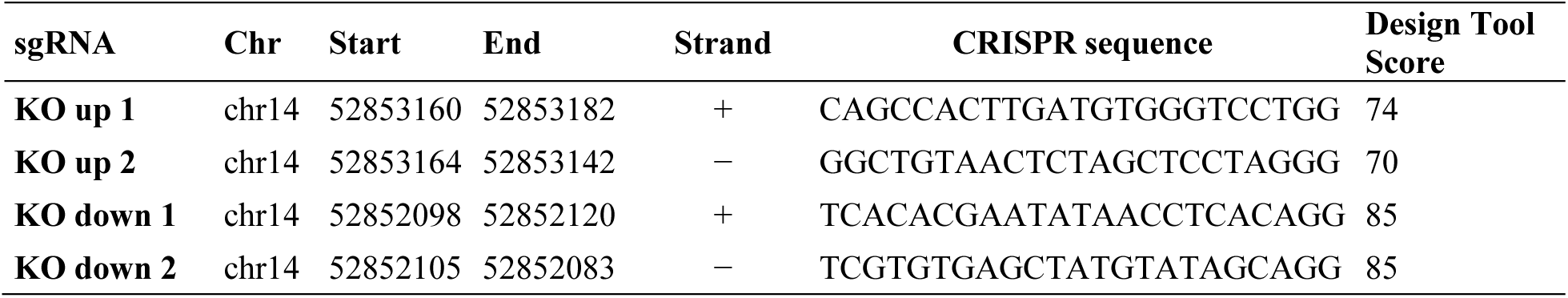

The sgRNA linker sequences were cloned into the BbsI site of plasmid pX335-U6-Chimeric_BB-CBh-hSpCas9n(D10A) (Addgene, #42335) to generate a chimeric sequence of sgRNA-trRNA downstream of the T7 promoter^94^. The plasmid containing the indicated sgRNA target sequences were individually PCR amplified and transcribed *in vitro* using the MEGAshortscript T7 kit (ThermoFisher Scientific, AM1354) and purified using the MEGAclear Transcription Clean-Up kit (ThermoFisher Scientific, AM1908). The Cas9 nickase (Cas9n) transcript was synthesized using the mMessage mMACHINE T7 ULTRA Transcription Kit (ThermoFisher Scientific, AM1345) and purified using the RNeasy Mini RNA Cleanup Kit (QIAGEN, #74104) and resuspended in embryo microinjection buffer (10 mM Tris-HCl, pH 7.4, 0.25 mM EDTA). Following confirmation of RNA purity and concentration using Agilent Bioanalyzer, sgRNAs (15 ng/μL each) and Cas9n (30 ng/μL) were combined at the indicated concentrations for injection into C57BL/6J zygotes. Following embryo transfer to super-ovulated female mice, live born pups were screened by PCR using the *Chd8* genotyping primers described in the following section. One male founder mouse was obtained with non-homologous end joining (NHEJ) of breakpoints generated by Cas9n, deleting exon 3. The *Chd8^+/−^* F0 founder was backcrossed to the wild type C57BL/6J strain (Jackson Laboratories, #000664) for over 5 generations before experimental use. Mice were maintained in a Yale Animal Resources Center (YARC) managed facility under a standard 12-hour light/dark cycle and environmental monitoring according to YARC policies and procedures.

### Genotyping

For embryonic and postnatal genotyping, tissue was lysed using the DNeasy Blood and Tissue Kit (QIAGEN, #69506). Twenty μg of genomic DNA were used as an input for the PCR reaction. PCR was performed with Q5 High Fidelity 2X Master Mix (New England Biosciences, M0492), 10 μM forward genotyping primer, 10 μM reverse genotyping primer, and nuclease-free water. For *Chd8* genotyping, PCR was performed under the following conditions: initial denaturation at 98°C for 30 seconds followed by 30 cycles at 98°C for 10 seconds, 66°C for 30 seconds, and 72°C for 1 minute, with a final extension at 72°C for 2 minutes. Forward and reverse *Chd8* genotyping primer sequences are 5’-AACAGGCTGTCTCATGGGAA-3’ and 5’-AAGCCACACTGCCTTGAAAG-3’, respectively. PCR products were resolved by 1.5% agarose gel electrophoresis and examined for expected product sizes of 1495bp for the wild type allele and 434bp for the edited allele (Fig. S3). Sex was determined by PCR using the primers 5’-GATGATTTGAGTGGAAATGTGAGGTA-3’ and 5’-CTTATGTTTATAGGCATGCACCATGTA-3’, as previously described^95^.

### Animal breeding and tissue preparation for immunohistochemistry (IHC) and imaging

Embryos were collected from timed pregnancies at E12.5, E14.5, E16.0, and E17.5 (E0.5 = vaginal plug date). Embryonic brains were dissected, immersion-fixed in 4% paraformaldehyde for 18-24 hours, and cryoprotected sequentially in 15% and 30% sucrose solution for 24 hours and 48-72 hours, respectively. The brains were frozen in Tissue-Tek OCT Compound (Electron Microscopy Sciences, #62550) on a dry ice-ethanol slurry, stored at -80°C, and cryosectioned into 30μm coronal or sagittal sections (Leica Biosystems, CM3050 S). For IHC, sections were hydrated in PBS, permeabilized in 0.3% Triton X-100 in PBS, and blocked in normal donkey serum (NDS; PBS with 5% NDS and 0.3% Triton X-100) for 15 minutes each. Sections were then incubated in primary antibody solution (PBS with 5%-7.5% NDS and 0.3% Triton X-100) in a humid chamber at room temperature for 23 hours. Following three 10-minute washes with PBS, sections were incubated with one of two Cy3 fluorophore-conjugated secondary antibodies raised in donkey hosts (Jackson ImmunoResearch; anti-rabbit: #711-165-152; anti-chicken: #703-165-155; 1:300) and Hoechst 33342 (ThermoFisher Scientific; 1:300) at room temperature, then washed with PBS (3 x 10 minutes). Sections were mounted with ProLong Gold Antifade Mountant (ThermoFisher Scientific, P10144). The following primary antibodies were used: CHD8 C-terminus (Abcam, ab84527, rabbit, 1:1,000 for E12.5, E14.5, E16.0; 1:600-1:750 for E17.5), POGZ (Abcam, ab167408, rabbit, 1:750), TBR1 (EMD Millipore, AB2261, chicken, 1:500-1:750), and PAX6 (EMD Millipore, AB2237, rabbit, 1:750). Immunostained specimens were imaged on a Carl Zeiss AxioCam MRm coupled to an Axioimager Z2 epifluorescence microscope equipped with the ApoTome2 imaging system (Carl Zeiss Microimaging). Image processing was performed using Carl Zeiss Axiovision and Carl Zeiss Zen LE software.

### Western blotting

Tissue extracts were obtained by lysing E16.0 mouse cortical samples in radioimmunoprecipitation assay (RIPA) buffer supplemented with a protease inhibitor solution (cOmplete, EDTA-free Protease Inhibitor Cocktail; Roche, #11873580001) and 100mM PMSF at 4°C for 2 hours. The lysates were then cleared by centrifugation at 16,000 x g, and the total protein concentration was determined with the Pierce BCA Protein Assay Kit (ThermoFisher Scientific, #23225). Lysates were mixed with Laemmli sample buffer, and equal amounts of protein samples were resolved on a 7.5% SDS-PAGE gel (7.5% Mini-PROTEAN TGX Precast Protein Gels, 10-well; Bio-Rad, #4561023). The proteins were transferred onto a PVDF membrane and blocked in 5% nonfat dry milk (NFDM) in TBS with 0.1% Tween-20 (TBS-T). For each primary antibody, the membrane was incubated overnight at 4°C, washed with TBS-T after each primary antibody incubation, and incubated with secondary antibodies conjugated to horseradish peroxidase for one hour at room temperature.

Primary antibodies used for immunodetection included anti-CHD8 (Abcam, ab114126, rabbit), diluted 1:1,000 in TBS-T with 5% BSA, and anti-actin (Abcam, ab8226, mouse), diluted 1:1,000 in TBS-T with 5% NFDM. Horseradish peroxidase-conjugated secondary antibodies used included donkey anti-rabbit secondary antibody (Cytiva, NA934) and sheep anti-mouse secondary antibody (Cytiva, NA931), both diluted 1:10,000 in TBS-T with 5% NFDM. Membranes were visualized using SuperSignal West Femto Maximum Sensitivity Substrate (ThermoFisher Scientific, #34096). Western blot quantification was performed on the scanned films based on densitometry measured in ImageJ. Actin was used as a loading control and for by-replicate normalization. Statistical significance was then determined by one-tailed Welch’s *t*-test, performed on the actin-normalized CHD8 signal (Fig. 1B, Table S1). For sex- and film-matched wild type and *Chd8*^+/-^ comparisons, these actin-normalized CHD8 signal values were then divided by the actin-normalized CHD8 signal in the corresponding sex- and film-matched wild type control (Table S1).

### Sample and library preparation for single-cell RNA-sequencing (scRNA-seq)

Wild type dams impregnated by *Chd8*^+/−^ males were euthanized at the indicated embryonic stage and the cortices were micro-dissected in ice-cold Hibernate EB media (Brain Bits, HEB). Medial structures of the dorsal telencephalon were excluded to avoid capturing the hippocampal primordium. After PCR confirmation of genotype and sex, sex- and litter-matched pairs of wild type and *Chd8*^+/−^ cortices were selected for single-cell capture and sequencing. Four litter-matched sample pairs, two male and two female, were processed from each stage (Figure S3, Table S2). We required samples to exhibit cell viability >85% prior to cell capture, determined using the Countess II automated cell counter (ThermoFisher Scientific, AMQAX1000) and Trypan Blue staining, per the manufacturer’s guidelines (ThermoFisher Scientific, T10282; Table S2). Cell preparations were loaded onto the 10x Genomics Chromium Controller (PN-120270) for single-cell encapsulation and barcoding, followed by library generation and sample indexing in accordance with the 10x Genomics Chromium Next GEM Single-Cell 3’ Reagent Kits for v2 chemistry (PN-120236, PN-120237, PN-120262)^96^. Barcoded scRNA-seq libraries were sequenced on an Illumina HiSeq2500, with 26 cycles for read 1 and 98 cycles for read 2. Single-cell capture, library preparation, and sequencing were performed by the Yale Center for Genome Analysis.

### Sample and library preparation for single-nucleus RNA-sequencing (snRNA-seq)

Postnatal day (P) 25 wild type and *Chd8* heterozygous, sex-matched littermates were euthanized by cervical dislocation followed by decapitation (Table S24). We performed cortical dissections in ice-cold Hibernate AB complete media (BrainBits, HAB). For isolating cortical structures, each brain was separated by hemisphere, followed by removal of subcerebral structures and the hippocampus. To match the cortical tissue collection strategy used for embryonic dissections, we discarded the medial cortex, then collected the remaining cortex by cutting along the medial edge of the corpus callosum, avoiding the striatum. Tissue samples were blotted dry, minced finely, then flash-frozen and stored in liquid nitrogen until nuclei isolation. Four litter-matched sample pairs (two male and two female) were processed across two batches, with each batch corresponding to two litter-matched sample pairs, one male and one female. Nuclei isolation was performed using the 10x Genomics Nuclei Isolation Kit with RNase Inhibitor (PN-1000494) in accordance with the manufacturer’s instructions. Nuclei integrity, debris contamination, and quantification were assessed using ReadyCount Green/Red Viability Stain (acridine orange/propidium iodide; ThermoFisher Scientific, A49905) and either a hemocytometer or fluorescent automated cell counter (DeNovix, CellDrop FLi). Nuclei preparations were loaded onto the 10x Genomics Chromium Controller (PN-120270) for single-nucleus encapsulation and barcoding, followed by library generation and sample indexing in accordance with the 10x Genomics Chromium Next GEM Single-Cell 3’ Reagent Kits for v3.1 chemistry (PN-1000120, PN-1000121, PN-1000213). Barcoded snRNA-seq libraries from the same batch were pooled into a single lane and sequenced on the Illumina NovaSeq 6000 platform, with 28 cycles for read 1 and 90 cycles for read 2. Single-nucleus capture, library preparation, and sequencing were performed by the Yale Center for Genome Analysis.

### Sample and library preparation for bulk RNA-sequencing

We employed the same dissection strategies for collection of (i) the developing cortex from E12.5 and E17.5 *Chd8*^+/−^ and wild type embryos and (ii) the juvenile cortex from P25 *Chd8*^+/−^ and wild type mice as performed for scRNA-seq and snRNA-seq, respectively. Six litter-matched sample pairs (three male and three female) were processed for each stage, which included microdissection of the cortex in ice-cold PBS and flash-freezing in liquid nitrogen until verification of sex and genotype (Table S19). For each sample, RNA was extracted with the miRNeasy Micro Kit with on-column DNase digestion (QIAGEN, #217084 and #79254; for E12.5 samples) or the miRNeasy Mini Kit with on-column DNase treatment (QIAGEN, #217004 and #79254; for E17.5 and P25 samples). Sample quality was assessed by Agilent Bioanalyzer to determine the sample’s RNA integrity number (RIN), requiring a RIN ≥ 9.0 to proceed to library preparation (Table S19). PolyA-enriched cDNA library preparation, using the KAPA mRNA HyperPrep Kit (Roche, #08098123702), and paired-end sequencing (2 x 100bp) on the Illumina NovaSeq 6000 platform were performed by the Yale Center for Genome Analysis.

### Single-Cell and Single-Nucleus Analysis

#### Single-cell and single-nucleus RNA-sequencing read alignment

Reads from all single-cell and single-nucleus RNA-sequencing experiments were aligned to the mm10 reference genome using STAR v2.7.9a (GENCODE vM23/Ensembl 98, cellranger reference 2020-A; parameters: --outSAMtype BAM Unsorted --outSAMattributes NH HI AS nM CR CY UR UY GX GN --outSAMprimaryFlag AllBestScore --outSAMmultNmax 10 --outBAMcompression 10)^97^. Genome indices were generated using the recommended settings (--sjdbOverhang 99 --genomeSAsparseD 3). By default, any reads that map to more than 10 loci were discarded. In order to mitigate the effect of read alignments to expressed pseudogenes which could inflate gene expression counts, we also discarded multi-mapped reads with more than one best scoring alignment using SAMtools and a custom shell script available at the GitHub repository associated with this paper^98^.

For scRNA-seq data, STARsolo was used to generate cell-by-gene count matrices and select cells for downstream analysis (implemented in STAR v2.7.9a; parameters: --soloType CB_UMI_Simple–-soloCBwhitelist 737K-august-2016.txt–-soloBarcodeReadLength 1 --soloCBmatchWLtype 1MM_multi–-soloInputSAMattrBarcodeSeq CR UR --soloInputSAMattrBarcodeQual CY UY--soloMultiMappers Unique --soloUMIdedup 1MM_CR --soloUMIfiltering ---soloCellFilter CellRanger2.2 3000 0.99 10). For snRNA-seq data, count matrices were generated using STARsolo with modifications recommended for single-nucleus data and 10x Genomics single-cell/nucleus 3’ libraries generated with v3.1 chemistry. Specifically, to capture both pre-mRNA and spliced mRNA transcripts, calculation of per-cell gene counts utilized reads aligning anywhere within the full gene body, including introns, exons, and exon junctions (--soloFeatures GeneFull)^97,99^. UMI length was also specified based on the read structure of libraries generated using 10x Genomics Chromium Next GEM Single-Cell 3’ Reagent Kits for v3.1 chemistry (--soloUMIlen 12). Nuclei filtering was performed using the EmptyDrops algorithm with default settings (--soloCellFilter EmptyDrops_CR 3000 0.99 10 45000 90000 500 0.01 20000 0.01 10000)^100^.

#### Quality control and cell filtering

For quality control of scRNA-seq data, we inspected the total feature counts per cell (nCount_RNA), the total genes detected per cell (nFeature_RNA), and the percentage of counts originating from mitochondrial RNA (percent_mito). Discarding multi-mapped reads in the previous step greatly reduced the relative contribution of mitochondrial RNA. We also observed higher percentages of mitochondrial content in samples collected at later time points compared to earlier time points. Therefore, based on the per-sample distributions of these quality metrics, we removed cells with >2% mitochondrial content in E12.5 and E14.5 samples and cells with >3% mitochondrial content in E16.0 and E17.5 samples (Fig. S4A). We also removed cells with fewer than 500 genes (nFeature_RNA < 500) and cells with total feature counts greater than 4 standard deviations above the per-sample mean (nCount_RNA.scale > 4; Fig. S4B). For quality control of snRNA-seq data, we inspected the percentage of counts originating from mitochondrial RNA per nucleus and removed nuclei with >1% mitochondrial content (Fig. S16A). Given that cortical neurons and glia have markedly different total counts and numbers of detected genes per nucleus (Fig. S16D), we did not employ these metrics for quality control of the snRNA-seq dataset^101^.

#### Normalization and regressing out cell cycle differences

Seurat v4.0.4 was used for preliminary normalization and single-cell analysis^102^. Counts were log-normalized using the “NormalizeData” function (default parameters: normalization.method = “LogNormalize”, scale.factor = 10000). For scRNA-seq data, cell cycle scores were assigned to each cell based on the expression of G2/M and S phase markers using the Seurat function “CellCycleScoring” (ctrl = NULL)^103^. Human gene symbols were converted to mouse using biomaRt v2.50.3^104^. Samples were grouped by batch (i.e., samples processed on the same cell-barcoding chip), and then each batch was normalized using SCTransform (ncells = 5000, residual.features = NULL, variable.features.n = 3000, vars.to.regress = c(“CC.Difference”, “percent_mito”), do.scale = F, do.center = T, clip.range = c(-sqrt(x = ncol(x = object[[assay]])/30), sqrt(x = ncol(x = object[[assay]])/30)), return.only.var.genes = T, seed.use = 1448145)^105^. The “SCTransform” function was also used to regress out the signal from percent mitochondrial content (percent_mito) for single-cell and single-nucleus RNA-sequencing data, along with the difference between G2M and S phase scores (CC.Difference) for scRNA-seq data.

#### Integration and UMAP embedding

We performed integration using the following Seurat functions with default parameters for datasets that have been normalized with SCTransform: “SelectIntegrationFeatures” (nfeatures = 2000), “PrepSCTIntegration” (sct.clip.range = NULL), “FindIntegrationAnchors” (scale = T, normalization.method = “SCT”, set.clip.range = NULL, reduction = “cca”, l2.norm = T, dims = 1:30, k.anchor = 5, k.filter = 200, k.score = 30, max.features = 200, nn.method = “annoy”, n.trees = 50, eps = 0), “IntegrateData” (normalization.method = “SCT”, features = NULL, features.to.integrate = NULL, dims = 1:30, k.weight = 100, weight.reduction = NULL, sd.weight = 1, sample.tree = NULL, preserve.order = F, eps = 0)^33^. For the scRNA-seq dataset, we used the largest batch that included both male and female samples (batch F) as the reference dataset (Table S2). Since the snRNA-seq samples were processed in two batches, we did not specify a reference dataset during integration.

We then performed principal component analysis (PCA) on the integrated datasets and embedded the first 12 PCs (for the scRNA-seq data) or first 17 PCs (for the snRNA-seq data) using uniform manifold approximation and projection, implemented in Seurat: “RunPCA” (npcs = 50, rev.pca = F, weight.by.var = T, seed.use = 42), “RunUMAP” (single-cell data: dims = 1:12, reduction = “pca”, umap.method = “uwot”, n.neighbors = 30, n.components = 2, metric = “cosine”, n.epochs = NULL, learning.rate = 1, min.dist = 0.2, spread = 1, set.op.mix.ratio = 1, local.connectivity = 1, repulsion.strength = 1, negative.sample.rate = 5, uwot.sgd = F, seed.use = 42; single-nucleus data: dims = 1:17, min.dist = 0.3, n.neighbors = 40, same parameters otherwise)^35^.

#### Cluster analysis and post-clustering filters

We also used Seurat to identify clusters of cells and nuclei. Seurat implements a graph-based clustering approach. First, a shared nearest neighbor graph was constructed in PCA space using the “FindNeighbors” function (single-cell data: reduction = “pca”, dims = 1:12, k.param = 20, prune.SNN = 1/15, nn.method = “annoy”, n.trees = 50, annoy.metric = “euclidean”, nn.eps = 0, l2.norm = F; single-nucleus data: dims = 1:17, same parameters otherwise), and then the Louvain algorithm was applied to define groups of cells using the “FindClusters” function (single-cell data: modularity.fxn = 1, resolution = 0.4, method = “matrix”, algorithm = 1, n.start = 10, n.iter = 10, random.seed = 0, group.singletons = T; single-nucleus data: resolution = 0.5, same parameters otherwise)^106^. We used the Seurat function “FindAllMarkers” to identify highly expressed genes in each cluster (single-cell data: assay = “RNA”, logfc.threshold = 0.5, test.use = “wilcox”, min.pct = 0.1, min.diff.pct = -Inf, only.pos = F, max.cells.per.ident = Inf, random.seed = 1, min.cells.feature = 3, min.cells.group = 3, mean.fxn = NULL, base = 2, return.thresh = 0.01; single-nucleus data: logfc.threshold = 0.25, same parameters otherwise). We then labeled and merged the initial clustering results (seurat_clusters) into broad cell type clusters based on their expression of canonical cell type markers (Fig. S7-S8, Table S3)^36–46^.

For the snRNA-seq dataset, we performed additional filtering steps to address doublets, clusters containing relatively few nuclei, and clusters disproportionately consisting of nuclei from one batch. We observed multiple small clusters (45 to 474 nuclei) marked by genes associated with disparate cell types, indicative of doublets within the data (Fig. S17, Table S25). To identify doublets, we employed scDblFinder v1.18.0 using default settings and cluster-informed artificial doublet generation for each sample (clusters = “seurat_clusters”, samples = “sample.id”, clustCor = NULL, artificialDoublets = NULL, knownDoublets = NULL, dbr = NULL, dbr.sd = NULL, nfeatures = 1352, dims = 20, k = NULL, removeUnidentifiable = T, includePCs = 19, propRandom = 0, propMarkers = 0, aggregateFeatures = F, returnType = “sce”, score = “xgb”, processing = “default”, metric = “logloss”, nrounds = 0.25, max_depth = 4, iter = 3, trainingFeatures = NULL, unident.th = NULL, multiSampleMode = “split”, threshold = T)^107^. This approach classified the vast majority of droplets in these small clusters as doublets (87.8% to 100% doublets; Fig. S17). After removing all droplets classified as doublets, we applied a series of cluster filtering steps before proceeding to downstream analyses. Specifically, we excluded any cluster that (i) contained <200 nuclei; (ii) could not be assigned a cell type identity, due to ambiguous marker genes (i.e., cluster “Ambig”); or (iii) disproportionately consisted of nuclei from one batch (≥75%), indicating dissection variability between batches (i.e., clusters mCtx and Str; Fig. S17C-D, Fig. S18-S19, gray clusters in Fig. 7B, Tables S26-S27).

In order to test whether loss of *Chd8* had an effect on cell type representation, for each cell type we used a two sample, two-tailed *t*-test (two-tailed Welch’s *t*-test) to compare the number of cells in wild type samples versus *Chd8*^+/−^ samples (t.test, “n_cells ∼ genotype”, alternative = “two.sided”) (Fig. S9, Table S5). The same test was performed to compare cell type representation in the single-nucleus data (Fig. S20B, Table S28).

#### PHATE and pseudotime analysis of single-cell RNA-sequencing data

We used potential of heat diffusion for affinity-based transition embedding (PHATE) to infer gene expression trajectories in our scRNA-seq datasets^47^. For this analysis, we focused on cells in the excitatory neuronal lineage (i.e., radial glia, intermediate progenitors, early neurons, upper-layer neurons, and deep-layer neurons). We embedded these cells (the first 50 PCs) using PHATE, implemented in phateR v1.0.7 (ndim = 2, knn = 80, decay = 40, n.landmark = 2000, gamma = 0, t = 80, mds.solver = “sgd”, knn.dist.method = “euclidean”, mds.method = “metric”, mds.dist.method = “euclidean”, npca = 100, seed = 1). To infer pseudotime ordering of cells, we fit a principal curve through the primary trajectory of the PHATE embedding (excluding the small, secondary trajectory of cells that branched from the primary radial glial population shown in Fig. 2D) using the “principal_curve” function implemented in princurve v2.1.6 (start = NULL, thresh = 0.001, maxit = 10, stretch = 2, smoother = “smooth_spline”, approx_points = 100)^48^. We assigned each cell a value from 0 to 1 based on its arc-length from the beginning of the curve. Correlation with *Sox2* expression was used to establish the start and end points of the trajectory.

#### Downsampling count matrices

In order to account for technical differences in count depth between scRNA-seq datasets, we used the “downsampleMatrix” function in DropletUtils v1.14.2 to downsample count matrices such that samples within each time point had the same median feature counts (prop = downsampling proportion, bycol = F; downsampling proportion was calculated by dividing the median counts per sample by the minimum median counts per time point)^108^. Downsampled counts were used for inferring gene expression trajectories and single-cell differential expression analysis. We did not downsample count matrices for snRNA-seq datasets.

#### Inferring gene expression trajectories

We defined gene expression trajectories by calculating the conditional mean along pseudotime (Fig. S10A). Male and female samples were aggregated for each time point and genotype; we then divided cells in the primary trajectory (as defined above) into 20 equally-spaced bins along the pseudotime scale and calculated average expression per bin for each gene. Gene expression trajectories were then identified as described in “Defining metagenes” below.

#### Defining metagenes

We classified gene expression trajectories into metagenes—groups of genes with convergent expression along the pseudotime scale. Prior to defining metagenes, we filtered out lowly expressed genes (i.e., genes detected in fewer than 5% of cells per bin across all pseudotime bins). E12.5 expression data were also excluded given the low representation of intermediate progenitors and neurons along the differentiation trajectory at this time point. Gene expression trajectories were then centered, scaled, and transformed using symbolic aggregate approximation (SAX) implemented in jmotif v1.1.1 (Fig. S10B; paa, paa_num = 8; series_to_chars, a_size = 9)^109,110^.

We then performed *k*-means clustering (*k* = 25) on SAX-transformed wild type gene expression trajectories using the “kmeans” function in the base R stats package (Fig. S10C; centers = 25, iter.max = 20, nstart = 1, algorithm = “Hartigan-Wong”). The resulting clusters of trajectories were further aggregated into 10 metagene centers by hierarchical clustering using correlation distance and complete linkage (Fig. S10D; *d* = 1 – *r,* where *r* is the pairwise Pearson correlation between *k*-means cluster centers; hclust, method = “complete”; cutree, h = 0.6). Metagene naming order (A through J) was determined using an optimal leaf ordering algorithm that minimizes the distance between neighboring objects (i.e., cluster centers) in the dendrogram, implemented in seriation v1.3.5 (reorder.hclust, method = “OLO”)^111^. Finally, each gene expression trajectory identified in wild type or *Chd8^+/-^*datasets was assigned to a metagene based on highest Pearson correlation between the SAX-transformed trajectory and the metagene center (Fig. S10E).

#### Metagene enrichment analysis

We performed Gene Ontology (GO) functional enrichment analysis using the g:GOSt tool in the g:Profiler web toolset (https://biit.cs.ut.ee/gprofiler/gost), implemented via the gprofiler2 interface^112,113^. To test each metagene gene set for enrichment, we used all genes assigned to that metagene across the E14.5, E16.0, and E17.5 wild type datasets. All genes, across all metagenes and time points, were used as a statistical background (domain_scope). For multiple testing correction and significance calling, we used default parameters (user_threshold = 0.05, correction_method = “g_SCS”). We then used the Revigo web tool (http://revigo.irb.hr/) to summarize the resulting lists of significant GO:BP terms for each metagene (resulting list size = Small, species = Mus musculus, semantic similarity measure = SimRel)^114^. Using the adjusted *p*-values generated by g:Profiler, we provided Revigo with values for each GO term equal to - log10(adjusted *p-*value); after grouping terms based on semantic similarity, these values were used by Revigo to select a representative “parent term” from each group. The full list of significantly enriched GO:BP terms is provided in Table S7. We also tested metagenes for enrichment for ASD risk-associated genes, genes associated with developmental and intellectual disability identified by the Deciphering Developmental Disorders (DDD) consortium, and genes whose promoters were bound by CHD8 in E17.5 mouse cortex using a one-tailed Fisher exact test in R (fisher.test, alternative = “greater”; Table S9)^2,6,18^. For multiple testing correction, we used the “p.adjust” function in the base R stats package (method = “BH”).

#### Pseudotime and metagene analysis of human cortex scRNA-seq data

Human cortex scRNA-seq data from gestational weeks 17 and 18 were generated by the Geschwind laboratory at The University of California, Los Angeles^52^. We downloaded the raw count matrix and associated metadata via the Cortical Development Expression viewer webtool (http://solo.bmap.ucla.edu/shiny/webapp/). We log-normalized the count matrix in Seurat using the “NormalizeData” function (normalization.method = “LogNormalize”, scale.factor = 10000) and identified highly variable features using the “FindVariableFeatures” function with default parameters (selection.method = “vst”, loess.span = 0.3, clip.max = “auto”, mean.function = FastExpMean, dispersion.function = FastLogVMR, num.bin = 20, binning.method = “equal_width”, nfeatures = 2000, mean.cutoff = c(0.1, 8), dispersion.cutoff = c(1, Inf)). Using variables included in the associated metadata and the “ScaleData” function, we regressed out counts per cell (Number_UMI), sample donor, library, percent mitochondrial content, and the difference between G2M and S phase scores (S_phase_score - G2M_phase_score). The resulting residuals were scaled and centered. We then performed PCA using the “RunPCA” function (npcs = 50, rev.pca = F, weight.by.var = T, seed.use = 42).

Prior to embedding with PHATE, we excluded the following cell types based on cluster labels included in the associated metadata: medial ganglionic eminence-derived interneurons (InMGE), caudal ganglionic eminence-derived interneurons (InCGE), oligodendrocyte precursor cells (OPC), endothelial cells (End), pericytes (Per), and microglia (Mic)^52^. We then embedded the remaining cells (first 40 PCs) using PHATE (knn = 40, t = 40, gamma = 0, seed = 1) and computed pseudotime along the primary trajectory by fitting a principal curve to the PHATE embedding as described above (principal_curve, start = NULL, thresh = 0.001, maxit = 10, stretch = 2, smoother = “smooth_spline”, approx_points = 100). Gene expression trajectories and metagenes were computed as described above (Fig. S10). We calculated the Pearson correlation between human metagene centers and mouse metagene centers, and then labeled human metagenes based on maximal correlation with mouse metagene centers, denoted with a prime (′) symbol. We used g:Profiler and Revigo for GO:BP enrichment analysis as described above. The full list of significantly enriched GO:BP terms are provided in Table S12. We used a one-tailed Fisher exact test to test for enrichment of ASD risk-associated genes, DDD genes, and genes whose promoters were bound by CHD8 in the human mid-fetal cortex as described above (Table S9)^2,6,18^.

#### Single-cell and single-nucleus differential expression analyses

We used Monocle 3 for single-cell and single-nucleus differential expression analysis^49,53^. In order to account for technical differences in count depth, we downsampled scRNA-seq count matrices as described above. We did not downsample snRNA-seq count matrices. Differential expression analyses were performed within each time point and within defined subsets of cells or nuclei. Namely, for scRNA-seq data we analyzed cells in the excitatory neuronal lineage (i.e., the “primary trajectory” as defined above) and within cell type partitions that were defined *a priori* based on marker gene expression along pseudotime (Fig. S13C, Table S14). At E12.5, the subplate partition was excluded due to low cell count (fewer than 15 cells). For snRNA-seq data, we analyzed clusters of nuclei passing the criteria defined above (i.e., excluding clusters L5-IT_2, mCtx, Clau, Str, Ambig, Glia+UL, Glia+iN, AG+DL, MG+UL, MG+DL; Fig. 7B, Table S29) .

For each comparison, a generalized linear model was fit for each gene using the “fit_models” function in Monocle 3; we used formula strings to account for sex and sample batch where applicable (e.g., model_formula_str = “∼genotype + sex + batch”, expression_family = “quasipoisson”, clean_model = T). Genes were excluded if they were not detected in at least 10% of cells (or nuclei) in either the wild type or Chd8^+/−^ subpopulations. The number of cells per comparison, the number of genes tested, formula strings used to specify covariates, and the range of pseudotime values corresponding to each cell type partition (for scRNA-seq data) are presented in Table S14, and the number of nuclei per comparison, number of genes tested, and formula string used to specify covariates (for snRNA-seq data) are presented in Table S29. The “coefficient_table” function was used to test each coefficient for significance under the Wald test, *p*-values were adjusted for multiple testing using the Benjamini and Hochberg (BH) method, and results were filtered for the “genotypehet” term (Tables S15-S16, Table S30). Genes with a BH-adjusted *p*-value < 0.05 were labeled as significantly differentially expressed. We used a one-tailed Fisher exact test to determine whether differentially expressed genes were enriched for CHD8 binding targets in the E17.5 wild type mouse cortex, ASD risk-associated genes, DDD genes, and genes associated with risk for epilepsy, schizophrenia, macrocephaly, or microcephaly (fisher.test, alternative = “greater”)^2,6,54–56^. We also tested for enrichment for FMRP binding targets using a one-tailed Fisher exact test^57^. CHD8 binding targets, FMRP binding targets, and NDD risk-associated gene lists are presented in Table S9^2,6,54–57^. For multiple testing correction, we used the “p.adjust” function (method = “BH”).

#### Bulk RNA-sequencing read alignment and differential expression analysis

Illumina paired-end reads from bulk RNA-sequencing experiments were trimmed using Trimmomatic v0.36 (ILLUMINACLIP:TruSeqAdapters.fa:2:30:10 LEADING:3 TRAILING:3 SLIDINGWINDOW:4:15 MINLEN:36) and aligned to the mm39 reference genome using STAR v2.7.9a (GENCODE vM27/Ensembl 104; parameters: --outSAMtype SAM --outSAMunmapped Within --outSAMattributes Standard --quantMode GeneCounts)^97,115^. Genome indices were generated using the recommended settings (--sjdbOverhang 99). SAM files were converted to sorted BAM files using SAMtools, and a count matrix was generated using the featureCounts program in the SourceForge Subread package (v2.0.3; featureCounts -p --countReadPairs -s 2 -T 6 -a gencode.vM27.annotation.gtf -t exon -g gene_id -o featurecount.txt <BAM_FILES>)^98,116^.

We used DESeq2 v1.38.3 for bulk RNA-sequencing differential expression analysis^117^. For each time point, we used the “DESeqDataSetFromMatrix” function to create a DESeqDataSet object with the following design formula: ∼batch + sex + sex:genotype. We tested genes with at least 10 counts in at least 3 out of 12 samples per time point: 20,214 genes in E12.5 cortical data and 19,141 genes in E17.5 cortical data. Differential expression analysis was performed using the likelihood ratio test (LRT), which allowed us to evaluate changes in expression across multiple factor levels (DESeq, test = “LRT”, fitType = “parametric”, sfType = “ratio”, reduced = ∼batch + sex, minReplicatesForReplace = 7, minmu = 0.5). Using this test, genes identified as differentially expressed were those that changed in expression in any direction (up or down) in either sex. We used the function “lfcShrink” to generate results tables with shrunken log2 fold changes for the “sexmale.genotypehet” and “sexfemale.genotypehet” coefficients (type = “apeglm”, lfcThreshold = 0, svalue = F, apeAdapt = T, apeMethod = “nbinomCR”)^118^. Genes with a BH-adjusted *p*-value < 0.05 were labeled as significantly differentially expressed, and directionality was based on average log2 fold change (avg_log2FC = log2((2^log2FC_male + 2^log2FC_female)/2). We used a one-tailed Fisher exact test to determine whether differentially expressed genes were enriched for CHD8 binding targets in E17.5 mouse cortex, ASD risk-associated genes, DDD genes, genes associated with risk for epilepsy, schizophrenia, macrocephaly, or microcephaly, and FMRP binding targets as described above (Table S9)^2,6,54–57^.

#### Gene ontology enrichment analysis of differentially expressed genes

For single-cell, single-nucleus, and bulk RNA-sequencing datasets, we performed GO:BP functional enrichment analysis using the g:GOSt tool in the g:Profiler web toolset (https://biit.cs.ut.ee/gprofiler/gost), implemented via the gprofiler2 interface. For the snRNA-seq dataset, we also tested for enrichment of pathway terms from the KEGG and Reactome databases (sources = c(“GO:BP”, “KEGG”, “REAC”)). To test differentially expressed genes in each cell type of interest for functional enrichment, we used significantly upregulated or downregulated genes in the *Chd8*^+/-^ cortex at each time point (E12.5, E14.5, E16.0, E17.5, and P25), as identified by Monocle 3 for single-cell and single-nucleus data, or DESeq2 for bulk data. All genes tested for differential expression within the corresponding time point and cell type (in the case of scRNA-seq and snRNA-seq datasets), were used as a statistical background. For multiple testing correction and significance calling, we used default parameters (user_threshold = 0.05, correction_method = “g_SCS”).

We then used the Revigo web tool (http://revigo.irb.hr/) to summarize the resulting lists of significant GO terms (resulting list size = Tiny, species = Mus musculus, semantic similarity measure = SimRel; for bulk RNA-seq data: same parameters, except for resulting list size = Small). For scRNA-seq datasets, we used Revigo to summarize a combined list of all GO:BP terms that were significantly enriched among upregulated or downregulated genes in at least one cell type, at any time point (252 GO:BP terms). We provided Revigo with values for each GO term equal to -log10(adjusted *p-*value), using the adjusted *p*-values generated by g:Profiler from functional enrichment analysis of downregulated genes identified in E12.5 radial glia. After grouping terms based on semantic similarity, these values were used by Revigo to select a representative “parent term” from each group. The full list of significant GO:BP terms, representative terms, and adjusted *p*-values generated by g:Profiler are provided in Table S22.

We also used Revigo to summarize the lists of GO:BP terms that were significantly enriched among upregulated genes (68 GO:BP terms) or downregulated genes (169 GO:BP terms) identified in the E12.5 bulk cortex. We provided values for each GO:BP term equal to -log10(adjusted *p-*value); after grouping terms based on semantic similarity, these values were used to select a parent term from each group. The term “cell part morphogenesis” (GO:0032990) is a duplicate of the term “cellular component morphogenesis” (GO:0032989). The term “regulation of neurotransmitter levels” (GO:0001505) is obsolete and was removed from the dataset by Revigo. The full list of significantly enriched GO:BP terms are provided in Table S23. No GO:BP terms were significantly enriched among upregulated or downregulated genes identified in the E17.5 bulk cortex.

For snRNA-seq datasets, Revigo was employed to summarize significant GO:BP terms enriched across multiple cell types, which were split into three groups: terms enriched by downregulated genes across multiple clusters (15 GO:BP terms), terms enriched by both downregulated and upregulated genes across multiple clusters (3 GO:BP terms), and terms enriched by upregulated genes across multiple clusters (5 GO:BP terms) (Fig. S23). Each of these three groups of GO:BP terms was independently submitted to Revigo, to avoid removing GO:BP terms with distinct patterns of enrichment across clusters, and we provided values for each GO:BP term equal to the number of cell types enriched for that term (Fig. 7D, Fig. S23). After grouping terms based on semantic similarity, these values were used to select a parent term from each group. For GO:BP terms enriched by downregulated genes across multiple clusters, “GO:BP synaptic signaling,” “GO:BP trans-synaptic signaling,” and “GO:BP anterograde trans-synaptic signaling” were collapsed under the parent term “GO:BP chemical synaptic transmission”; “GO:BP synapse assembly” and “GO:BP cell junction assembly” were collapsed under the parent term “GO:BP synapse organization”; and “GO:BP regulation of trans-synaptic signaling” was collapsed under the parent term “GO:BP modulation of chemical synaptic transmission.” For GO:BP terms enriched by both downregulated and upregulated genes across multiple clusters, the terms “GO:BP neurogenesis” and “GO:BP generation of neurons” were collapsed under the parent term “GO:BP neuron development.” For GO:BP terms enriched by upregulated genes across multiple clusters, “GO:BP neuron differentiation” and “GO:BP nervous system development” were collapsed under the parent term “GO:BP neuron projection development.” The full list of significantly enriched GO terms is provided in Tables S33-S34 and Figure S23.

#### Gene Set Enrichment Analysis (GSEA)

We performed GSEA using the GSEAPreranked module in GSEA v4.3.2 for command line^60,61^. To run GSEA, we provided GSEAPreranked with a GeneMatrix (GMX) file containing ASD risk-associated genes, DDD genes, and genes associated with risk for epilepsy, schizophrenia, macrocephaly, or microcephaly (Table S9) and pre-ranked gene list files (RNK) generated from Monocle 3 differential expression results (for single-cell and single-nucleus datasets; Tables S15-S16, Table S30) or DESeq2 differential expression results (for bulk datasets; Tables S19 & S35). For each gene included in the differential expression analyses, we calculated a rank metric by multiplying the sign of the log2 fold change by the negative log10 *p*-value generated by Monocle 3 or DESeq2 (rank_metric = sign(avg_log2FC) * -log10(p_value)). To avoid infinite values, genes with a nominal *p*-value of zero were assigned a rank_metric = sign(avg_log2FC) * 350. For each pre-ranked gene list, GSEA was run from the command line using the following parameters: gsea-cli.sh GSEAPreranked -gmx <GMX_FILE> -rnk <RNK_FILE> -rpt_label <LABEL> -out <OUT_FILE> -rnd_seed 149 -collapse No_Collapse -mode Abs_max_of_probes -norm meandiv -nperm 1000 -scoring_scheme weighted -create_svgs false -include_only_symbols true -make_sets true -plot_top_x 10 -set_max 500 -set_min 1 -zip_report true. GSEA calculates a false discovery rate (FDR) equal to the ratio of two distributions: “(1) the actual enrichment score versus the enrichment scores for all gene sets against all permutations of the dataset and (2) the actual enrichment score versus the enrichment scores of all gene sets against the actual dataset” (https://docs.gsea-msigdb.org/#GSEA/GSEA_User_Guide/). We labeled gene sets that had an FDR q-value < 0.05 as significantly enriched: “NEG” if they were enriched at the bottom of the ranked list (i.e., among downregulated genes) and “POS” if they were enriched at the top of the ranked list (i.e., among upregulated genes).

## Acknowledgements

We thank A. Tong, D. van Dijk, and S. Krishnaswamy for helpful discussions and feedback on the analyses, and members of the Noonan laboratory for comments on the manuscript. We also thank K. Bilguvar, C. Castaldi, G. Wang, and S. Mane at the Yale Center for Genome Analysis for assistance in generating single-cell, single-nucleus, and bulk RNA-sequencing libraries and for sequencing data for scRNA-seq, snRNA-seq, and bulk RNA-seq experiments.

## Data availability and materials

All RNA-seq data have been deposited under GEO accessions GSE273271 (scRNA-seq), GSE273765 (snRNA-seq), and GSE273270 (bulk RNA-seq). All code for the study is available at GitHub (https://github.com/NoonanLab/Yim_et_al_Chd8) and Zenodo (https://zenodo.org/doi/10.5281/zenodo.13324274)

## Author contributions

K.M.Y., R.A.M., and J.P.N. conceived of and designed the study. T.N. and R.A.M. generated the *Chd8* knockout model. R.A.M. and G.H.-T. performed dissections, cell dissociations, and cell suspension quality control of embryonic mouse cortical samples submitted for scRNA-sequencing. M.B. dissected, isolated nuclei from, and assessed nuclear suspension quality of juvenile mouse cortical samples submitted for snRNA-sequencing. R.A.M. obtained RNA for bulk RNA-sequencing from the embryonic and juvenile mouse cortex. K.M.Y. conceived of and supervised all computational analyses in the study, identified and characterized cell types and developmental trajectories in the embryonic data, developed and implemented the metagene analyses, and carried out differential expression analyses between the wild type and *Chd8^+/−^* cortex. M.B. analyzed snRNA-seq data from the juvenile cortex and characterized differentially expressed genes. M.K. carried out immunohistochemistry studies. M.F.R.L. carried out Western blot analyses of CHD8 expression. M.F.R.L. and G.H.-T. performed genotyping throughout the project. K.M.Y., M.B., M.F.R.L., and J.P.N. wrote the manuscript with input from all authors.

## References

1. Sanders, S.J., Murtha, M.T., Gupta, A.R., Murdoch, J.D., Raubeson, M.J., Willsey, A.J., Ercan-Sencicek, A.G., DiLullo, N.M., Parikshak, N.N., Stein, J.L., et al. (2012). De novo mutations revealed by whole-exome sequencing are strongly associated with autism. Nature 485, 237–241. 10.1038/nature10945.

2. Satterstrom, F.K., Kosmicki, J.A., Wang, J., Breen, M.S., Rubeis, S.D., An, J.-Y., Peng, M., Collins, R., Grove, J., Klei, L., et al. (2020). Large-Scale Exome Sequencing Study Implicates Both Developmental and Functional Changes in the Neurobiology of Autism. Cell 180, 568–584.e23. 10.1016/j.cell.2019.12.036.

3. Rubeis, S.D., He, X., Goldberg, A.P., Poultney, C.S., Samocha, K., Cicek, A.E., Kou, Y., Liu, L., Fromer, M., Walker, S., et al. (2014). Synaptic, transcriptional and chromatin genes disrupted in autism. Nature 515, 209–215. 10.1038/nature13772.

4. Iossifov, I., O’Roak, B.J., Sanders, S.J., Ronemus, M., Krumm, N., Levy, D., Stessman, H.A., Witherspoon, K.T., Vives, L., Patterson, K.E., et al. (2014). The contribution of de novo coding mutations to autism spectrum disorder. Nature 515, 216–221. 10.1038/nature13908.

5. Fitzgerald, T.W., Gerety, S.S., Jones, W.D., Kogelenberg, M. van, King, D.A., McRae, J., Morley, K.I., Parthiban, V., Al-Turki, S., Ambridge, K., et al. (2015). Large-scale discovery of novel genetic causes of developmental disorders. Nature 519, 223–228. 10.1038/nature14135.

6. McRae, J.F., Clayton, S., Fitzgerald, T.W., Kaplanis, J., Prigmore, E., Rajan, D., Sifrim, A., Aitken, S., Akawi, N., Alvi, M., et al. (2017). Prevalence and architecture of de novo mutations in developmental disorders. Nature 542, 433–438. 10.1038/nature21062.

7. Abrahams, B.S., Arking, D.E., Campbell, D.B., Mefford, H.C., Morrow, E.M., Weiss, L.A., Menashe, I., Wadkins, T., Banerjee-Basu, S., and Packer, A. (2013). SFARI Gene 2.0: a community-driven knowledgebase for the autism spectrum disorders (ASDs). Mol. Autism 4, 36. 10.1186/2040-2392-4-36.

8. SFARI Gene 3.0 2023 Q3 release. Available at gene.sfari.org.

9. Parikshak, N.N., Luo, R., Zhang, A., Won, H., Lowe, J.K., Chandran, V., Horvath, S., and Geschwind, D.H. (2013). Integrative Functional Genomic Analyses Implicate Specific Molecular Pathways and Circuits in Autism. Cell 155, 1008–1021. 10.1016/j.cell.2013.10.031.

10. Willsey, A.J., Sanders, S.J., Li, M., Dong, S., Tebbenkamp, A.T., Muhle, R.A., Reilly, S.K., Lin, L., Fertuzinhos, S., Miller, J.A., et al. (2013). Coexpression Networks Implicate Human Midfetal Deep Cortical Projection Neurons in the Pathogenesis of Autism. Cell 155, 997–1007. 10.1016/j.cell.2013.10.020.

11. Li, M., Santpere, G., Kawasawa, Y.I., Evgrafov, O.V., Gulden, F.O., Pochareddy, S., Sunkin, S.M., Li, Z., Shin, Y., Zhu, Y., et al. (2018). Integrative functional genomic analysis of human brain development and neuropsychiatric risks. Science 362, eaat7615. 10.1126/science.aat7615.

12. Velmeshev, D., Schirmer, L., Jung, D., Haeussler, M., Perez, Y., Mayer, S., Bhaduri, A., Goyal, N., Rowitch, D.H., and Kriegstein, A.R. (2019). Single-cell genomics identifies cell type– specific molecular changes in autism. Science 364, 685–689. 10.1126/science.aav8130.

13. O’Roak, B.J., Vives, L., Fu, W., Egertson, J.D., Stanaway, I.B., Phelps, I.G., Carvill, G., Kumar, A., Lee, C., Ankenman, K., et al. (2012). Multiplex Targeted Sequencing Identifies Recurrently Mutated Genes in Autism Spectrum Disorders. Science 338, 1619–1622. 10.1126/science.1227764.

14. Katayama, Y., Nishiyama, M., Shoji, H., Ohkawa, Y., Kawamura, A., Sato, T., Suyama, M., Takumi, T., Miyakawa, T., and Nakayama, K.I. (2016). CHD8 haploinsufficiency results in autistic-like phenotypes in mice. Nature 537, 675–679. 10.1038/nature19357.

15. Nishiyama, M., Skoultchi, A.I., and Nakayama, K.I. (2012). Histone H1 Recruitment by CHD8 Is Essential for Suppression of the Wnt–β-Catenin Signaling Pathway. Mol. Cell. Biol. 32, 501–512. 10.1128/mcb.06409-11.

16. Bernier, R., Golzio, C., Xiong, B., Stessman, H.A., Coe, B.P., Penn, O., Witherspoon, K., Gerdts, J., Baker, C., Vulto-van Silfhout, A.T., et al. (2014). Disruptive CHD8 Mutations Define a Subtype of Autism Early in Development. Cell 158, 263–276. 10.1016/j.cell.2014.06.017.

17. Gompers, A.L., Su-Feher, L., Ellegood, J., Copping, N.A., Riyadh, M.A., Stradleigh, T.W., Pride, M.C., Schaffler, M.D., Wade, A.A., Catta-Preta, R., et al. (2017). Germline Chd8 haploinsufficiency alters brain development in mouse. Nat. Neurosci. 20, 1062–1073. 10.1038/nn.4592.

18. Cotney, J., Muhle, R.A., Sanders, S.J., Liu, L., Willsey, A.J., Niu, W., Liu, W., Klei, L., Lei, J., Yin, J., et al. (2015). The autism-associated chromatin modifier CHD8 regulates other autism risk genes during human neurodevelopment. Nat. Commun. 6, 6404. 10.1038/ncomms7404.

19. Sugathan, A., Biagioli, M., Golzio, C., Erdin, S., Blumenthal, I., Manavalan, P., Ragavendran, A., Brand, H., Lucente, D., Miles, J., et al. (2014). CHD8 regulates neurodevelopmental pathways associated with autism spectrum disorder in neural progenitors. Proc. Natl. Acad. Sci. 111, E4468–E4477. 10.1073/pnas.1405266111.

20. Suetterlin, P., Hurley, S., Mohan, C., Riegman, K.L.H., Pagani, M., Caruso, A., Ellegood, J., Galbusera, A., Crespo-Enriquez, I., Michetti, C., et al. (2018). Altered Neocortical Gene Expression, Brain Overgrowth and Functional Over-Connectivity in Chd8 Haploinsufficient Mice. Cereb. Cortex 28, 2192–2206. 10.1093/cercor/bhy058.

21. Zhao, C., Dong, C., Frah, M., Deng, Y., Marie, C., Zhang, F., Xu, L., Ma, Z., Dong, X., Lin, Y., et al. (2018). Dual Requirement of CHD8 for Chromatin Landscape Establishment and Histone Methyltransferase Recruitment to Promote CNS Myelination and Repair. Dev. Cell 45, 753–768.e8. 10.1016/j.devcel.2018.05.022.

22. Marie, C., Clavairoly, A., Frah, M., Hmidan, H., Yan, J., Zhao, C., Steenwinckel, J.V., Daveau, R., Zalc, B., Hassan, B., et al. (2018). Oligodendrocyte precursor survival and differentiation requires chromatin remodeling by Chd7 and Chd8. Proc. Natl. Acad. Sci. 115, E8246–E8255. 10.1073/pnas.1802620115.

23. Jin, X., Simmons, S.K., Guo, A., Shetty, A.S., Ko, M., Nguyen, L., Jokhi, V., Robinson, E., Oyler, P., Curry, N., et al. (2020). In vivo Perturb-Seq reveals neuronal and glial abnormalities associated with autism risk genes. Science 370, eaaz6063. 10.1126/science.aaz6063.

24. Durak, O., Gao, F., Kaeser-Woo, Y.J., Rueda, R., Martorell, A.J., Nott, A., Liu, C.Y., Watson, L.A., and Tsai, L.-H. (2016). Chd8 mediates cortical neurogenesis via transcriptional regulation of cell cycle and Wnt signaling. Nat. Neurosci. 19, 1477–1488. 10.1038/nn.4400.

25. Geller, E., Gockley, J., Emera, D., Uebbing, S., Cotney, J., and Noonan, J.P. (2019). Massively parallel disruption of enhancers active during human corticogenesis. bioRxiv, 852673. 10.1101/852673.

26. Shi, X., Lu, C., Corman, A., Nikish, A., Zhou, Y., Platt, R.J., Iossifov, I., Zhang, F., Pan, J.Q., and Sanjana, N.E. (2023). Heterozygous deletion of the autism-associated gene CHD8 impairs synaptic function through widespread changes in gene expression and chromatin compaction. Am. J. Hum. Genet. 110, 1750–1768. 10.1016/j.ajhg.2023.09.004.

27. Paulsen, B., Velasco, S., Kedaigle, A.J., Pigoni, M., Quadrato, G., Deo, A.J., Adiconis, X., Uzquiano, A., Sartore, R., Yang, S.M., et al. (2022). Autism genes converge on asynchronous development of shared neuron classes. Nature 602, 268–273. 10.1038/s41586-021-04358-6.

28. Nishiyama, M., Oshikawa, K., Tsukada, Y., Nakagawa, T., Iemura, S., Natsume, T., Fan, Y., Kikuchi, A., Skoultchi, A.I., and Nakayama, K.I. (2009). CHD8 suppresses p53-mediated apoptosis through histone H1 recruitment during early embryogenesis. Nat. Cell Biol. 11, 172–182. 10.1038/ncb1831.

29. Platt, R.J., Zhou, Y., Slaymaker, I.M., Shetty, A.S., Weisbach, N.R., Kim, J.-A., Sharma, J., Desai, M., Sood, S., Kempton, H.R., et al. (2017). Chd8 Mutation Leads to Autistic-like Behaviors and Impaired Striatal Circuits. Cell Rep. 19, 335–350. 10.1016/j.celrep.2017.03.052.

30. Hafemeister, C., and Satija, R. (2019). Normalization and variance stabilization of single-cell RNA-seq data using regularized negative binomial regression. Genome Biol. 20, 296. 10.1186/s13059-019-1874-1.

31. Choudhary, S., and Satija, R. (2022). Comparison and evaluation of statistical error models for scRNA-seq. 23, 27. 10.1186/s13059-021-02584-9.

32. Hao, Y., Hao, S., Andersen-Nissen, E., Mauck, W.M., Zheng, S., Butler, A., Lee, M.J., Wilk, A.J., Darby, C., Zager, M., et al. (2021). Integrated analysis of multimodal single-cell data. Cell 184, 3573–3587.e29. 10.1016/j.cell.2021.04.048.

33. Stuart, T., Butler, A., Hoffman, P., Hafemeister, C., Papalexi, E., Mauck, W.M., Hao, Y., Stoeckius, M., Smibert, P., and Satija, R. (2019). Comprehensive Integration of Single-Cell Data. Cell 177, 1888–1902.e21. 10.1016/j.cell.2019.05.031.

34. Butler, A., Hoffman, P., Smibert, P., Papalexi, E., and Satija, R. (2018). Integrating single-cell transcriptomic data across different conditions, technologies, and species. 36, 411–420. 10.1038/nbt.4096.

35. McInnes, L., Healy, J., and Melville, J. (2018). UMAP: Uniform Manifold Approximation and Projection for Dimension Reduction. arXiv. 10.48550/arxiv.1802.03426.

36. Ellis, P., Fagan, B.M., Magness, S.T., Hutton, S., Taranova, O., Hayashi, S., McMahon, A., Rao, M., and Pevny, L. (2004). SOX2, a persistent marker for multipotential neural stem cells derived from embryonic stem cells, the embryo or the adult. Dev Neurosci 26, 148–165. 10.1159/000082134.

37. Götz, M., Stoykova, A., and Gruss, P. (1998). Pax6 Controls Radial Glia Differentiation in the Cerebral Cortex. 21, 1031–1044. 10.1016/s0896-6273(00)80621-2.

38. Hevner, R.F. (2019). Intermediate progenitors and Tbr2 in cortical development. J Anat 235, 616–625. 10.1111/joa.12939.

39. Kwan, K.Y., Sestan, N., and Anton, E.S. (2012). Transcriptional co-regulation of neuronal migration and laminar identity in the neocortex. Development 139, 1535–1546. 10.1242/dev.069963.

40. Tarabykin, V., Stoykova, A., Usman, N., and Gruss, P. (2001). Cortical upper layer neurons derive from the subventricular zone as indicated by Svet1 gene expression. 128, 1983–1993. 10.1242/dev.128.11.1983.

41. Stühmer, T., Puelles, L., Ekker, M., and Rubenstein, J.L.R. (2002). Expression from a Dlx Gene Enhancer Marks Adult Mouse Cortical GABAergic Neurons. 12, 75–85. 10.1093/cercor/12.1.75.

42. Baek, S.-H., Maiorino, E., Kim, H., Glass, K., Raby, B.A., and Yuan, K. (2022). Single Cell Transcriptomic Analysis Reveals Organ Specific Pericyte Markers and Identities. Frontiers Cardiovasc Medicine 9, 876591. 10.3389/fcvm.2022.876591.

43. Kalucka, J., Rooij, L.P.M.H. de, Goveia, J., Rohlenova, K., Dumas, S.J., Meta, E., Conchinha, N.V., Taverna, F., Teuwen, L.-A., Veys, K., et al. (2020). Single-Cell Transcriptome Atlas of Murine Endothelial Cells. Cell 180, 764–779.e20. 10.1016/j.cell.2020.01.015.

44. Miquelajauregui, A., Varela-Echavarria, A., Ceci, M.L., Garcia-Moreno, F., Ricano, I., Hoang, K., Frade-Perez, D., Portera-Cailliau, C., Tamariz, E., Carlos, J.A.D., et al. (2010). LIM-Homeobox Gene Lhx5 Is Required for Normal Development of Cajal-Retzius Cells. J Neurosci 30, 10551–10562. 10.1523/jneurosci.5563-09.2010.

45. Lu, Q.R., Yuk, D., Alberta, J.A., Zhu, Z., Pawlitzky, I., Chan, J., McMahon, A.P., Stiles, C.D., and Rowitch, D.H. (2000). Sonic Hedgehog–Regulated Oligodendrocyte Lineage Genes Encoding bHLH Proteins in the Mammalian Central Nervous System. 25, 317–329. 10.1016/s0896-6273(00)80897-1.

46. Hammond, T.R., Dufort, C., Dissing-Olesen, L., Giera, S., Young, A., Wysoker, A., Walker, A.J., Gergits, F., Segel, M., Nemesh, J., et al. (2019). Single-Cell RNA Sequencing of Microglia throughout the Mouse Lifespan and in the Injured Brain Reveals Complex Cell-State Changes. Immunity 50, 253–271.e6. 10.1016/j.immuni.2018.11.004.

47. Moon, K.R., Dijk, D. van, Wang, Z., Gigante, S., Burkhardt, D.B., Chen, W.S., Yim, K., Elzen, A. van den, Hirn, M.J., Coifman, R.R., et al. (2019). Visualizing structure and transitions in high-dimensional biological data. Nat. Biotechnol. 37, 1482–1492. 10.1038/s41587-019-0336-3.

48. Hastie, T., and Stuetzle, W. (1989). Principal Curves. J Am Stat Assoc 84, 502. 10.2307/2289936.

49. Trapnell, C., Cacchiarelli, D., Grimsby, J., Pokharel, P., Li, S., Morse, M., Lennon, N.J., Livak, K.J., Mikkelsen, T.S., and Rinn, J.L. (2014). The dynamics and regulators of cell fate decisions are revealed by pseudotemporal ordering of single cells. Nat. Biotechnol. 32, 381–386. 10.1038/nbt.2859.

50. Götz, M., Stoykova, A., and Gruss, P. (1998). Pax6 Controls Radial Glia Differentiation in the Cerebral Cortex. Neuron 21, 1031–1044. 10.1016/s0896-6273(00)80621-2.

51. Hevner, R.F., Shi, L., Justice, N., Hsueh, Y.-P., Sheng, M., Smiga, S., Bulfone, A., Goffinet, A.M., Campagnoni, A.T., and Rubenstein, J.L.R. (2001). Tbr1 Regulates Differentiation of the Preplate and Layer 6. Neuron 29, 353–366. 10.1016/s0896-6273(01)00211-2.

52. Polioudakis, D., Torre-Ubieta, L. de la, Langerman, J., Elkins, A.G., Shi, X., Stein, J.L., Vuong, C.K., Nichterwitz, S., Gevorgian, M., Opland, C.K., et al. (2019). A Single-Cell Transcriptomic Atlas of Human Neocortical Development during Mid-gestation. Neuron 103, 785–801.e8. 10.1016/j.neuron.2019.06.011.

53. Cao, J., Spielmann, M., Qiu, X., Huang, X., Ibrahim, D.M., Hill, A.J., Zhang, F., Mundlos, S., Christiansen, L., Steemers, F.J., et al. (2019). The single-cell transcriptional landscape of mammalian organogenesis. Nature 566, 496–502. 10.1038/s41586-019-0969-x.

54. Singh, T., Poterba, T., Curtis, D., Akil, H., Eissa, M.A., Barchas, J.D., Bass, N., Bigdeli, T.B., Breen, G., Bromet, E.J., et al. (2022). Rare coding variants in ten genes confer substantial risk for schizophrenia. Nature 604, 509–516. 10.1038/s41586-022-04556-w.

55. Tatton-Brown, K., Loveday, C., Yost, S., Clarke, M., Ramsay, E., Zachariou, A., Elliott, A., Wylie, H., Ardissone, A., Rittinger, O., et al. (2017). Mutations in Epigenetic Regulation Genes Are a Major Cause of Overgrowth with Intellectual Disability. Am. J. Hum. Genet. 100, 725– 736. 10.1016/j.ajhg.2017.03.010.

56. Jayaraman, D., Bae, B.-I., and Walsh, C.A. (2018). The Genetics of Primary Microcephaly. Annu. Rev. Genom. Hum. Genet. 19, 1–24. 10.1146/annurev-genom-083117-021441.

57. Darnell, J.C., Van Driesche, S.J., Zhang, C., Hung, K.Y.S., Mele, A., Fraser, C.E., Stone, E.F., Chen, C., Fak, J.J., Chi, S.W., et al. (2011). FMRP Stalls Ribosomal Translocation on mRNAs Linked to Synaptic Function and Autism. Cell 146, 247–261. 10.1016/j.cell.2011.06.013.

58. Verkerk, A.J.M.H., Pieretti, M., Sutcliffe, J.S., Fu, Y.-H., Kuhl, D.P.A., Pizzuti, A., Reiner, O., Richards, S., Victoria, M.F., Zhang, F., et al. (1991). Identification of a gene (FMR-1) containing a CGG repeat coincident with a breakpoint cluster region exhibiting length variation in fragile X syndrome. Cell 65, 905–914. 10.1016/0092-8674(91)90397-h.

59. Niu, M., Han, Y., Dy, A.B.C., Du, J., Jin, H., Qin, J., Zhang, J., Li, Q., and Hagerman, R.J. (2017). Autism Symptoms in Fragile X Syndrome. J. Child Neurol. 32, 903–909. 10.1177/0883073817712875.

60. Subramanian, A., Tamayo, P., Mootha, V.K., Mukherjee, S., Ebert, B.L., Gillette, M.A., Paulovich, A., Pomeroy, S.L., Golub, T.R., Lander, E.S., et al. (2005). Gene set enrichment analysis: A knowledge-based approach for interpreting genome-wide expression profiles. Proc. Natl. Acad. Sci. 102, 15545–15550. 10.1073/pnas.0506580102.

61. Mootha, V.K., Lindgren, C.M., Eriksson, K.-F., Subramanian, A., Sihag, S., Lehar, J., Puigserver, P., Carlsson, E., Ridderstråle, M., Laurila, E., et al. (2003). PGC-1α-responsive genes involved in oxidative phosphorylation are coordinately downregulated in human diabetes. Nat. Genet. 34, 267–273. 10.1038/ng1180.

62. Yao, Z., Velthoven, C.T.J. van, Nguyen, T.N., Goldy, J., Sedeno-Cortes, A.E., Baftizadeh, F., Bertagnolli, D., Casper, T., Chiang, M., Crichton, K., et al. (2021). A taxonomy of transcriptomic cell types across the isocortex and hippocampal formation. Cell 184, 3222–3241.e26. 10.1016/j.cell.2021.04.021.

63. Yao, Z., Liu, H., Xie, F., Fischer, S., Adkins, R.S., Aldridge, A.I., Ament, S.A., Bartlett, A., Behrens, M.M., Berge, K.V. den, et al. (2021). A transcriptomic and epigenomic cell atlas of the mouse primary motor cortex. Nature 598, 103–110. 10.1038/s41586-021-03500-8.

64. Yao, Z., Velthoven, C.T.J. van, Kunst, M., Zhang, M., McMillen, D., Lee, C., Jung, W., Goldy, J., Abdelhak, A., Aitken, M., et al. (2023). A high-resolution transcriptomic and spatial atlas of cell types in the whole mouse brain. Nature 624, 317–332. 10.1038/s41586-023-06812-z.

65. Tasic, B., Yao, Z., Graybuck, L.T., Smith, K.A., Nguyen, T.N., Bertagnolli, D., Goldy, J., Garren, E., Economo, M.N., Viswanathan, S., et al. (2018). Shared and distinct transcriptomic cell types across neocortical areas. Nature 563, 72–78. 10.1038/s41586-018-0654-5.

66. Bhattacherjee, A., Djekidel, M.N., Chen, R., Chen, W., Tuesta, L.M., and Zhang, Y. (2019). Cell type-specific transcriptional programs in mouse prefrontal cortex during adolescence and addiction. Nat. Commun. 10, 4169. 10.1038/s41467-019-12054-3.

67. Bruguier, H., Suarez, R., Manger, P., Hoerder-Suabedissen, A., Shelton, A.M., Oliver, D.K., Packer, A.M., Ferran, J.L., García-Moreno, F., Puelles, L., et al. (2020). In search of common developmental and evolutionary origin of the claustrum and subplate. J. Comp. Neurol. 528, 2956–2977. 10.1002/cne.24922.

68. Sullivan, K.E., Kraus, L., Kapustina, M., Wang, L., Stach, T.R., Lemire, A.L., Clements, J., and Cembrowski, M.S. (2023). Sharp cell-type-identity changes differentiate the retrosplenial cortex from the neocortex. Cell Rep. 42, 112206. 10.1016/j.celrep.2023.112206.

69. Lein, E.S., Hawrylycz, M.J., Ao, N., Ayres, M., Bensinger, A., Bernard, A., Boe, A.F., Boguski, M.S., Brockway, K.S., Byrnes, E.J., et al. (2007). Genome-wide atlas of gene expression in the adult mouse brain. Nature 445, 168–176. 10.1038/nature05453.

70. Castiglioni, V., Faedo, A., Onorati, M., Bocchi, V.D., Li, Z., Iennaco, R., Vuono, R., Bulfamante, G.P., Muzio, L., Martino, G., et al. (2019). Dynamic and Cell-Specific DACH1 Expression in Human Neocortical and Striatal Development. Cereb. Cortex 29, 2115–2124. 10.1093/cercor/bhy092.

71. Tasic, B., Menon, V., Nguyen, T.N., Kim, T.K., Jarsky, T., Yao, Z., Levi, B., Gray, L.T., Sorensen, S.A., Dolbeare, T., et al. (2016). Adult mouse cortical cell taxonomy revealed by single cell transcriptomics. Nat. Neurosci. 19, 335–346. 10.1038/nn.4216.

72. Batiuk, M.Y., Martirosyan, A., Wahis, J., Vin, F. de, Marneffe, C., Kusserow, C., Koeppen, J., Viana, J.F., Oliveira, J.F., Voet, T., et al. (2020). Identification of region-specific astrocyte subtypes at single cell resolution. Nat. Commun. 11, 1220. 10.1038/s41467-019-14198-8.

73. Marques, S., Zeisel, A., Codeluppi, S., Bruggen, D. van, Falcão, A.M., Xiao, L., Li, H., Häring, M., Hochgerner, H., Romanov, R.A., et al. (2016). Oligodendrocyte heterogeneity in the mouse juvenile and adult central nervous system. Science 352, 1326–1329. 10.1126/science.aaf6463.

74. Trimm, E., and Red-Horse, K. (2023). Vascular endothelial cell development and diversity. Nat. Rev. Cardiol. 20, 197–210. 10.1038/s41569-022-00770-1.

75. Hansen, K.B., Yi, F., Perszyk, R.E., Menniti, F.S., and Traynelis, S.F. (2017). NMDA Receptors, Methods and Protocols. Methods Mol. Biol. 1677, 1–80. 10.1007/978-1-4939-7321-7_1.

76. Henley, J.M., and Wilkinson, K.A. (2016). Synaptic AMPA receptor composition in development, plasticity and disease. Nat. Rev. Neurosci. 17, 337–350. 10.1038/nrn.2016.37.

77. Negrete-Díaz, J.V., Falcón-Moya, R., and Rodríguez-Moreno, A. (2022). Kainate receptors: from synaptic activity to disease. FEBS J. 289, 5074–5088. 10.1111/febs.16081.

78. Niswender, C.M., and Conn, P.J. (2010). Metabotropic Glutamate Receptors: Physiology, Pharmacology, and Disease. Annu. Rev. Pharmacol. Toxicol. 50, 295–322. 10.1146/annurev.pharmtox.011008.145533.

79. Greger, I.H., Watson, J.F., and Cull-Candy, S.G. (2017). Structural and Functional Architecture of AMPA-Type Glutamate Receptors and Their Auxiliary Proteins. Neuron 94, 713–730. 10.1016/j.neuron.2017.04.009.

80. Kaizuka, T., and Takumi, T. (2018). Postsynaptic density proteins and their involvement in neurodevelopmental disorders. J. Biochem. 163, 447–455. 10.1093/jb/mvy022.

81. Henderson, N.T., and Dalva, M.B. (2018). EphBs and ephrin-Bs: Trans-synaptic organizers of synapse development and function. Mol. Cell. Neurosci. 91, 108–121. 10.1016/j.mcn.2018.07.002.

82. Naito, Y., Lee, A.K., and Takahashi, H. (2017). Emerging roles of the neurotrophin receptor TrkC in synapse organization. Neurosci. Res. 116, 10–17. 10.1016/j.neures.2016.09.009.

83. Brouwer, M., Farzana, F., Koopmans, F., Chen, N., Brunner, J.W., Oldani, S., Li, K.W., Weering, J.R. van, Smit, A.B., Toonen, R.F., et al. (2019). SALM1 controls synapse development by promoting F-actin/PIP2-dependent Neurexin clustering. EMBO J. 38, e101289. 10.15252/embj.2018101289.

84. Roppongi, R.T., Dhume, S.H., Padmanabhan, N., Silwal, P., Zahra, N., Karimi, B., Bomkamp, C., Patil, C.S., Champagne-Jorgensen, K., Twilley, R.E., et al. (2020). LRRTMs Organize Synapses through Differential Engagement of Neurexin and PTPσ. Neuron 106, 108–125.e12. 10.1016/j.neuron.2020.01.003.

85. Akaneya, Y., Sohya, K., Kitamura, A., Kimura, F., Washburn, C., Zhou, R., Ninan, I., Tsumoto, T., and Ziff, E.B. (2010). Ephrin-A5 and EphA5 Interaction Induces Synaptogenesis during Early Hippocampal Development. PLoS ONE 5, e12486. 10.1371/journal.pone.0012486.

86. Liu, L., Lei, J., Sanders, S.J., Willsey, A.J., Kou, Y., Cicek, A.E., Klei, L., Lu, C., He, X., Li, M., et al. (2014). DAWN: a framework to identify autism genes and subnetworks using gene expression and genetics. Mol. Autism 5, 22. 10.1186/2040-2392-5-22.

87. Wamsley, B., Bicks, L., Cheng, Y., Kawaguchi, R., Quintero, D., Margolis, M., Grundman, J., Liu, J., Xiao, S., Hawken, N., et al. (2024). Molecular cascades and cell type–specific signatures in ASD revealed by single-cell genomics. Science 384, eadh2602. 10.1126/science.adh2602.

88. Catania, M.V., Landwehrmeyer, G.B., Testa, C.M., Standaert, D.G., Penney, J.B., and Young, A.B. (1994). Metabotropic glutamate receptors are differentially regulated during development. Neuroscience 61, 481–495. 10.1016/0306-4522(94)90428-6.

89. Wong, H., Liu, X., Matos, M.F., Chan, S.F., Pérez-Otaño, I., Boysen, M., Cui, J., Nakanishi, N., Trimmer, J.S., Jones, E.G., et al. (2002). Temporal and regional expression of NMDA receptor subunit NR3A in the mammalian brain. J. Comp. Neurol. 450, 303–317. 10.1002/cne.10314.

90. Pérez-Otaño, I., Larsen, R.S., and Wesseling, J.F. (2016). Emerging roles of GluN3-containing NMDA receptors in the CNS. Nat. Rev. Neurosci. 17, 623–635. 10.1038/nrn.2016.92.

91. Luchkina, N.V., Coleman, S.K., Huupponen, J., Cai, C., Kivistö, A., Taira, T., Keinänen, K., and Lauri, S.E. (2017). Molecular mechanisms controlling synaptic recruitment of GluA4 subunit-containing AMPA-receptors critical for functional maturation of CA1 glutamatergic synapses. Neuropharmacology 112, 46–56. 10.1016/j.neuropharm.2016.04.049.

92. Subtil-Rodríguez, A., Vázquez-Chávez, E., Ceballos-Chávez, M., Rodríguez-Paredes, M., Martín-Subero, J.I., Esteller, M., and Reyes, J.C. (2014). The chromatin remodeller CHD8 is required for E2F-dependent transcription activation of S-phase genes. Nucleic Acids Res. 42, 2185–2196. 10.1093/nar/gkt1161.

93. Derafshi, B.H., Danko, T., Chanda, S., Batista, P.J., Litzenburger, U., Lee, Q.Y., Ng, Y.H., Sebin, A., Chang, H.Y., Südhof, T.C., et al. (2022). The autism risk factor CHD8 is a chromatin activator in human neurons and functionally dependent on the ERK-MAPK pathway effector ELK1. Sci. Rep. 12, 22425. 10.1038/s41598-022-23614-x.

94. Cong, L., Ran, F.A., Cox, D., Lin, S., Barretto, R., Habib, N., Hsu, P.D., Wu, X., Jiang, W., Marraffini, L.A., et al. (2013). Multiplex Genome Engineering Using CRISPR/Cas Systems. Science 339, 819–823. 10.1126/science.1231143.

95. McFarlane, L., Truong, V., Palmer, J.S., and Wilhelm, D. (2013). Novel PCR Assay for Determining the Genetic Sex of Mice. Sex. Dev. 7, 207–211. 10.1159/000348677.

96. Zheng, G.X.Y., Terry, J.M., Belgrader, P., Ryvkin, P., Bent, Z.W., Wilson, R., Ziraldo, S.B., Wheeler, T.D., McDermott, G.P., Zhu, J., et al. (2017). Massively parallel digital transcriptional profiling of single cells. Nat Commun 8, 14049. 10.1038/ncomms14049.

97. Dobin, A., Davis, C.A., Schlesinger, F., Drenkow, J., Zaleski, C., Jha, S., Batut, P., Chaisson, M., and Gingeras, T.R. (2013). STAR: ultrafast universal RNA-seq aligner. Bioinformatics 29, 15–21. 10.1093/bioinformatics/bts635.

98. Danecek, P., Bonfield, J.K., Liddle, J., Marshall, J., Ohan, V., Pollard, M.O., Whitwham, A., Keane, T., McCarthy, S.A., Davies, R.M., et al. (2021). Twelve years of SAMtools and BCFtools. Gigascience 10. 10.1093/gigascience/giab008.

99. Kaminow, B., Yunusov, D., and Dobin, A. (2021). STARsolo: accurate, fast and versatile mapping/quantification of single-cell and single-nucleus RNA-seq data. bioRxiv, 2021.05.05.442755. 10.1101/2021.05.05.442755.

100. Lun, A.T.L., Riesenfeld, S., Andrews, T., Dao, T.P., Gomes, T., Marioni, J.C., and Jamboree, participants in the 1st H.C.A. (2019). EmptyDrops: distinguishing cells from empty droplets in droplet-based single-cell RNA sequencing data. 20, 63. 10.1186/s13059-019-1662-y.

101. Zeisel, A., Hochgerner, H., Lönnerberg, P., Johnsson, A., Memic, F., Zwan, J. van der, Häring, M., Braun, E., Borm, L.E., Manno, G.L., et al. (2018). Molecular Architecture of the Mouse Nervous System. 174, 999–1014.e22. 10.1016/j.cell.2018.06.021.

102. Hao, Y., Hao, S., Andersen-Nissen, E., Mauck, W.M., Zheng, S., Butler, A., Lee, M.J., Wilk, A.J., Darby, C., Zager, M., et al. (2021). Integrated analysis of multimodal single-cell data. Cell 184, 3573–3587.e29. 10.1016/j.cell.2021.04.048.

103. Tirosh, I., Izar, B., Prakadan, S.M., II, M.H.W., Treacy, D., Trombetta, J.J., Rotem, A., Rodman, C., Lian, C., Murphy, G., et al. (2016). Dissecting the multicellular ecosystem of metastatic melanoma by single-cell RNA-seq. Science 352, 189–196. 10.1126/science.aad0501.

104. Durinck, S., Spellman, P.T., Birney, E., and Huber, W. (2009). Mapping identifiers for the integration of genomic datasets with the R/Bioconductor package biomaRt. Nat Protoc 4, 1184– 1191. 10.1038/nprot.2009.97.

105. Hafemeister, C., and Satija, R. (2019). Normalization and variance stabilization of single-cell RNA-seq data using regularized negative binomial regression. Genome Biol 20, 296. 10.1186/s13059-019-1874-1.

106. Waltman, L., and Eck, N.J. van (2013). A smart local moving algorithm for large-scale modularity-based community detection. Arxiv. 10.48550/arxiv.1308.6604.

107. Germain, P.-L., Lun, A., Meixide, C.G., Macnair, W., and Robinson, M.D. (2021). Doublet identification in single-cell sequencing data using scDblFinder. F1000Res 10, 979. 10.12688/f1000research.73600.2.

108. Griffiths, J.A., Richard, A.C., Bach, K., Lun, A.T.L., and Marioni, J.C. (2018). Detection and removal of barcode swapping in single-cell RNA-seq data. Nat Commun 9, 2667. 10.1038/s41467-018-05083-x.

109. Lin, J., Keogh, E., Wei, L., and Lonardi, S. (2007). Experiencing SAX: a novel symbolic representation of time series. Data Min. Knowl. Discov. 15, 107–144. 10.1007/s10618-007-0064-z.

110. Senin, P., and Malinchik, S. (2013). SAX-VSM: Interpretable Time Series Classification using SAX and Vector Space Model. 2013 Ieee 13th Int Conf Data Min, 1175–1180. 10.1109/icdm.2013.52.

111. Bar-Joseph, Z., Gifford, D.K., and Jaakkola, T.S. (2001). Fast optimal leaf ordering for hierarchical clustering. Bioinformatics 17, S22–S29. 10.1093/bioinformatics/17.suppl_1.s22.

112. Raudvere, U., Kolberg, L., Kuzmin, I., Arak, T., Adler, P., Peterson, H., and Vilo, J. (2019). g:Profiler: a web server for functional enrichment analysis and conversions of gene lists (2019 update). Nucleic Acids Res 47, W191–W198. 10.1093/nar/gkz369.

113. Kolberg, L., Raudvere, U., Kuzmin, I., Vilo, J., and Peterson, H. (2020). gprofiler2 -- an R package for gene list functional enrichment analysis and namespace conversion toolset g:Profiler. F1000Research 9, ELIXIR-709. 10.12688/f1000research.24956.2.

114. Supek, F., Bošnjak, M., Škunca, N., and Šmuc, T. (2011). REVIGO Summarizes and Visualizes Long Lists of Gene Ontology Terms. Plos One 6, e21800. 10.1371/journal.pone.0021800.

115. Bolger, A.M., Lohse, M., and Usadel, B. (2014). Trimmomatic: a flexible trimmer for Illumina sequence data. Bioinformatics 30, 2114–2120. 10.1093/bioinformatics/btu170.

116. Liao, Y., Smyth, G.K., and Shi, W. (2014). featureCounts: an efficient general purpose program for assigning sequence reads to genomic features. Bioinformatics 30, 923–930. 10.1093/bioinformatics/btt656.

117. Love, M.I., Huber, W., and Anders, S. (2014). Moderated estimation of fold change and dispersion for RNA-seq data with DESeq2. Genome Biol 15, 550. 10.1186/s13059-014-0550-8.

118. Zhu, A., Ibrahim, J.G., and Love, M.I. (2019). Heavy-tailed prior distributions for sequence count data: removing the noise and preserving large differences. 35, 2084–2092. 10.1093/bioinformatics/bty895.

