## Supplementary_Figures for "Cell type-specific dysregulation of gene expression due to *Chd8* haploinsufficiency during mouse cortical development"

#### Supplemental Figures and Legends

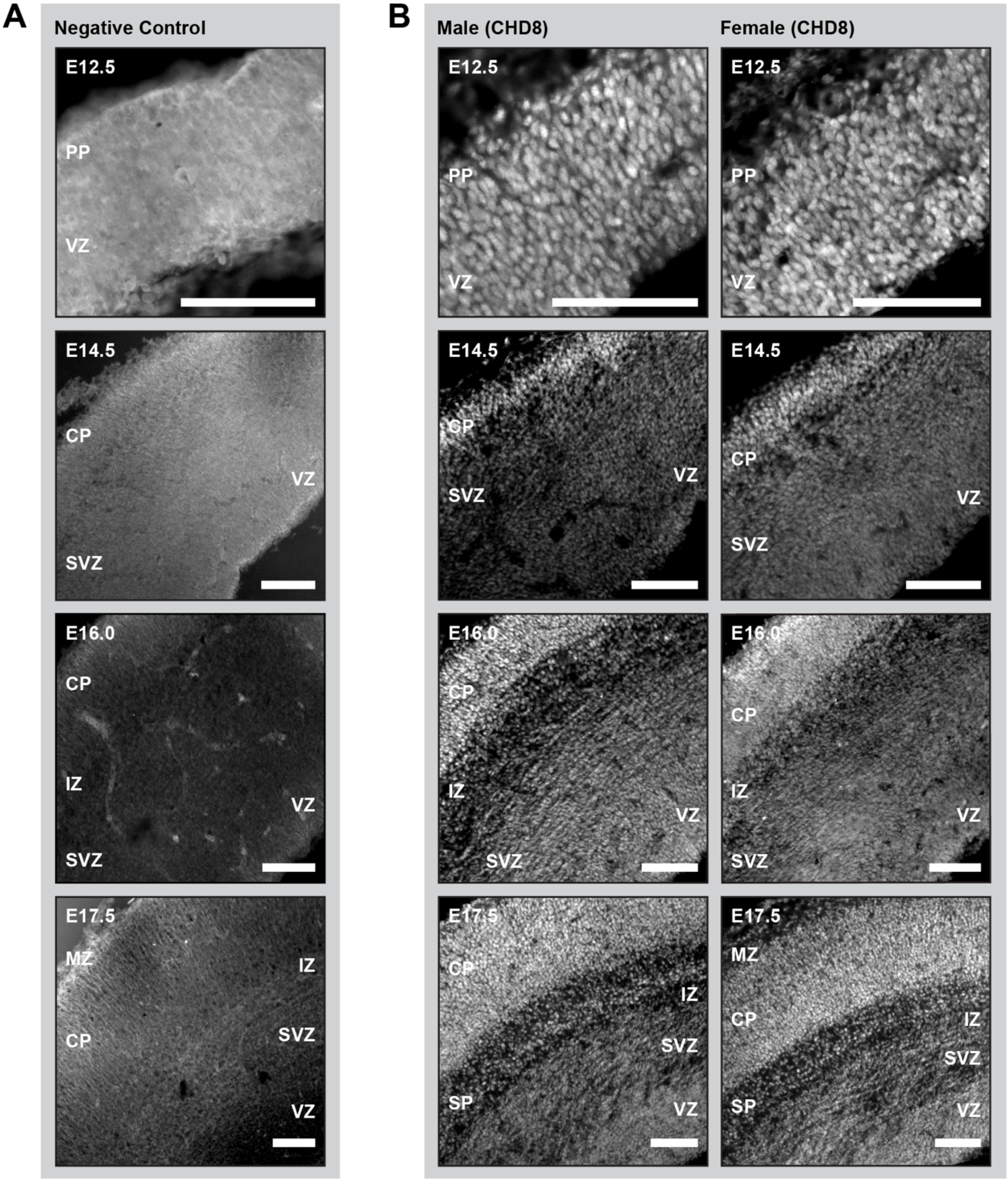

**Figure S1. Validation of CHD8 immunostaining in coronal sections of the wild type embryonic cortex.** (**A**) Results of negative immunostaining controls, using only anti-rabbit secondary antibody in cortical sections from embryonic day (E) 12.5, E14.5, E16.0, and E17.5 wild type embryos. (**B**) Immunostaining for CHD8 (white) in litter-matched male (*left*) and female (*right*) wild type cortical sections. See also Figure 1 and Figure S2. Scale bars: 100µm. VZ = ventricular zone; SVZ = subventricular zone; IZ = intermediate zone; PP = preplate; SP = subplate; CP = cortical plate; MZ = marginal zone.

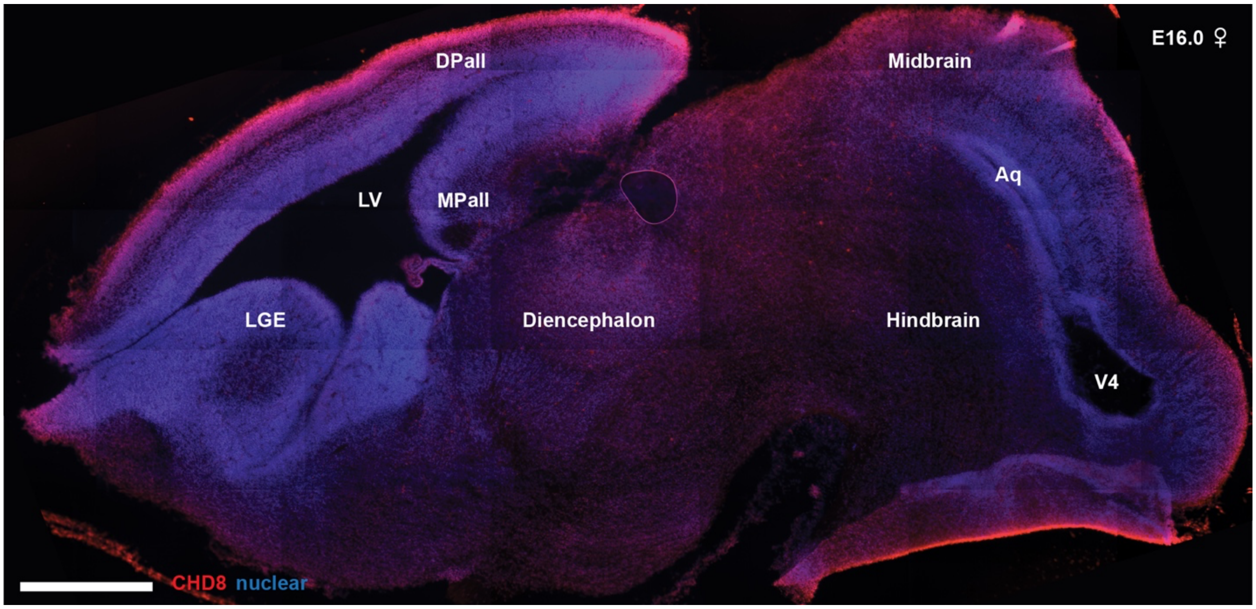


**Figure S2. Immunohistochemistry for CHD8 along the rostrocaudal axis in a sagittal section of the embryonic day (E) 16.0 wild type female brain.** See also Figure 1 and Figure S1. Scale bar: 500µm. Red = CHD8; blue = Hoechst 33342. Dpall = dorsal pallium; LV = lateral ventricle; MPall = medial pallium; LGE = lateral ganglionic eminence; Aq = aqueduct of Sylvius; V4 = fourth ventricle.

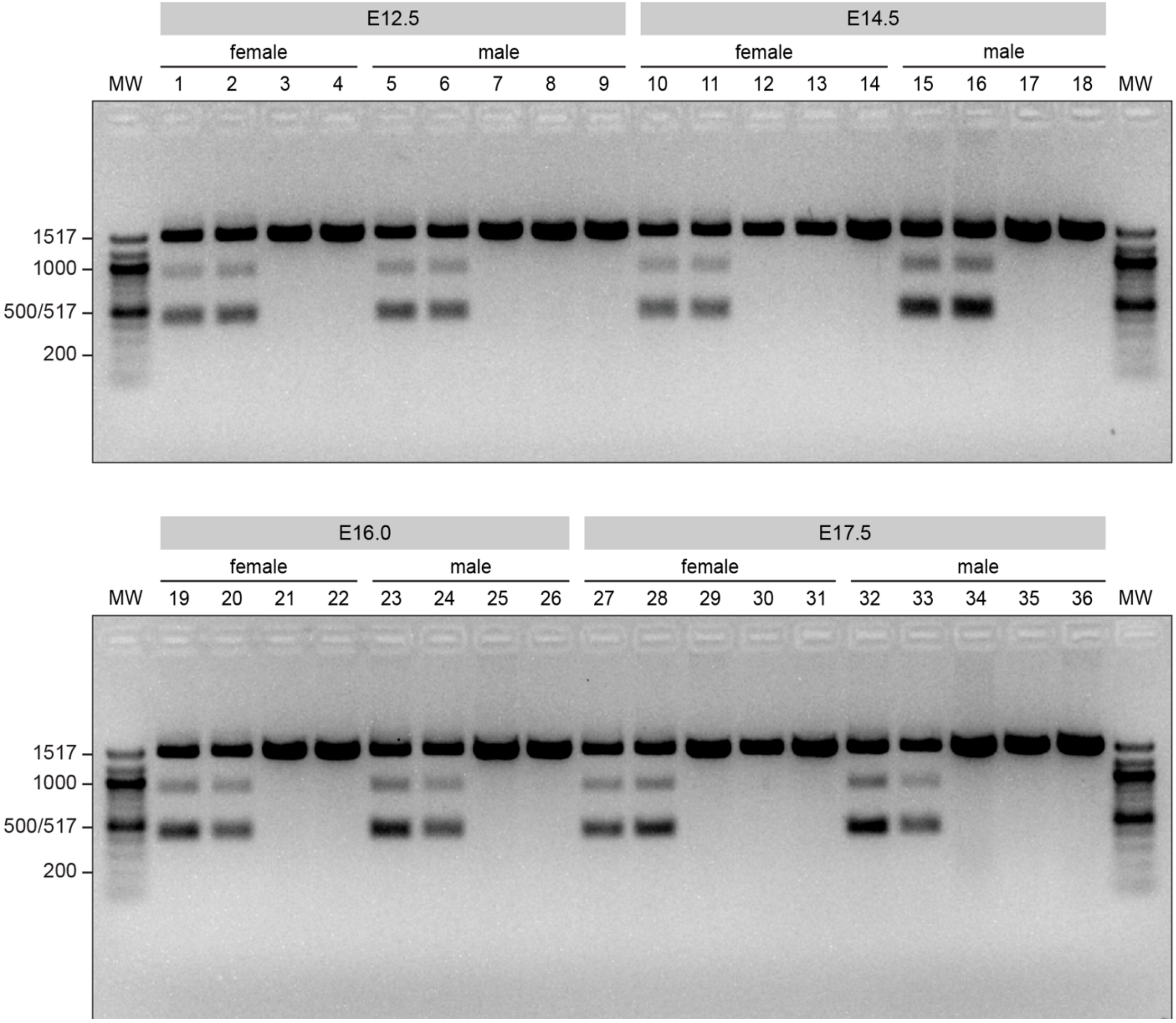


**Figure S3. Genotyping results for wild type and *Chd8*$^{+/-}$ embryos harvested for single-cell RNA-sequencing.** Agarose gel image showing PCR results using *Chd8* genotyping primers (5'-AACAGGCTGTCTCATGGGAA-3' and 5'-AAGCCACACTGCCTTGAAAG-3') across all embryos used for single-cell RNA-sequencing experiments (Methods). Homozygous *Chd8*$^{+/+}$ (wild type) mice exhibited the expected 1495bp PCR product, while *Chd8*$^{+/-}$ mice also displayed a 434bp product corresponding to the disrupted *Chd8* allele (see Figure 1A, Methods). Embryo samples are numbered by lane (1-36); corresponding embryo sample IDs are listed in Table S2. Cortical samples from embryos genotyped in lanes 8, 13, 30, and 35 were excluded from scRNA-seq analysis due to redundancy with litter- and sex-matched sample pairs included in our analysis (Methods). See also Figures 1-2 and Table S2. E = embryonic day; MW = molecular weight.

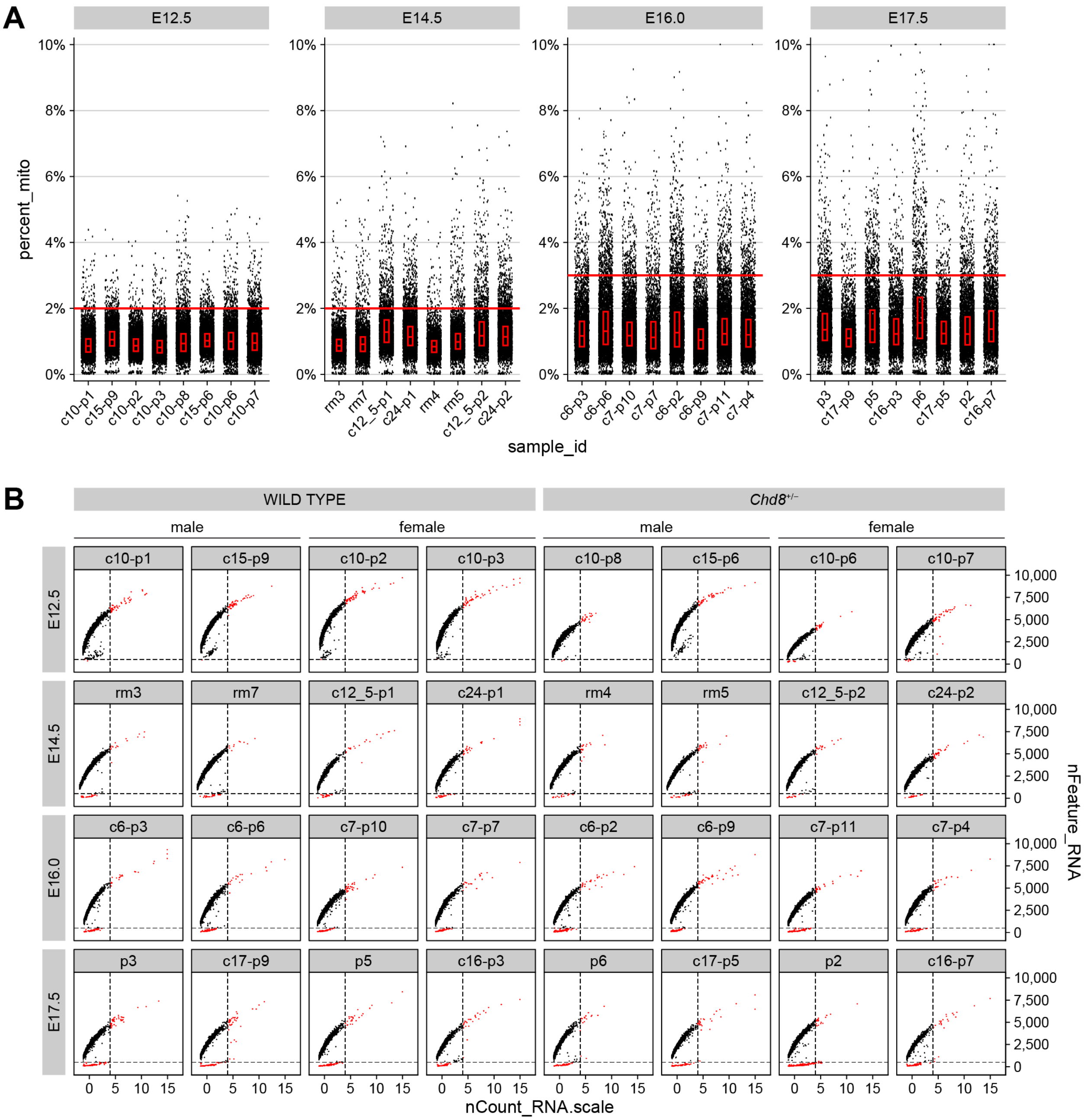


**Figure S4. Filtering cells by counts, genes detected, and percent mitochondrial content for the embryonic single-cell RNA-sequencing data.** (**A**) Percent mitochondrial content (percent_mito) per cell for each sample, with red box plots indicating median and interquartile range. Red lines indicate thresholds used for cell filtering: we removed cells with >2% mitochondrial reads in embryonic day (E) 12.5 and E14.5 samples, and we removed cells with >3% mitochondrial reads in E16.0 and E17.5 samples (Methods). (**B**) Z-scored total feature counts per cell (nCount_RNA.scale) and total genes detected per cell (nFeature_RNA) plotted for each sample. Dashed lines indicate thresholds used for cell filtering: we removed cells with nCount_RNA.scale > 4 and cells with nFeature_RNA < 500 (Methods). Cells are color-coded based on whether they were removed (red) or retained (black). See also Figure 2 and Table S2.

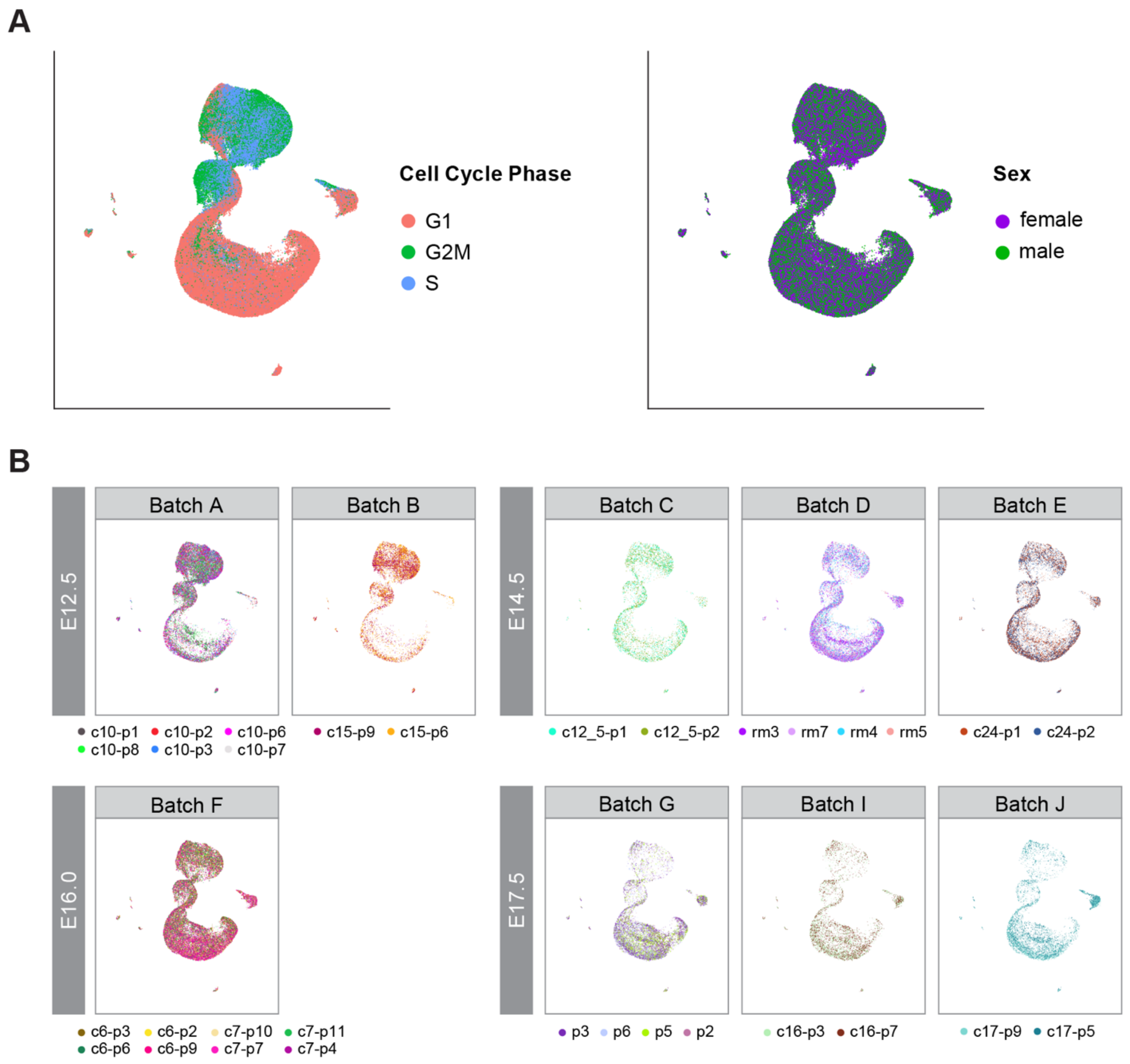


**Figure S5. UMAP embedding of the embryonic single-cell RNA-sequencing data colored by cell cycle phase, sex, and sample ID.** (**A**) UMAP embedding of 135,926 cells colored by cell cycle phase (*left*), inferred by expression of S phase and G2/M phase markers using Seurat's "CellCycleScoring" function, and sex (*right*) (Methods). (**B**) UMAP embedding of each batch plotted separately with cells colored by sample ID. See also Figure 2 and Table S2. E = embryonic day.

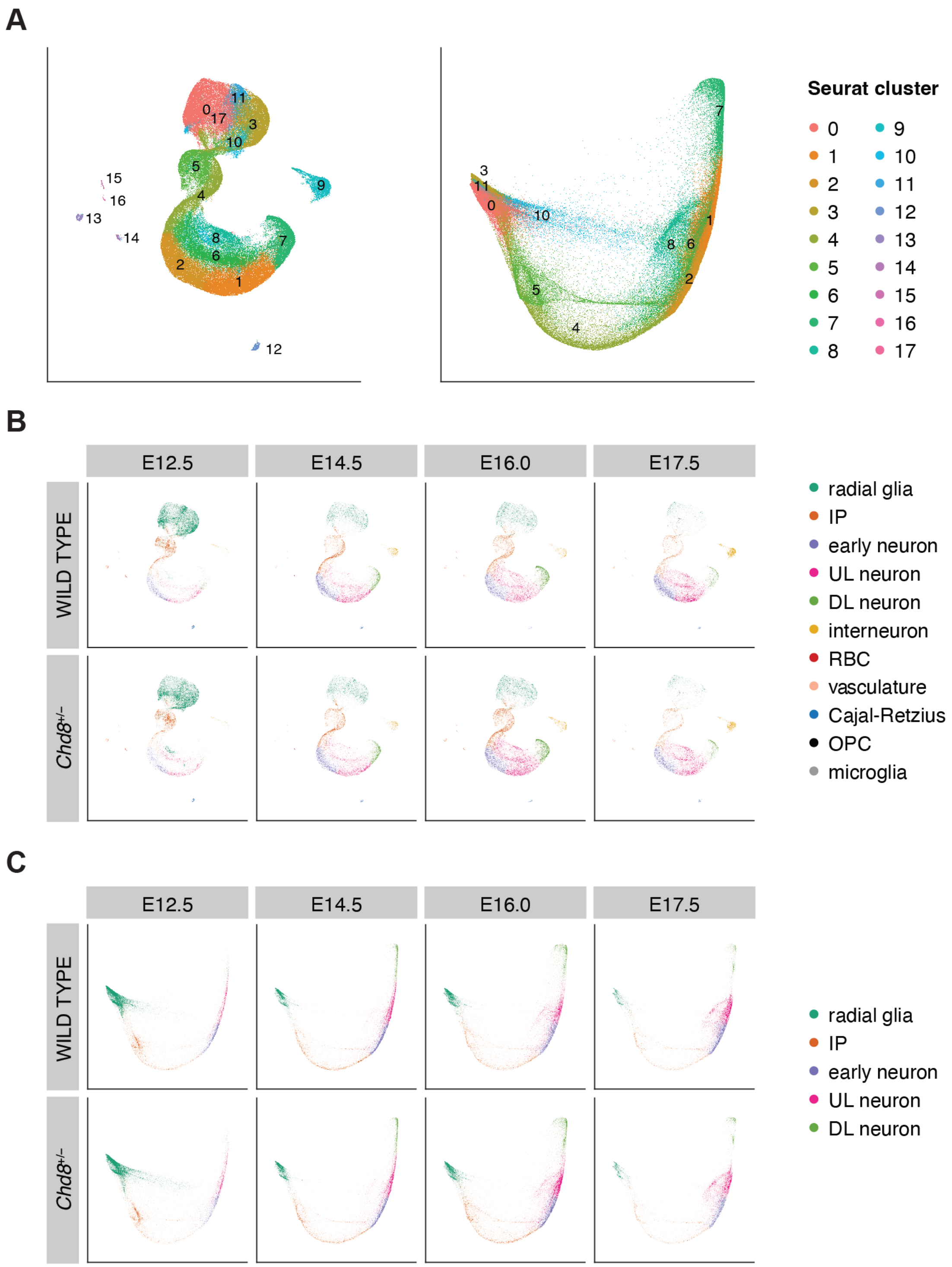


**Figure S6. UMAP and PHATE embeddings of the embryonic single-cell RNA-sequencing data colored by initial clustering results and cell type assignments.** (**A**) UMAP (*left*) and PHATE (*right*) embeddings colored by initial clustering results (Seurat cluster; Methods). (**B-C**) UMAP (**B**) and PHATE (**C**) embeddings colored by cell type assignment, plotted separately by time point and genotype. See also Figure 2, Figures S7-S8, and Tables S3-S4. E = embryonic day; IP = intermediate progenitors; UL = upper-layer cortical neurons; DL = deep-layer cortical neurons; RBC = red blood cells; OPC = oligodendrocyte precursor cells.

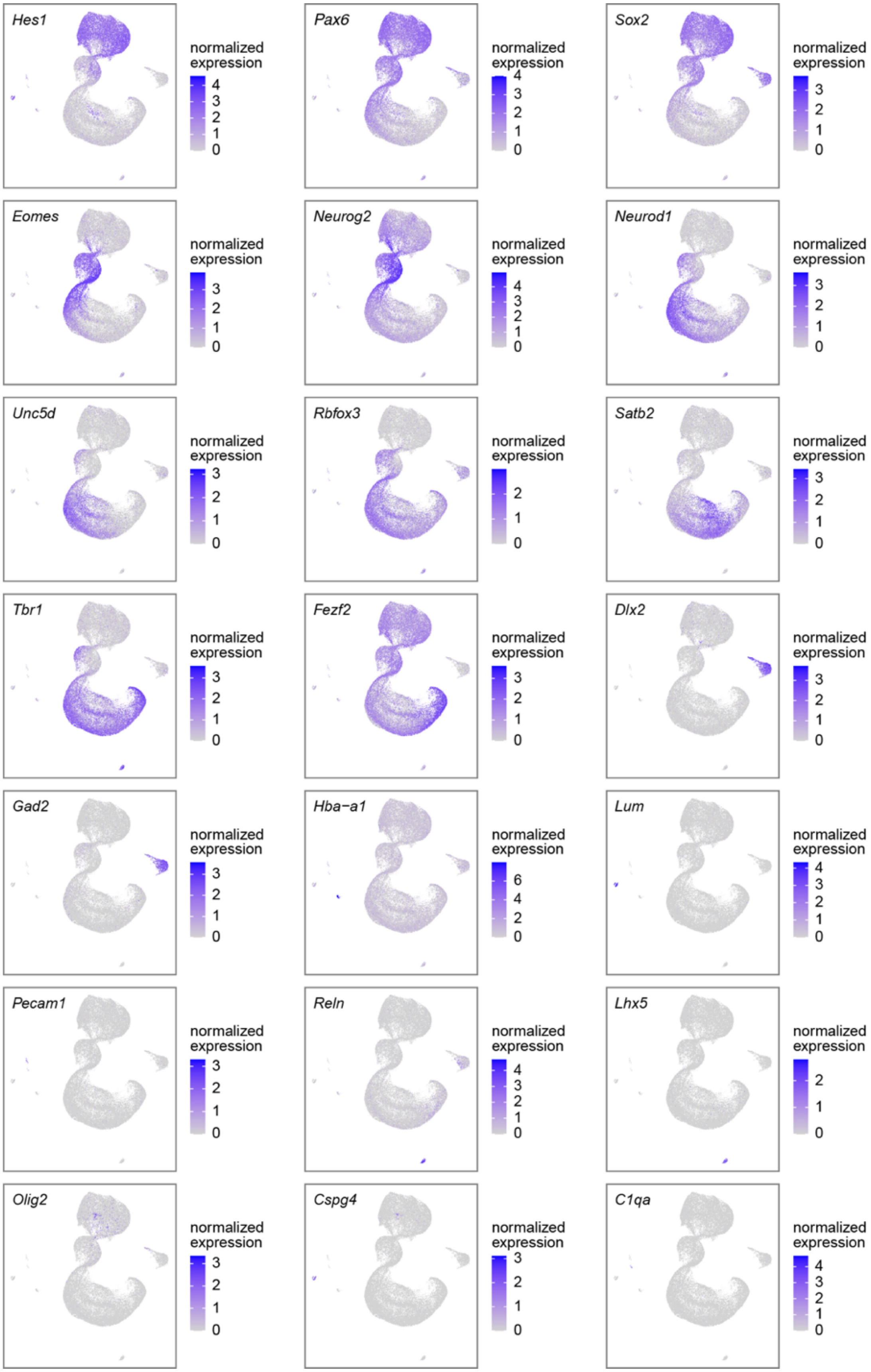
Hes1
normalized expression
4 3 2 1 0
Pax6
normalized expression
4 3 2 1 0
Sox2
normalized expression
3 2 1 0
Eomes
normalized expression
3 2 1 0
Neurog2
normalized expression
4 3 2 1 0
Neurod1
normalized expression
3 2 1 0
Unc5d
normalized expression
3 2 1 0
Rbfox3
normalized expression
2 1 0
Satb2
normalized expression
3 2 1 0
Tbr1
normalized expression
3 2 1 0
Fezf2
normalized expression
3 2 1 0
Dlx2
normalized expression
3 2 1 0
Gad2
normalized expression
3 2 1 0
Hba−a1
normalized expression
6 4 2 0
Lum
normalized expression
4 3 2 1 0
Pecam1
normalized expression
3 2 1 0
Reln
normalized expression
4 3 2 1 0
Lhx5
normalized expression
2 1 0
Olig2
normalized expression
3 2 1 0
Cspg4
normalized expression
3 2 1 0
C1qa
normalized expression
4 3 2 1 0

**Figure S7. UMAP embedding of the embryonic single-cell RNA-sequencing data colored by normalized expression of cell type marker genes.** Marker genes include those specific to early progenitor cells (*Sox2*), radial glia (*Pax6*, *Hes1*), intermediate progenitors (*Eomes*, *Neurog2*), early neurons (*Neurod1*, *Unc5d*), neurons (*Rbfox3*), upper-layer neurons (*Satb2*), deep-layer neurons (*Tbr1*, *Fezf2*), the inhibitory neuron lineage (*Dlx2*), interneurons (*Gad2*), red blood cells (*Hba-a1*), vasculature (*Lum*, *Pecam1*), Cajal-Retzius cells (*Reln*, *Lhx5*), oligodendrocyte precursor cells (*Olig2*, *Cspg4*), and microglia (*C1qa*; Methods). See also Figure 2, Figure S8, and Table S3.

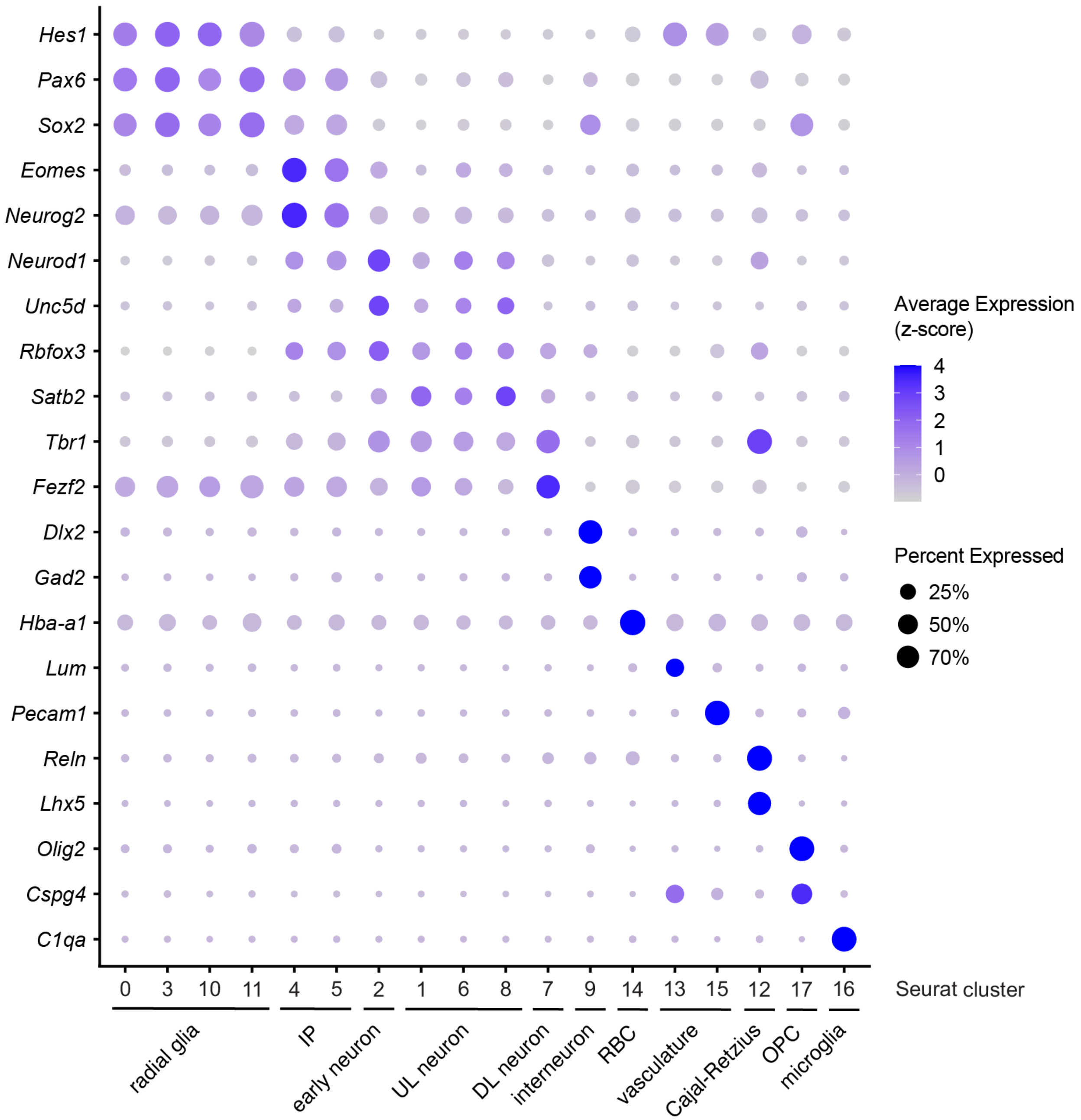


**Figure S8. Expression of cell type marker genes within each cluster in the embryonic single-cell RNA-sequencing dataset.** Dot plot of select cell type marker genes highly expressed in each cluster, using initial clustering results (Seurat cluster; Methods). Circle color corresponds to average expression within each cluster, z-scored per gene. Circle size corresponds to the percentage of cells in a cluster in which a given gene is detected. See also Figure 2, Figure S7, and Tables S3-S4. IP = intermediate progenitors; UL = upper-layer cortical neurons; DL = deep-layer cortical neurons; RBC = red blood cells; OPC = oligodendrocyte precursor cells.

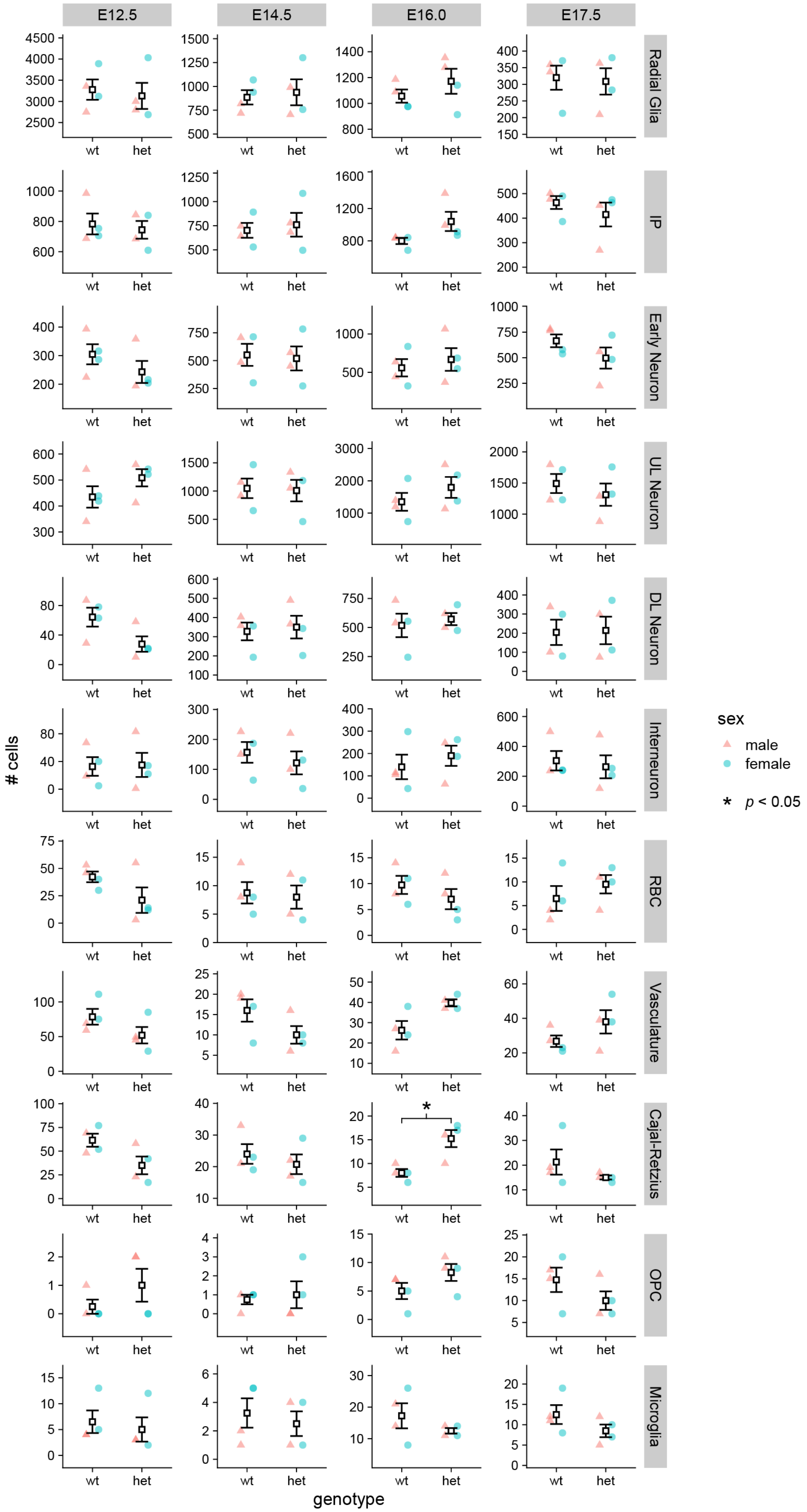

E12.5
E14.5
E16.0
E17.5
Radial Glia
IP
Early Neuron
UL Neuron
DL Neuron
Interneuron
RBC
Vasculature
Cajal-Retzius
OPC
Microglia
### cells
genotype
wt
het
sex
male
female
* p < 0.05

**Figure S9. Comparison of cell type representation between the wild type and *Chd8*$^{+/-}$ embryonic cortex single-cell RNA-sequencing data.** Comparison of the number of cells within each cluster between the wild type (wt) and *Chd8*$^{+/-}$ (het) embryonic mouse cortex, separated by time point. Data are represented as mean ± SEM, with datapoints for individual replicates color-coded by sex. *P* values were calculated by two-tailed Welch's *t*-test (Methods). See also Figure 2, Figure S8, and Tables S4-S5. E = embryonic day; RG = radial glia; IP = intermediate progenitors; UL = upper-layer; DL = deep-layer.

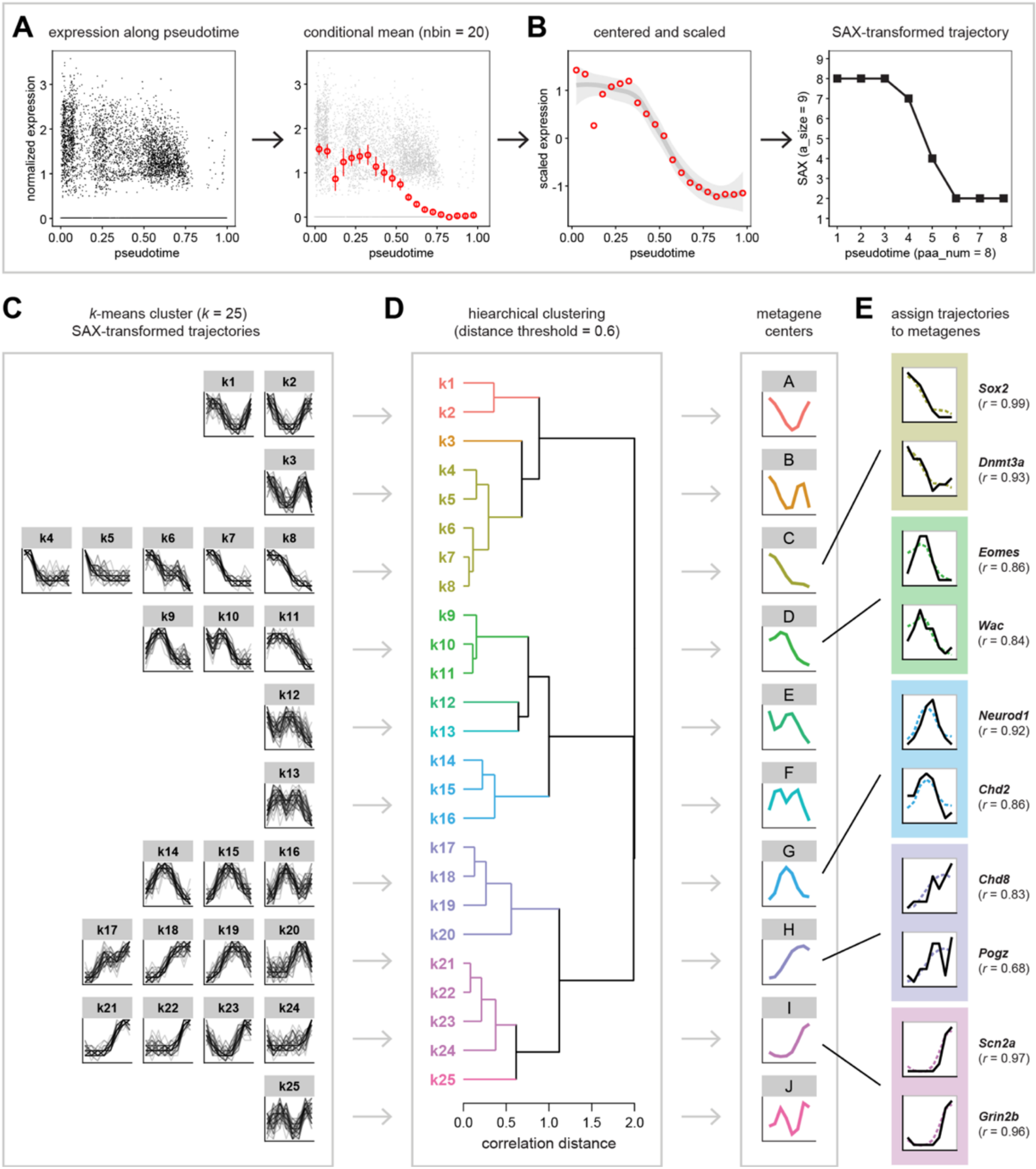
A
expression along pseudotime
normalized expression
pseudotime
conditional mean (nbin = 20)
B
centered and scaled
scaled expression
SAX-transformed trajectory
SAX (a_size = 9)
pseudotime (paa_num = 8)
C
k-means cluster (k = 25)
SAX-transformed trajectories
D
hiearchical clustering
(distance threshold = 0.6)
correlation distance
metagene
centers
E
assign trajectories
to metagenes
Sox2
(r = 0.99)
Dnmt3a
(r = 0.93)
Eomes
(r = 0.86)
Wac
(r = 0.84)
Neurod1
(r = 0.92)
Chd2
(r = 0.86)
Chd8
(r = 0.83)
Pogz
(r = 0.68)
Scn2a
(r = 0.97)
Grin2b
(r = 0.96)

**Figure S10. Inferring gene expression trajectories and defining metagenes from single-cell RNA-sequencing data.** (**A**) Schematic showing our method of gene expression trajectory definition, which involved aggregation of male and female samples for each time point and genotype, division of primary trajectory cells into 20 equally-spaced bins along pseudotime, and calculation of the average expression per bin per gene (Methods). (**B**) Schematic of gene expression trajectory centering, scaling, and transformation using symbolic aggregate approximation (SAX; Methods). (**C**) Schematic of our clustering approach, in which *k*-means clustering ($k$ = 25) was performed on SAX-transformed gene expression trajectories from embryonic day (E) 14.5, E16.0, and E17.5 wild type mouse cortex data (Methods). (**D**) Schematic of metagene identification from the clusters of trajectories from (**C**), which were aggregated into 10 metagene centers by hierarchical clustering using correlation distance and complete linkage (Methods). (**E**) Examples of gene expression trajectory assignment to metagenes identified in (**D**), based on highest Pearson correlation (*r*) between the SAX-transformed trajectory and the metagene center (Methods). The metagene center is plotted as dashed line, and the SAX-transformed gene expression trajectory is plotted as a solid line. See also Figures 3-4, Figures S11-S12, Table S6, and Table S10**.**

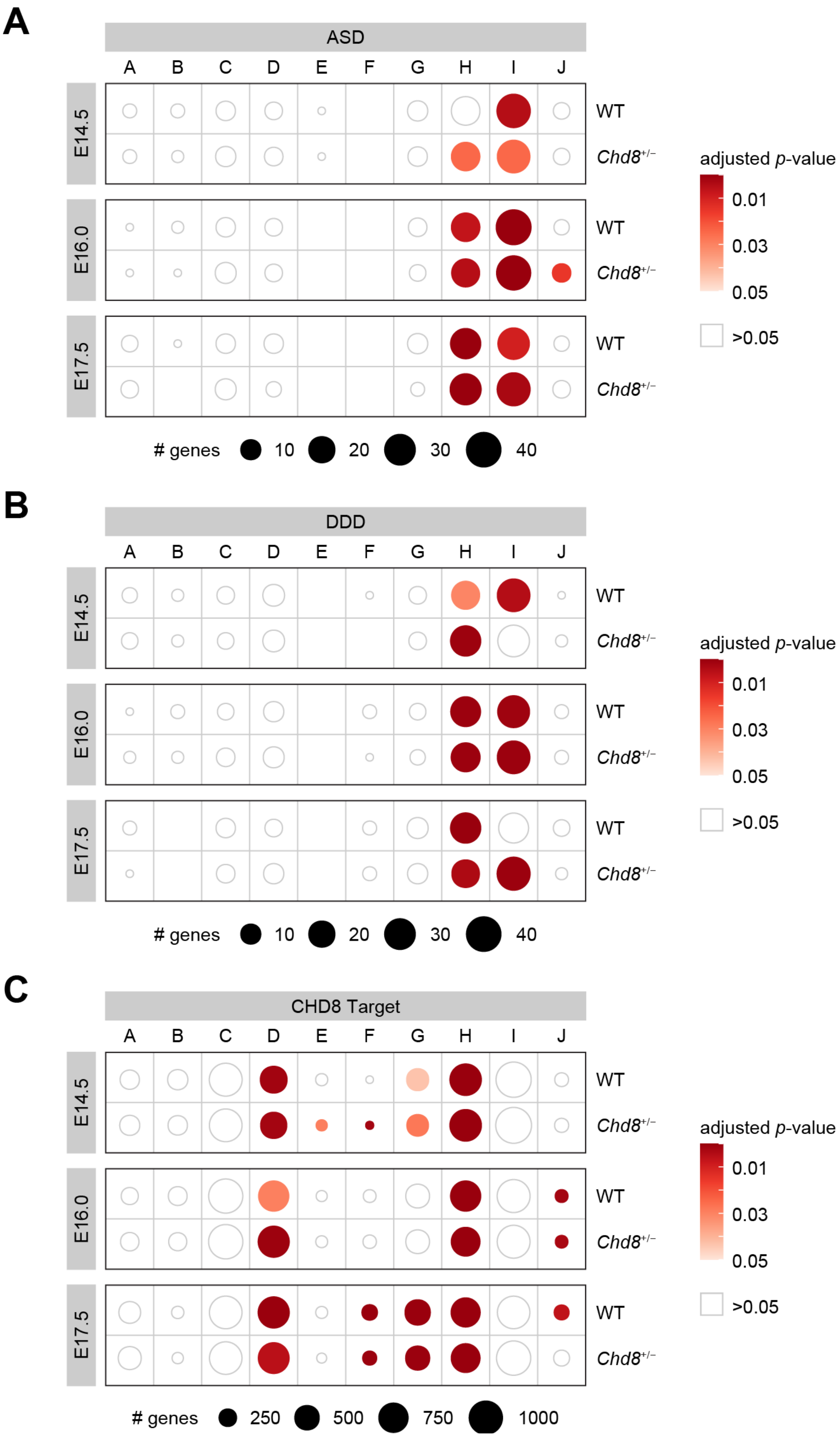
A
ASD
A B C D E F G H I J
E14.5
E16.0
E17.5
WT
Chd8+/−
adjusted p-value
0.01
0.03
0.05
>0.05
### genes 10 20 30 40
B
DDD
A B C D E F G H I J
E14.5
E16.0
E17.5
WT
Chd8+/−
adjusted p-value
0.01
0.03
0.05
>0.05
### genes 10 20 30 40
C
CHD8 Target
A B C D E F G H I J
E14.5
E16.0
E17.5
WT
Chd8+/−
adjusted p-value
0.01
0.03
0.05
>0.05
### genes 250 500 750 1000

**Figure S11. Metagene enrichment for ASD risk-associated genes, DDD genes, and CHD8 binding targets in the embryonic mouse cortex single-cell RNA-sequencing data.** Metagene enrichment for (**A**) autism spectrum disorder risk-associated genes (ASD), (**B**) Deciphering Developmental Disorders gene set (DDD), and (**C**) CHD8 binding target genes in the embryonic mouse cortex. Circle size corresponds to the number of genes within each metagene at each time point and genotype. Circle color corresponds to Benjamini Hochberg-adjusted *p*-value from one-tailed Fisher exact test (Methods). E = embryonic day; WT = wild type. See also Figure 4, Figure S10, and Tables S8-S9.

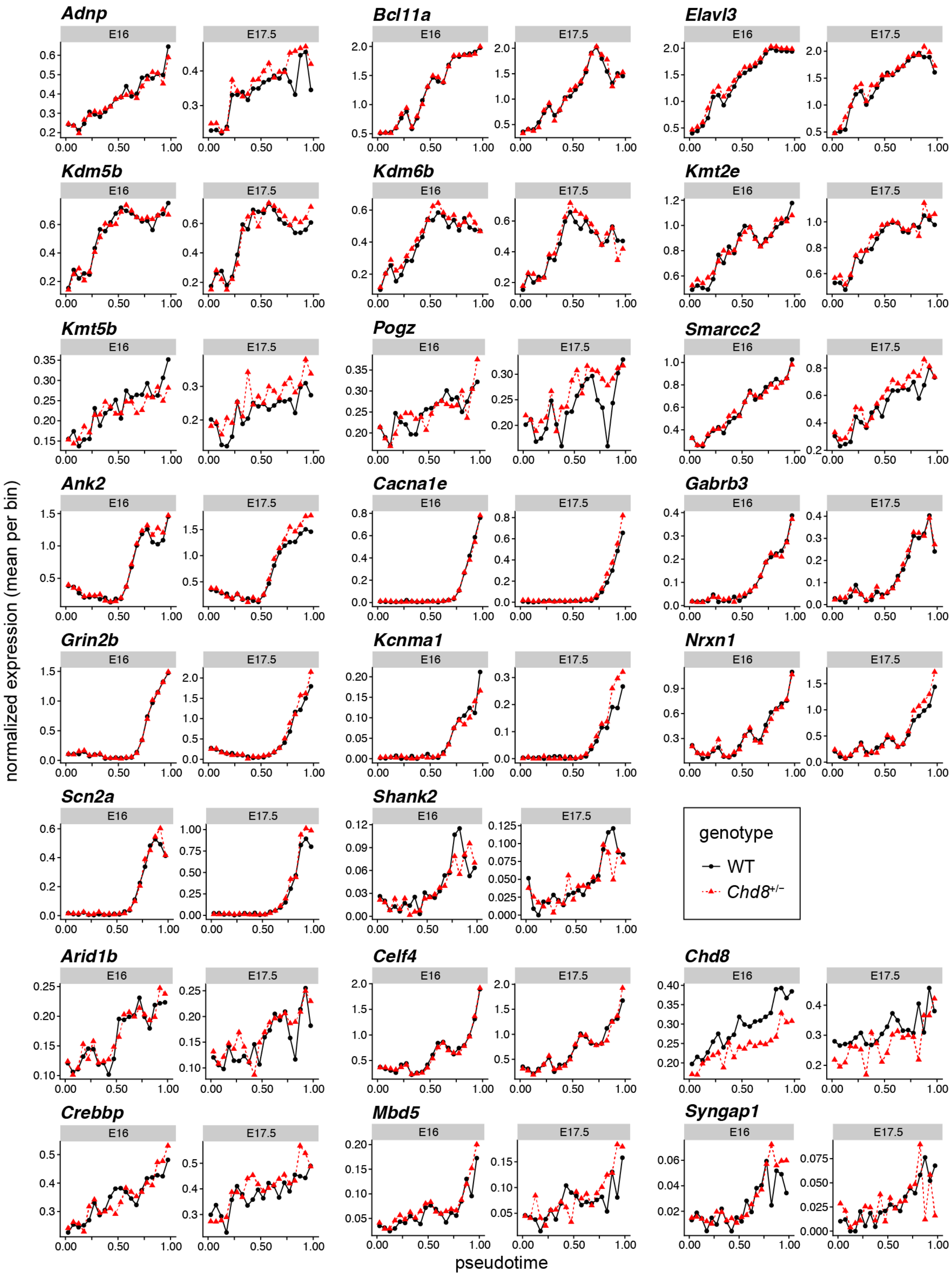

Adnp
Bcl11a
Elavl3
Kdm5b
Kdm6b
Kmt2e
Kmt5b
Pogz
Smarcc2
Ank2
Cacna1e
Gabrb3
Grin2b
Kcnma1
Nrxn1
Scn2a
Shank2
Arid1b
Celf4
Chd8
Crebbp
Mbd5
Syngap1
E16
E17.5
normalized expression (mean per bin)
pseudotime
genotype
WT
Chd8+/-

**Figure S12. Expression trajectories of autism spectrum disorder risk-associated genes in the embryonic day (E) 16.0 and E17.5 wild type (WT) and *Chd8*$^{+/-}$ cortex.** Gene expression trajectories inferred for select autism spectrum disorder risk-associated genes in metagenes H and I at E16.0 and E17.5 in the wild type (black solid line, circles) and *Chd8*$^{+/-}$ (red dotted line, triangles) mouse cortex. Mean expression per bin is plotted against pseudotime (Methods). See also Figure 4, Figure S10, Table S6, and Table S9.

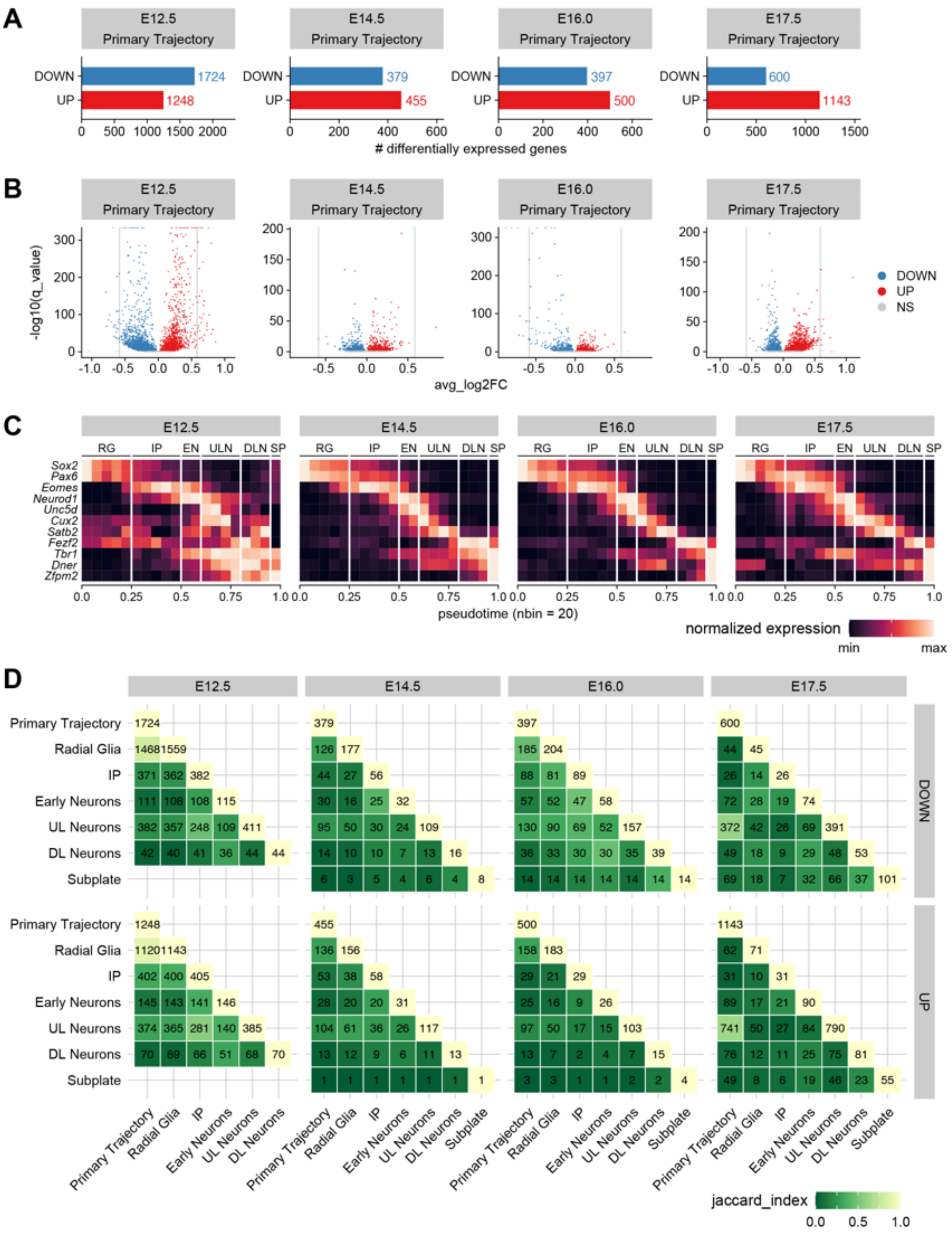

A
E12.5 Primary Trajectory
E14.5 Primary Trajectory
E16.0 Primary Trajectory
E17.5 Primary Trajectory
DOWN
UP
1724
1248
379
455
397
500
600
1143
### differentially expressed genes
B
-log10(q_value)
avg_log2FC
DOWN
UP
NS
C
E12.5
E14.5
E16.0
E17.5
RG IP EN ULN DLN SP
Sox2
Pax6
Eomes
Neurod1
Unc5d
Cux2
Satb2
Fezf2
Tbr1
Dner
Zfpm2
pseudotime (nbin = 20)
normalized expression
min
max
D
E12.5
E14.5
E16.0
E17.5
Primary Trajectory
Radial Glia
IP
Early Neurons
UL Neurons
DL Neurons
Subplate
DOWN
UP
jaccard_index
0.0 0.5 1.0

**Figure S13. Comparison of differential gene expression determined in the primary trajectory and in distinct cell types within the wild type and *Chd8*$^{+/-}$ embryonic mouse cortex single-cell RNA-sequencing data.** (**A**) Number of downregulated (blue; DOWN) and upregulated (red; UP) differentially expressed genes (DEGs) identified in the primary trajectory at each time point within the embryonic dataset, determined by Monocle 3 (Methods). (**B**) Volcano plots of Monocle 3 differential expression results for the primary trajectory at each time point, with genes color-coded by differential expression call (DOWN, blue; UP, red; not significantly different: NS, gray). Vertical gray lines = ±log2(1.5 fold-change). (**C**) Heat maps of normalized marker gene expression along pseudotime. For each time point, cells in the primary trajectory were divided into 20 bins, average gene expression was computed per bin, and then these values were min-max normalized per gene. Labels at top indicate cell type assignments by pseudotime partitions, determined by maximal marker gene expression (Methods). (**D**) Comparison of the lists of downregulated (DOWN, *top*) or upregulated (UP, *bottom*) DEGs identified from the primary trajectory or from individual cell types, separated by time point (Methods). Subplate cells in the embryonic day (E) 12.5 data were excluded, given the small number of cells (≤10 per genotype) identified at this time point. Each cell in the heat map is color-coded by the degree of similarity (jaccard_index) in DEG identity for a given comparison, while the number within each cell indicates the number of DEGs shared in that comparison. See also Figure 6, Figure S8, and Tables S14-S16. RG = radial glia; IP = intermediate progenitors; EN = early neurons; ULN = upper-layer (UL) cortical neurons; DLN = deep-layer (DL) cortical neurons; SP = subplate neurons.

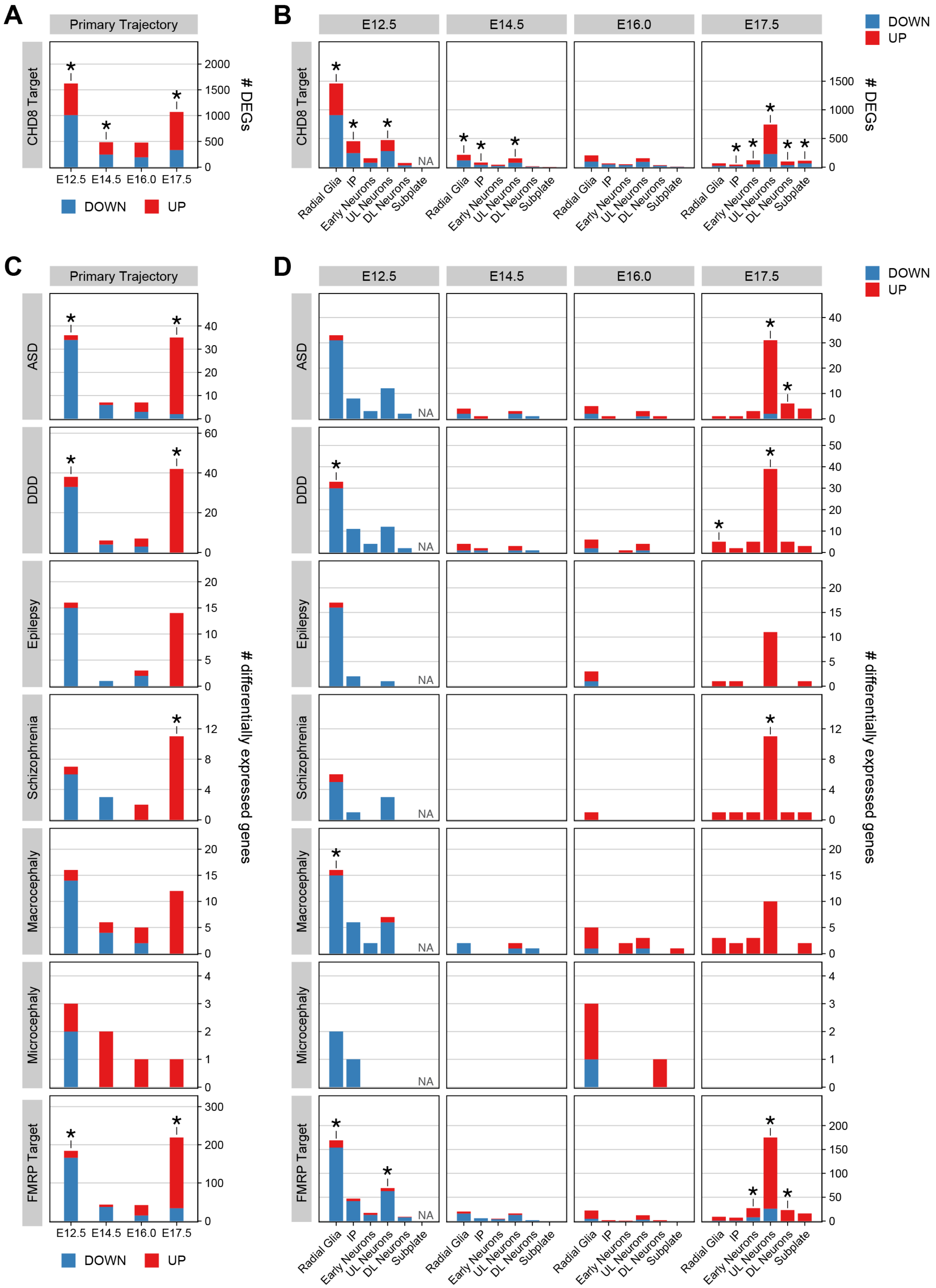
A
Primary Trajectory
CHD8 Target
### DEGs
0
500
1000
1500
2000
E12.5
E14.5
E16.0
E17.5
DOWN
UP
B
E12.5
E14.5
E16.0
E17.5
DOWN
UP
CHD8 Target
### DEGs
NA
Radial Glia
IP
Early Neurons
UL Neurons
DL Neurons
Subplate
C
Primary Trajectory
ASD
DDD
Epilepsy
Schizophrenia
Macrocephaly
Microcephaly
FMRP Target
### differentially expressed genes
E12.5
E14.5
E16.0
E17.5
DOWN
UP
D
E12.5
E14.5
E16.0
E17.5
DOWN
UP
ASD
DDD
Epilepsy
Schizophrenia
Macrocephaly
Microcephaly
FMRP Target
### differentially expressed genes
NA
Radial Glia
IP
Early Neurons
UL Neurons
DL Neurons
Subplate

**Figure S14. Enrichment of CHD8 target genes and neurodevelopmental disorder-associated genes among differentially expressed genes in the *Chd8*$^{+/-}$ embryonic cortex, separated by time point.** Bars are color-coded by the number of downregulated (DOWN; blue) and upregulated (UP; red) differentially expressed genes (DEGs) identified by Monocle 3 at each time point (Methods). (**A**-**B**) Intersection between CHD8 target genes and (**A**) DEGs identified in the primary trajectory or (**B**) DEGs identified in each cell type within the primary trajectory (Methods). (**C**-**D**) Intersection between neurodevelopmental disorder-associated genes or FMRP target genes and (**C**) DEGs identified in the primary trajectory or (**D**) DEGs identified in each cell type within the primary trajectory (Methods). Significance was determined by one-way Fisher's exact test, with adjustment for multiple testing (Methods); * = Benjamini Hochberg-adjusted $p$-value < 0.05. See also Figure 6, Figure S13, Table S9, and Tables S15-S17. E = embryonic day; RG = radial glia; IP = intermediate progenitors; UL = upper-layer; DL = deep-layer; ASD = autism spectrum disorder risk-associated genes, DDD = Deciphering Developmental Disorders gene set; NA = not applicable.

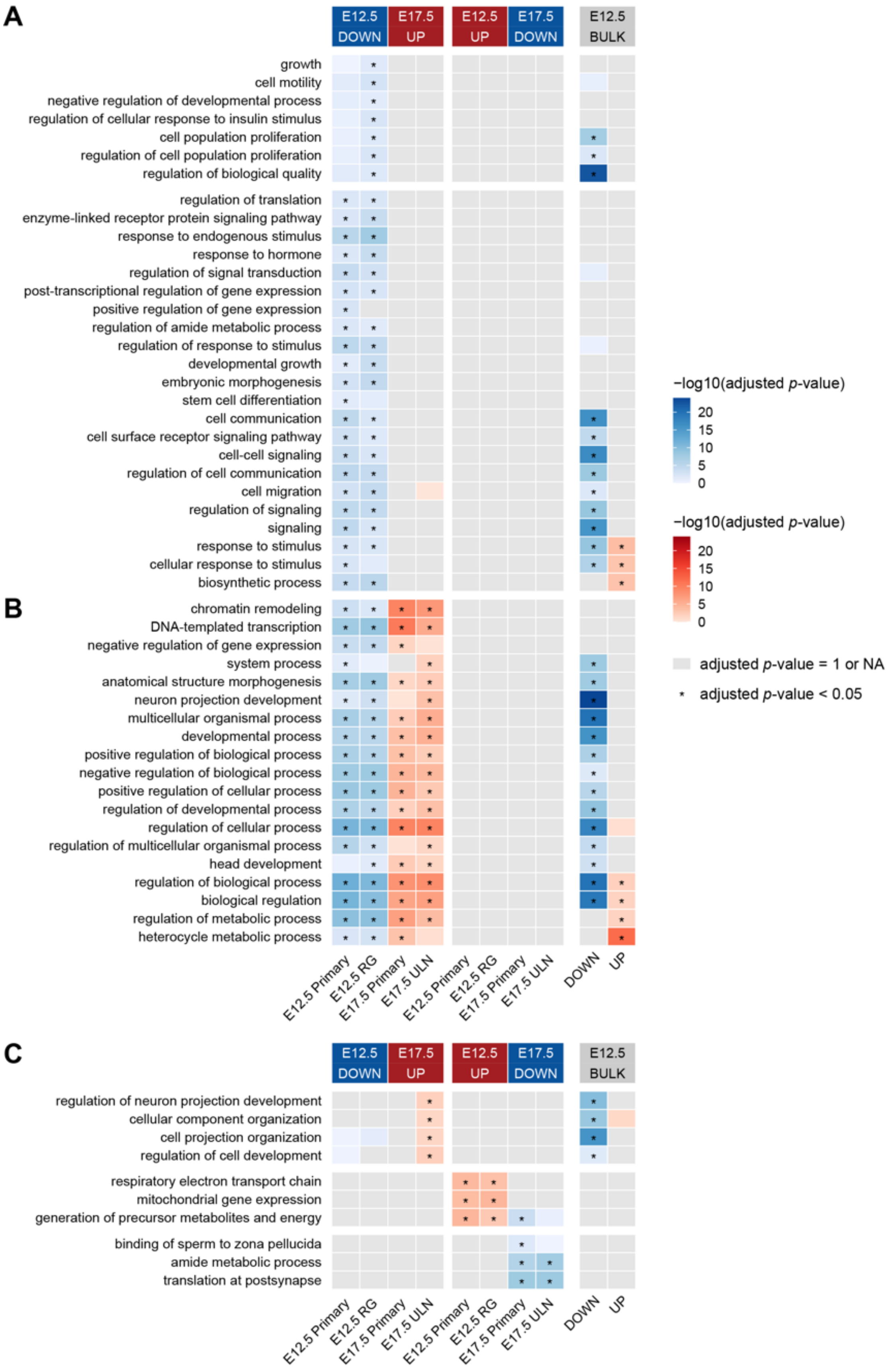

A
B
C
E12.5 DOWN
E17.5 UP
E12.5 UP
E17.5 DOWN
E12.5 BULK
growth
cell motility
negative regulation of developmental process
regulation of cellular response to insulin stimulus
cell population proliferation
regulation of cell population proliferation
regulation of biological quality
regulation of translation
enzyme-linked receptor protein signaling pathway
response to endogenous stimulus
response to hormone
regulation of signal transduction
post-transcriptional regulation of gene expression
positive regulation of gene expression
regulation of amide metabolic process
regulation of response to stimulus
developmental growth
embryonic morphogenesis
stem cell differentiation
cell communication
cell surface receptor signaling pathway
cell-cell signaling
regulation of cell communication
cell migration
regulation of signaling
signaling
response to stimulus
cellular response to stimulus
biosynthetic process
chromatin remodeling
DNA-templated transcription
negative regulation of gene expression
system process
anatomical structure morphogenesis
neuron projection development
multicellular organismal process
developmental process
positive regulation of biological process
negative regulation of biological process
positive regulation of cellular process
regulation of developmental process
regulation of cellular process
regulation of multicellular organismal process
head development
regulation of biological process
biological regulation
regulation of metabolic process
heterocycle metabolic process
E12.5 Primary
E12.5 RG
E17.5 Primary
E17.5 ULN
DOWN
UP
regulation of neuron projection development
cellular component organization
cell projection organization
regulation of cell development
respiratory electron transport chain
mitochondrial gene expression
generation of precursor metabolites and energy
binding of sperm to zona pellucida
amide metabolic process
translation at postsynapse
−log10(adjusted p-value)
20
15
10
5
0
adjusted p-value = 1 or NA
adjusted p-value < 0.05

**Figure S15. Enrichment of Gene Ontology Biological Process (GO:BP) terms among differentially expressed genes in the *Chd8*$^{+/-}$ embryonic day (E) 12.5 and E17.5 cortex.** (**A-B**) Heat maps showing enrichment of representative functional terms among downregulated genes (DOWN) identified in the primary trajectory (Primary) and radial glia (RG) at E12.5 that (**A**) were not enriched among upregulated genes (UP) at E17.5 or (**B**) were also enriched among upregulated genes in the primary trajectory and upper-layer cortical neurons (ULN) at E17.5. Enrichment among DOWN/UP genes in E12.5 bulk cortical data is shown at right. (**C**) Enrichment of representative functional terms among upregulated genes identified in ULN at E17.5 (*top*), upregulated genes in the primary trajectory and RG at E12.5 (*middle*), and downregulated genes in the primary trajectory and ULN at E17.5 (*bottom*). Enrichment among DOWN/UP genes in E12.5 bulk cortical data is shown (*right*). Differentially expressed genes were determined by Monocle 3 (for scRNA-seq data) or DESeq2 (for bulk RNA-seq data), and functional enrichment analysis was performed using g:Profiler and summarized using Revigo (Methods). No GO:BP terms were enriched among DOWN/UP genes in E17.5 bulk cortical data. Cells in the heat map are colored by significance of enrichment and direction of differential expression (blue color scale = DOWN DEG enrichment; red color scale = UP DEG enrichment; gray cells = adjusted $p$-value = 1 or NA (no genes intersect that term); * = g:SCS-adjusted $p$-value < 0.05) (Methods). See also Figure 6, Tables S15-S16, Table S19, and Tables S22-S23. E = embryonic day.

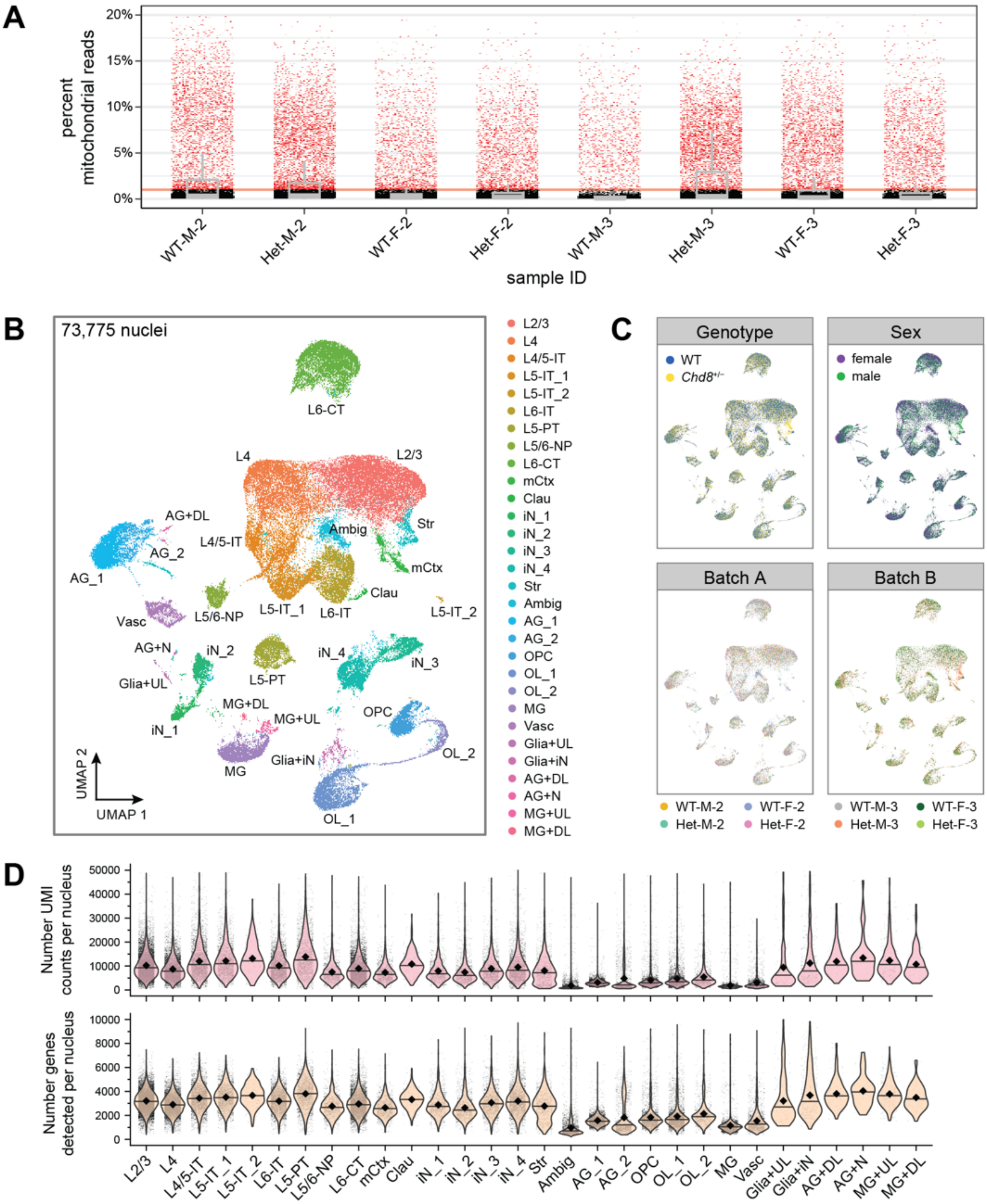
A
percent mitochondrial reads
20%
15%
10%
5%
0%
WT-M-2
Het-M-2
WT-F-2
Het-F-2
WT-M-3
Het-M-3
WT-F-3
Het-F-3
sample ID
B
73,775 nuclei
L6-CT
L4
L2/3
AG+DL
L4/5-IT
Ambig
Str
AG_2
mCtx
AG_1
Clau
Vasc
L5/6-NP
L5-IT_1
L6-IT
L5-IT_2
AG+N
iN_2
iN_4
iN_3
L5-PT
Glia+UL
MG+DL
MG+UL
OPC
iN_1
Glia+iN
OL_2
MG
UMAP 2
UMAP 1
OL_1
L2/3
L4
L4/5-IT
L5-IT_1
L5-IT_2
L6-IT
L5-PT
L5/6-NP
L6-CT
mCtx
Clau
iN_1
iN_2
iN_3
iN_4
Str
Ambig
AG_1
AG_2
OPC
OL_1
OL_2
MG
Vasc
Glia+UL
Glia+iN
AG+DL
AG+N
MG+UL
MG+DL
C
Genotype
WT
Chd8+/−
Sex
female
male
Batch A
Batch B
WT-M-2
Het-M-2
WT-F-2
Het-F-2
WT-M-3
Het-M-3
WT-F-3
Het-F-3
D
Number UMI counts per nucleus
50000
40000
30000
20000
10000
0
Number genes detected per nucleus
10000
8000
6000
4000
2000
0
L2/3
L4
L4/5-IT
L5-IT_1
L5-IT_2
L6-IT
L5-PT
L5/6-NP
L6-CT
mCtx
Clau
iN_1
iN_2
iN_3
iN_4
Str
Ambig
AG_1
AG_2
OPC
OL_1
OL_2
MG
Vasc
Glia+UL
Glia+iN
AG+DL
AG+N
MG+UL
MG+DL

**Figure S16. Quality control for juvenile mouse cortex single-nucleus RNA-seq data.** (**A**) By-sample percentages of mitochondrial reads, with filtering threshold of 1% shown as a dotted line. (**B**) UMAP embedding of the full dataset of 73,775 nuclei passing the filter described in panel A, prior to doublet detection or other cluster-based filtering steps (Methods). (**C**) UMAP embedding from panel B colored by batch, sex, and genotype. (**D**) Violin plots showing total UMI counts (nUMI; *top*) and number of genes detected per nucleus (*bottom*) plotted for each cluster. Within each violin, the line corresponds to the median and the black diamond indicates the mean. See also Figure 7, Figure S17, and Tables S24-S25. L = layer of excitatory cortical neuron; IT = intratelencephalic; PT = pyramidal tract; NP = near-projecting; CT = corticothalamic; iN = inhibitory neuron; AG = astroglia; OL = oligodendrocyte; OPC = oligodendrocyte precursor cell; MG = microglia; Vasc = vasculature; mCtx = medial cortex; Clau = claustrum; Str = striatum; Ambig = ambiguous cluster; UL = upper-layer cortical neuron; DL = deep-layer cortical neuron; N = neuron.

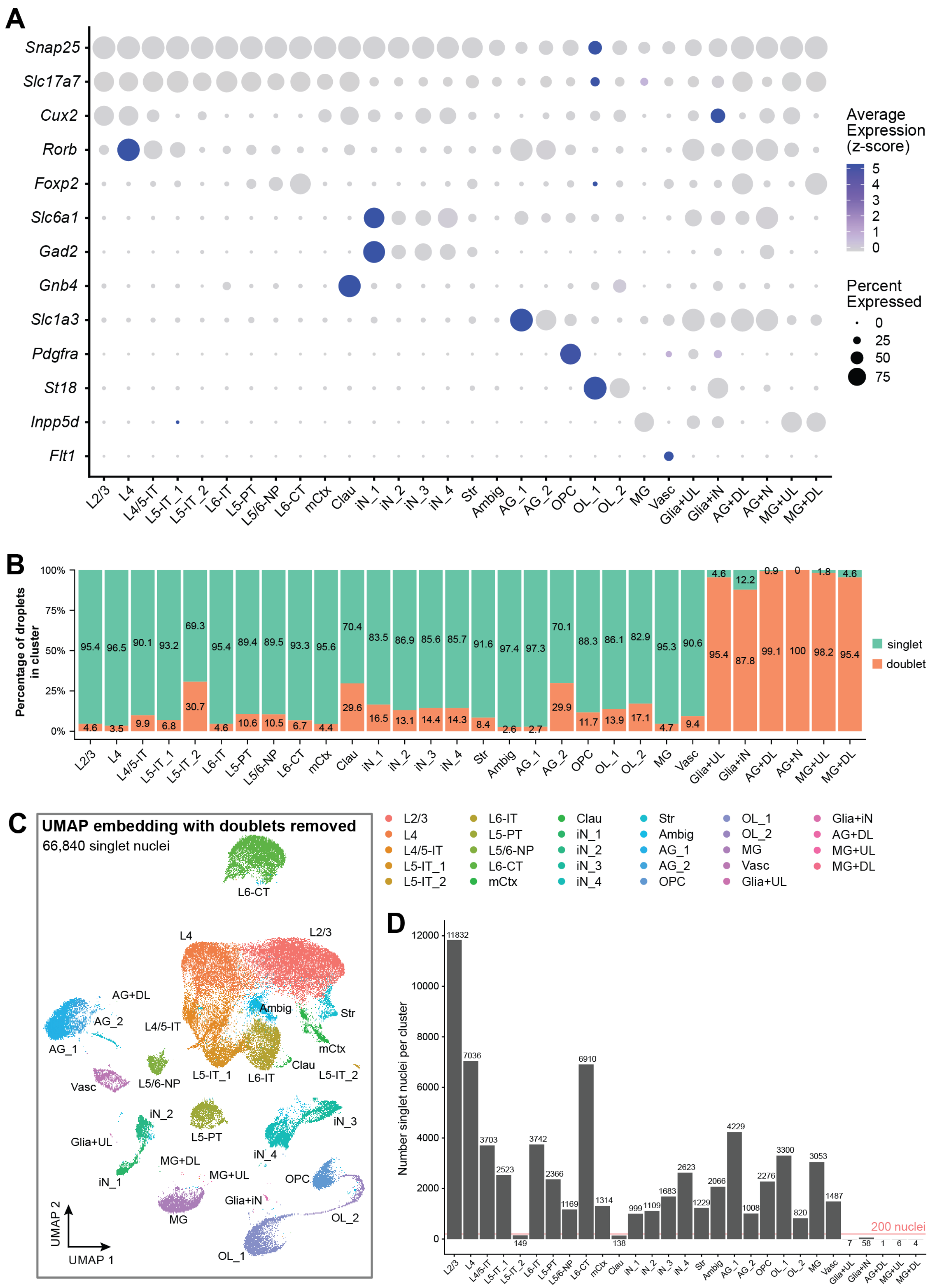
A
Snap25
Slc17a7
Cux2
Rorb
Foxp2
Slc6a1
Gad2
Gnb4
Slc1a3
Pdgfra
St18
Inpp5d
Flt1
Average Expression (z-score)
Percent Expressed
L2/3 L4 L4/5-IT L5-IT_1 L5-IT_2 L6-IT L5-PT L5/6-NP L6-CT mCtx Clau iN_1 iN_2 iN_3 iN_4 Str Ambig AG_1 AG_2 OPC OL_1 OL_2 MG Vasc Glia+UL Glia+iN AG+DL AG+N MG+UL MG+DL
B
Percentage of droplets in cluster
singlet
doublet
C
UMAP embedding with doublets removed
66,840 singlet nuclei
UMAP 1
UMAP 2
D
Number singlet nuclei per cluster
200 nuclei

**Figure S17. Broad cell type identification and doublet filtering for juvenile mouse cortex single-nucleus RNA-seq data.** (**A**) Dot plot showing expression of broad cell type marker genes for excitatory neurons, inhibitory neurons, astroglia, the oligodendrocyte lineage, microglia, and vasculature, prior to cluster-based filtering. (**B**) Percentage doublet and singlet composition per cluster, prior to cluster filtering, based on scDblFinder calls (Methods). (**C**) UMAP embedding of 66,840 singlet-only nuclei colored by cell type assignment (Methods). (**D**) Number of singlet nuclei per cluster, with the 200-nuclei threshold indicated by red line. See also Figure 7, Figure S16, and Tables S24-S27. L = layer of excitatory cortical neuron; IT = intratelencephalic; PT = pyramidal tract; NP = near-projecting; CT = corticothalamic; iN = inhibitory neuron; AG = astroglia; OL = oligodendrocyte; OPC = oligodendrocyte precursor cell; MG = microglia; Vasc = vasculature; mCtx = medial cortex; Clau = claustrum; Str = striatum; Ambig = ambiguous cluster; UL = upper-layer cortical neuron; DL = deep-layer cortical neuron; N = neuron.

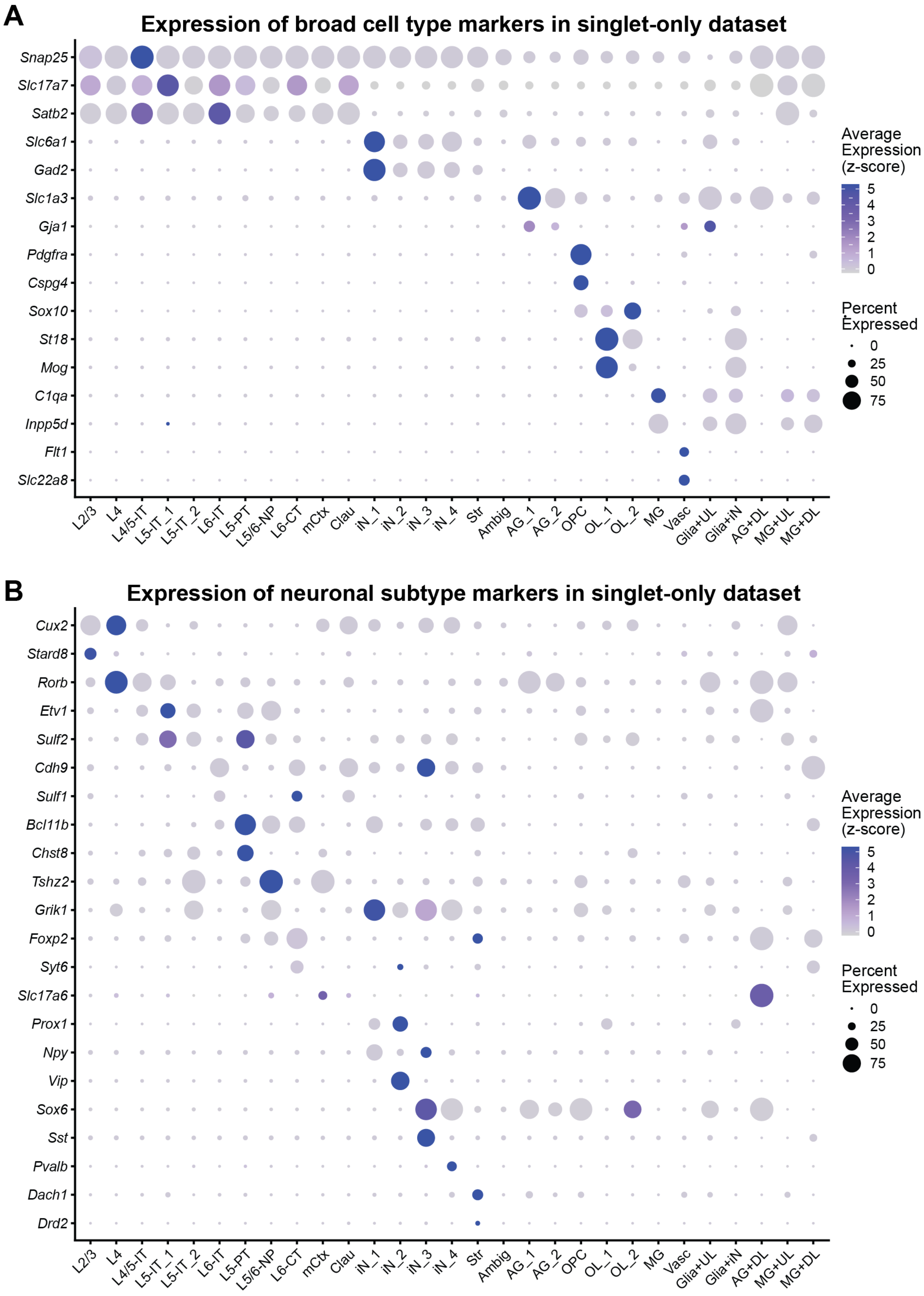
A
Expression of broad cell type markers in singlet-only dataset
Snap25
Slc17a7
Satb2
Slc6a1
Gad2
Slc1a3
Gja1
Pdgfra
Cspg4
Sox10
St18
Mog
C1qa
Inpp5d
Flt1
Slc22a8
L2/3
L4
L4/5-IT
L5-IT_1
L5-IT_2
L6-IT
L5-PT
L5/6-NP
L6-CT
mCtx
Clau
iN_1
iN_2
iN_3
iN_4
Str
Ambig
AG_1
AG_2
OPC
OL_1
OL_2
MG
Vasc
Glia+UL
Glia+iN
AG+DL
MG+UL
MG+DL
Average Expression (z-score)
5
4
3
2
1
0
Percent Expressed
0
25
50
75
B
Expression of neuronal subtype markers in singlet-only dataset
Cux2
Stard8
Rorb
Etv1
Sulf2
Cdh9
Sulf1
Bcl11b
Chst8
Tshz2
Grik1
Foxp2
Syt6
Slc17a6
Prox1
Npy
Vip
Sox6
Sst
Pvalb
Dach1
Drd2
L2/3
L4
L4/5-IT
L5-IT_1
L5-IT_2
L6-IT
L5-PT
L5/6-NP
L6-CT
mCtx
Clau
iN_1
iN_2
iN_3
iN_4
Str
Ambig
AG_1
AG_2
OPC
OL_1
OL_2
MG
Vasc
Glia+UL
Glia+iN
AG+DL
MG+UL
MG+DL
Average Expression (z-score)
5
4
3
2
1
0
Percent Expressed
0
25
50
75

**Figure S18. Expression of cell type marker genes in the singlet-only juvenile mouse cortex single-nucleus RNA-seq dataset.** (**A**-**B**) Dot plots showing expression of (**A**) broad cell type marker genes for excitatory cortical neurons, inhibitory neurons (iN), astroglia (AG), oligodendrocyte precursor cells (OPCs), oligodendrocytes (OL), microglia (MG), and vasculature (Vasc), or (**B**) neuronal subtype marker genes for subtypes of cortical excitatory neurons, subtypes of cortical inhibitory neurons, medial cortex (mCtx), and striatum (Str). See also Figure 7, Figures S16-S17, and Tables S26-S27. L = layer of excitatory cortical neuron; IT = intratelencephalic; PT = pyramidal tract; NP = near-projecting; CT = corticothalamic; Clau = claustrum; Ambig = ambiguous cluster; UL = upper-layer cortical neuron; DL = deep-layer cortical neuron.

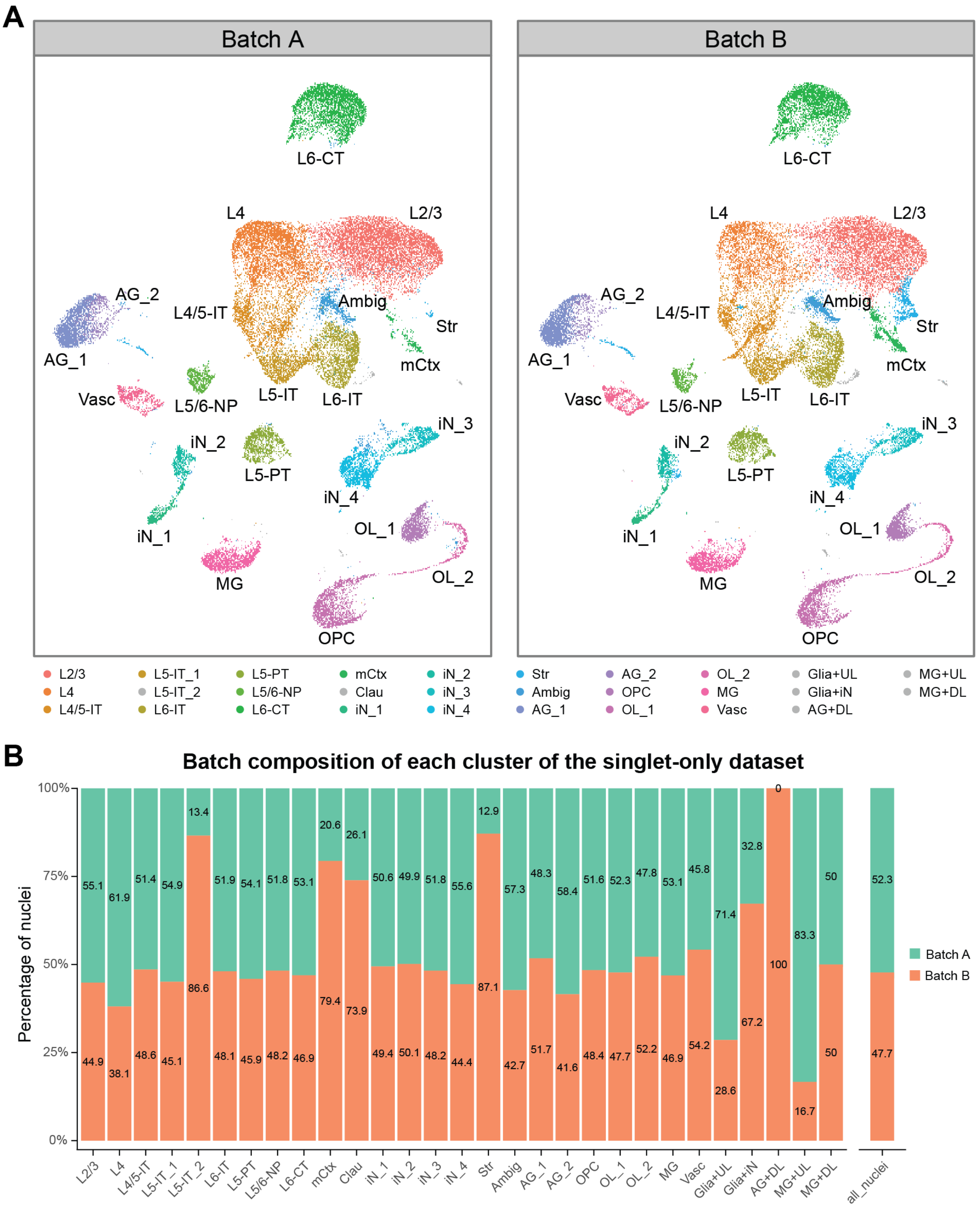

A
Batch A
Batch B
L6-CT
L4
L2/3
AG_2
L4/5-IT
Ambig
Str
AG_1
mCtx
Vasc
L5/6-NP
L5-IT
L6-IT
iN_3
iN_2
L5-PT
iN_4
iN_1
OL_1
OL_2
MG
OPC
L2/3 L4 L4/5-IT L5-IT_1 L5-IT_2 L6-IT L5-PT L5/6-NP L6-CT mCtx Clau iN_1 iN_2 iN_3 iN_4 Str Ambig AG_1 AG_2 OPC OL_1 OL_2 MG Vasc Glia+UL Glia+iN AG+DL MG+UL MG+DL
B
Batch composition of each cluster of the singlet-only dataset
Percentage of nuclei
100% 75% 50% 25% 0%
Batch A
Batch B
L2/3 55.1 44.9
L4 61.9 38.1
L4/5-IT 51.4 48.6
L5-IT_1 54.9 45.1
L5-IT_2 13.4 86.6
L6-IT 51.9 48.1
L5-PT 54.1 45.9
L5/6-NP 51.8 48.2
L6-CT 53.1 46.9
mCtx 20.6 79.4
Clau 26.1 73.9
iN_1 50.6 49.4
iN_2 49.9 50.1
iN_3 51.8 48.2
iN_4 55.6 44.4
Str 12.9 87.1
Ambig 57.3 42.7
AG_1 48.3 51.7
AG_2 58.4 41.6
OPC 51.6 48.4
OL_1 52.3 47.7
OL_2 47.8 52.2
MG 53.1 46.9
Vasc 45.8 54.2
Glia+UL 71.4 28.6
Glia+iN 32.8 67.2
AG+DL 0 100
MG+UL 83.3 16.7
MG+DL 50 50
all_nuclei 52.3 47.7

**Figure S19. Identification of clusters with biased batch contribution in the singlet-only juvenile mouse cortex single-nucleus RNA-seq dataset.** (**A**) UMAP embedding of 66,840 singlet-only nuclei colored by cell type assignment, split by batch (Methods). Clusters that did not pass the cluster size filter are shown in gray (Methods; Figure S17D). (**B**) Percentage batch contribution to each cluster in the singlet-only data, with the percentage batch contribution to the entire singlet-only dataset shown at far right (all_nuclei). See also Figure 7, Figure S17, and Tables S24 and S27. L = layer of excitatory cortical neuron; IT = intratelencephalic; PT = pyramidal tract; NP = near-projecting; CT = corticothalamic; iN = inhibitory neuron; AG = astroglia; OL = oligodendrocyte; OPC = oligodendrocyte precursor cell; MG = microglia; Vasc = vasculature; mCtx = medial cortex; Clau = claustrum; Str = striatum; Ambig = ambiguous cluster; UL = upper-layer cortical neuron; DL = deep-layer cortical neuron.

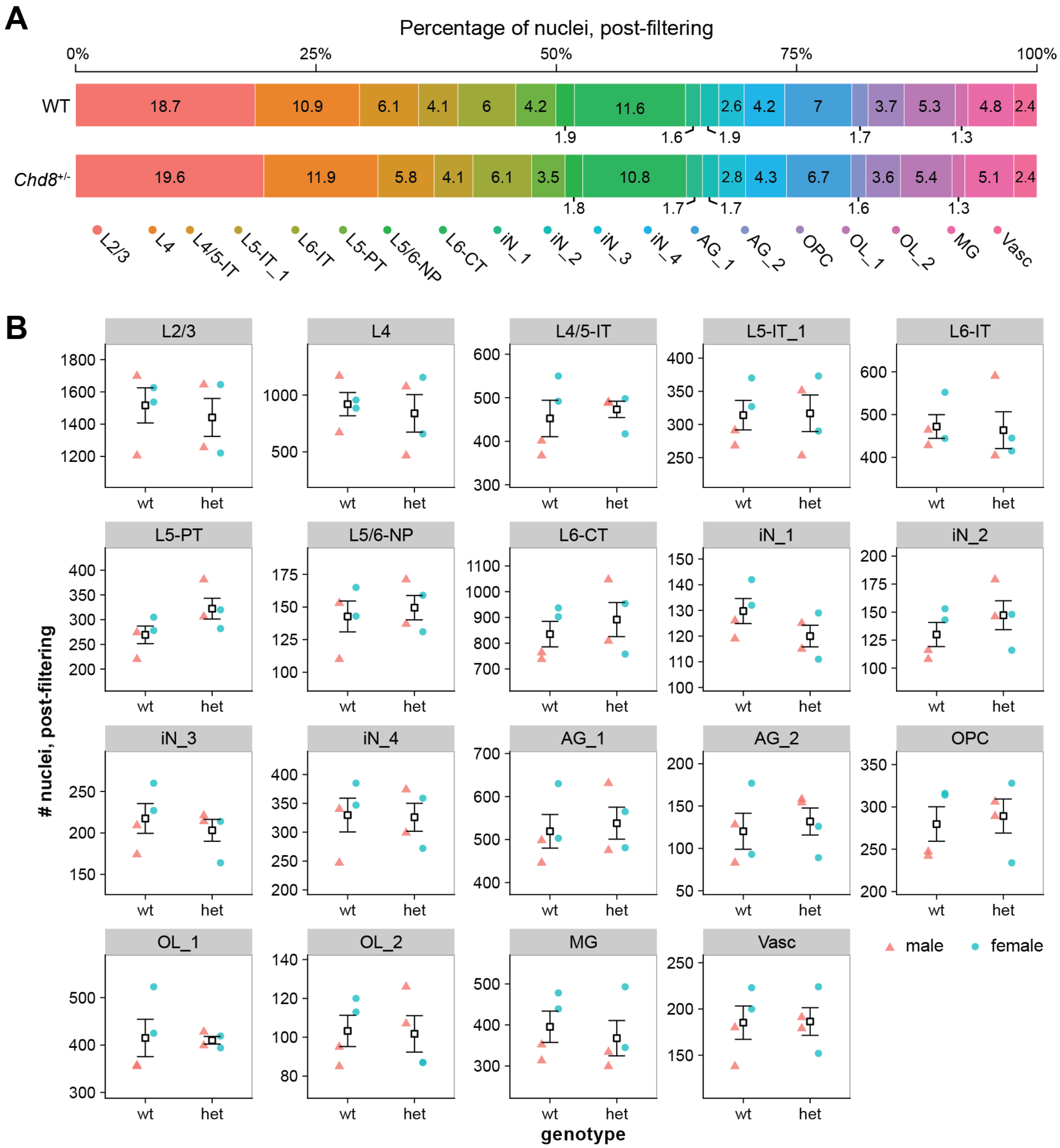


**Figure S20. By-genotype comparison of cluster composition of the singlet-only juvenile cortex single-nucleus RNA-seq data** (**A**) Percentage cluster contribution to the singlet-only wild type (WT) and *Chd8*$^{+/-}$ juvenile mouse cortex data, excluding clusters that did not pass the filtering criteria described in Figures S16-S19 and Methods. (**B**) Comparison of the number of singlet-only nuclei within each cluster between the wild type (wt) and *Chd8*$^{+/-}$ (het) juvenile mouse cortex. Data are represented as mean ± SEM, with datapoints for individual replicates color-coded by sex. *P*-values were calculated by two-tailed Welch's *t*-test (Methods). See also Figure 7, Figures S16-S19, and Tables S27-S28. L = layer of excitatory cortical neuron; IT = intratelencephalic; PT = pyramidal tract; NP = near-projecting; CT = corticothalamic; iN = inhibitory neuron; AG = astroglia; OL = oligodendrocyte; OPC = oligodendrocyte precursor cell; MG = microglia; Vasc = vasculature.

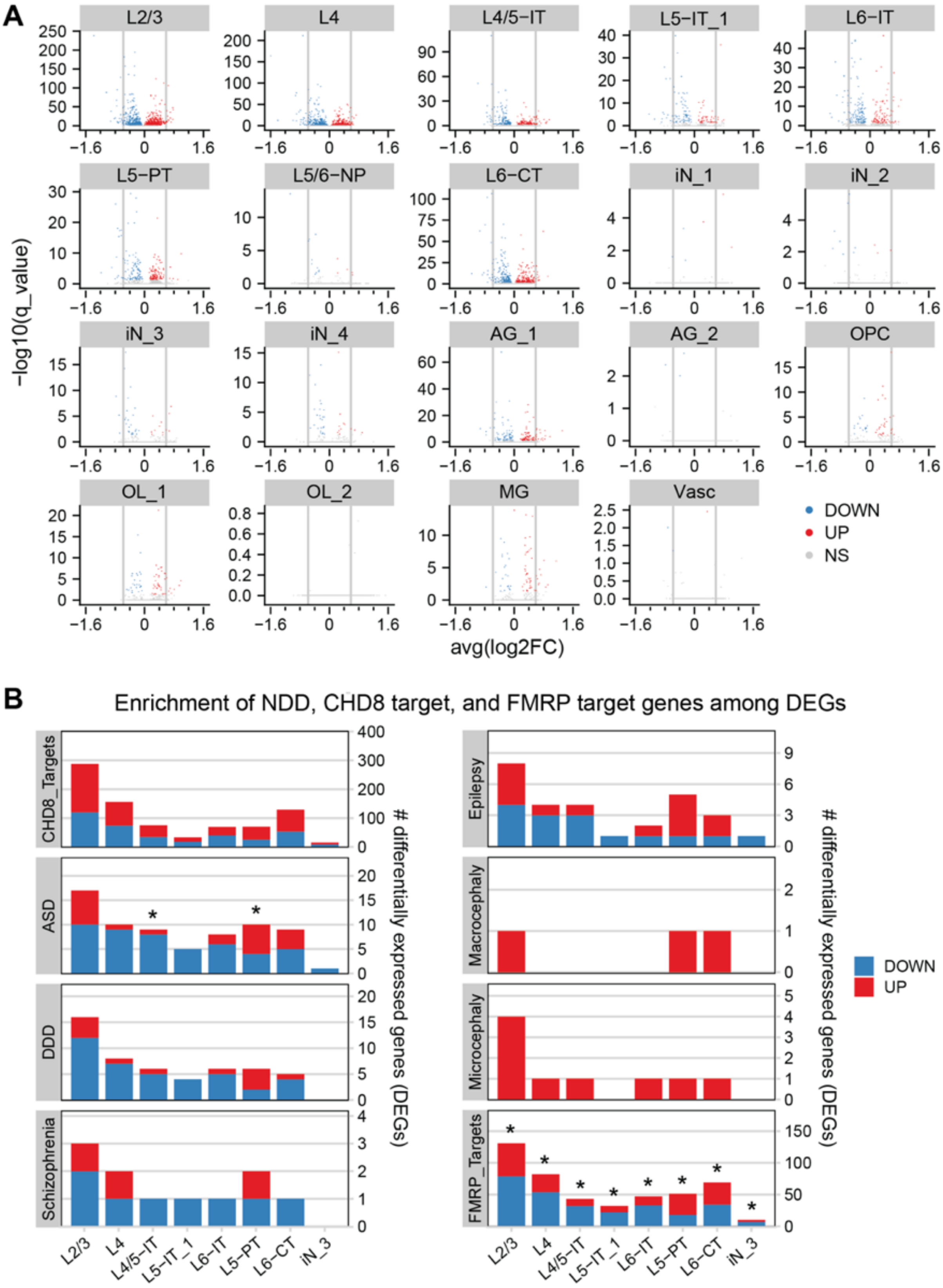
A
L2/3
L4
L4/5−IT
L5−IT_1
L6−IT
L5−PT
L5/6−NP
L6−CT
iN_1
iN_2
iN_3
iN_4
AG_1
AG_2
OPC
OL_1
OL_2
MG
Vasc
−log10(q_value)
avg(log2FC)
DOWN
UP
NS
B
Enrichment of NDD, CHD8 target, and FMRP target genes among DEGs
CHD8_Targets
ASD
DDD
Schizophrenia
Epilepsy
Macrocephaly
Microcephaly
FMRP_Targets
### differentially expressed genes (DEGs)
L2/3
L4
L4/5−IT
L5−IT_1
L6−IT
L5−PT
L6−CT
iN_3
DOWN
UP

**Figure S21. Differential gene expression by cell type and enrichment of CHD8 target genes and neurodevelopmental disorder (NDD)-associated genes among differentially expressed genes by cell type in the filtered wild type and *Chd8*$^{+/-}$ juvenile mouse cortex single-nucleus RNA-seq dataset.** (**A**) Volcano plots of by-cluster differential gene expression between the wild type and *Chd8*$^{+/-}$ juvenile mouse cortex, determined by Monocle 3, with genes color-coded by significant downregulation (DOWN; blue), significant upregulation (UP; red), or not significantly different (NS; gray) (Methods). Vertical gray lines = ±log2(1.5 fold-change). (**B**) Bar charts of the intersection between differentially expressed genes of each cluster and NDD-associated gene sets, CHD8 target genes, or FMRP target genes. Bars are color-coded by the number of downregulated (DOWN; blue) or upregulated (UP; red) genes within each intersection. Significance was determined by one-way Fisher's exact test, with adjustment for multiple testing (Methods); * = Benjamini Hochberg-adjusted *p*-value < 0.05 (Table S31). Only clusters with at least one significant enrichment result are shown. L = layer of excitatory cortical neuron; IT = intratelencephalic; PT = pyramidal tract; NP = near-projecting; CT = corticothalamic; iN = inhibitory neuron; AG = astroglia; OL = oligodendrocyte; OPC = oligodendrocyte precursor cell; MG = microglia; Vasc = vasculature. ASD = autism spectrum disorder risk-associated genes, DDD = Deciphering Developmental Disorders gene set. See also Figure 7, Table S9, and Tables S29-S31.

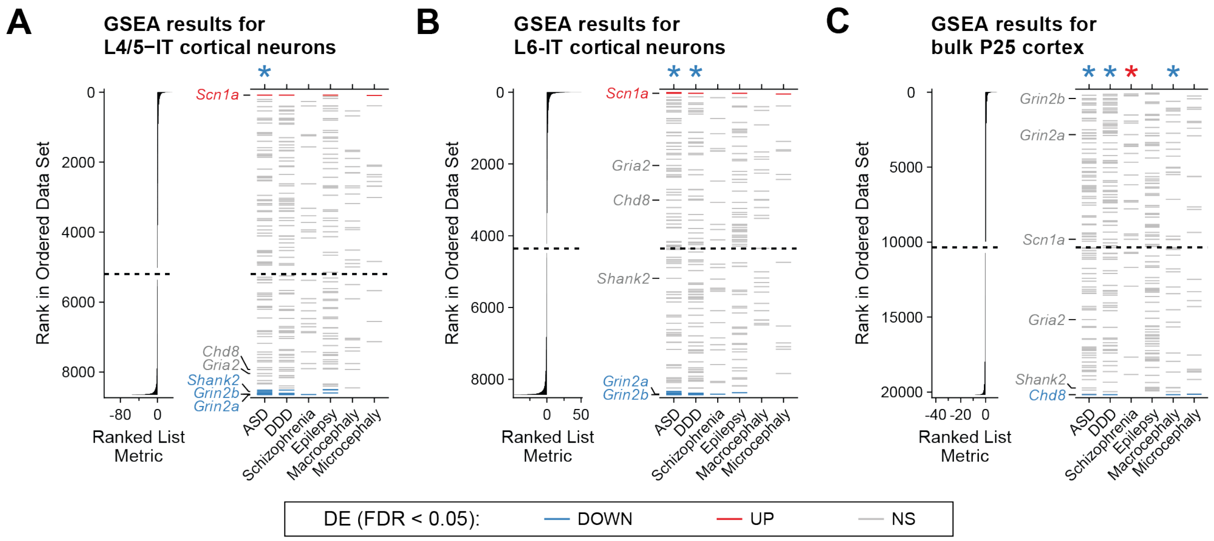


**Figure S22. Gene set enrichment analysis (GSEA) of differential gene expression in the wild type and *Chd8*$^{+/-}$ juvenile cortex, assessing neurodevelopmental disorder (NDD), CHD8 target, and FMRP target gene sets.** (**A-B**) GSEA results for (**A**) L4/5-IT and (**B**) L6-IT clusters for the wild type vs. *Chd8*$^{+/-}$ juvenile cortex single-nucleus RNA-sequencing data. For each cell type, input genes are ordered by a "Ranked List Metric" calculated from the Monocle 3 output equal to sign(avg_log2FC) * -log10(*p*-value). Genes in the ranked list that overlap each NDD gene set are shown as horizontal ticks and color-coded by differential expression calls determined by Monocle 3 (red = upregulated, UP; blue = downregulated, DOWN; gray = not significantly different, NS; Methods). The dotted line indicates the zero-cross rank separating positive and negative values. * = FDR < 0.05. (**C**) GSEA results for the wild type vs. *Chd8*$^{+/-}$ juvenile cortex bulk RNA-sequencing data comparison (Methods). Input genes are ordered by a "Ranked List Metric" calculated from the DESeq2 output equal to sign(avg_log2FC) * -log10(*p*-value) (Methods). Genes in the ranked list that overlap each NDD gene set are shown as horizontal ticks and color-coded by differential expression calls determined by DESeq2 (Methods). See also Tables S9, S30, S32, S35, and S37. L = layer of excitatory cortical neuron; IT = intratelencephalic. ASD = autism spectrum disorder risk-associated genes; DDD = Deciphering Developmental Disorders gene set. * = FDR < 0.05.

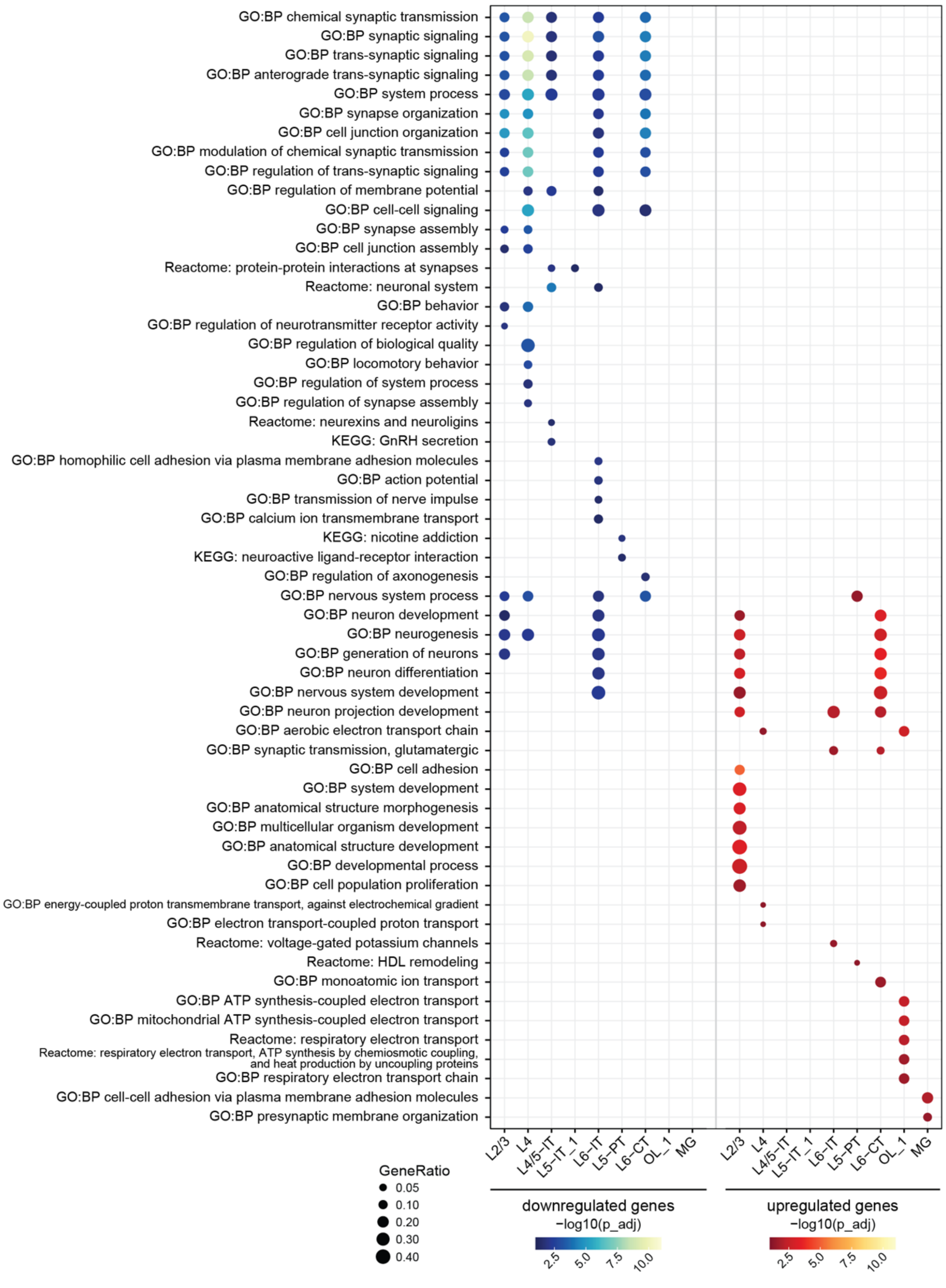

GO:BP chemical synaptic transmission
GO:BP synaptic signaling
GO:BP trans-synaptic signaling
GO:BP anterograde trans-synaptic signaling
GO:BP system process
GO:BP synapse organization
GO:BP cell junction organization
GO:BP modulation of chemical synaptic transmission
GO:BP regulation of trans-synaptic signaling
GO:BP regulation of membrane potential
GO:BP cell-cell signaling
GO:BP synapse assembly
GO:BP cell junction assembly
Reactome: protein-protein interactions at synapses
Reactome: neuronal system
GO:BP behavior
GO:BP regulation of neurotransmitter receptor activity
GO:BP regulation of biological quality
GO:BP locomotory behavior
GO:BP regulation of system process
GO:BP regulation of synapse assembly
Reactome: neurexins and neuroligins
KEGG: GnRH secretion
GO:BP homophilic cell adhesion via plasma membrane adhesion molecules
GO:BP action potential
GO:BP transmission of nerve impulse
GO:BP calcium ion transmembrane transport
KEGG: nicotine addiction
KEGG: neuroactive ligand-receptor interaction
GO:BP regulation of axonogenesis
GO:BP nervous system process
GO:BP neuron development
GO:BP neurogenesis
GO:BP generation of neurons
GO:BP neuron differentiation
GO:BP nervous system development
GO:BP neuron projection development
GO:BP aerobic electron transport chain
GO:BP synaptic transmission, glutamatergic
GO:BP cell adhesion
GO:BP system development
GO:BP anatomical structure morphogenesis
GO:BP multicellular organism development
GO:BP anatomical structure development
GO:BP developmental process
GO:BP cell population proliferation
GO:BP energy-coupled proton transmembrane transport, against electrochemical gradient
GO:BP electron transport-coupled proton transport
Reactome: voltage-gated potassium channels
Reactome: HDL remodeling
GO:BP monoatomic ion transport
GO:BP ATP synthesis-coupled electron transport
GO:BP mitochondrial ATP synthesis-coupled electron transport
Reactome: respiratory electron transport
Reactome: respiratory electron transport, ATP synthesis by chemiosmotic coupling, and heat production by uncoupling proteins
GO:BP respiratory electron transport chain
GO:BP cell-cell adhesion via plasma membrane adhesion molecules
GO:BP presynaptic membrane organization
L2/3
L4
L4/5−IT
L5−IT_1
L6−IT
L5−PT
L6−CT
OL_1
MG
L2/3
L4
L4/5−IT
L5−IT_1
L6−IT
L5−PT
L6−CT
OL_1
MG
GeneRatio
0.05
0.10
0.20
0.30
0.40
downregulated genes
−log10(p_adj)
2.5
5.0
7.5
10.0
upregulated genes
−log10(p_adj)
2.5
5.0
7.5
10.0

**Figure S23. Dot plot of by-cluster functional term enrichment among downregulated and upregulated differentially expressed genes (DEGs) in the *Chd8*$^{+/-}$ juvenile cortex.** Enrichment was determined by g:Profiler (Methods). Only clusters enriching for at least one functional term are shown. Dot size corresponds to the ratio of downregulated or upregulated genes intersecting with the denoted functional term out of all downregulated or upregulated genes of that cluster submitted as a query to g:Profiler (GeneRatio). Dots are colored by significance of enrichment and direction of differential expression (blue color scale = downregulated DEG enrichment, g:SCS-adjusted *p*-value < 0.05; red color scale = downregulated DEG enrichment, g:SCS-adjusted *p*-value < 0.05). p_adj = g:SCS-adjusted *p*-value calculated by g:Profiler. See also Figure 7 and Tables S33-S34. L = layer of excitatory cortical neuron; IT = intratelencephalic; PT = pyramidal tract; CT = corticothalamic; OL = oligodendrocyte; MG = microglia.

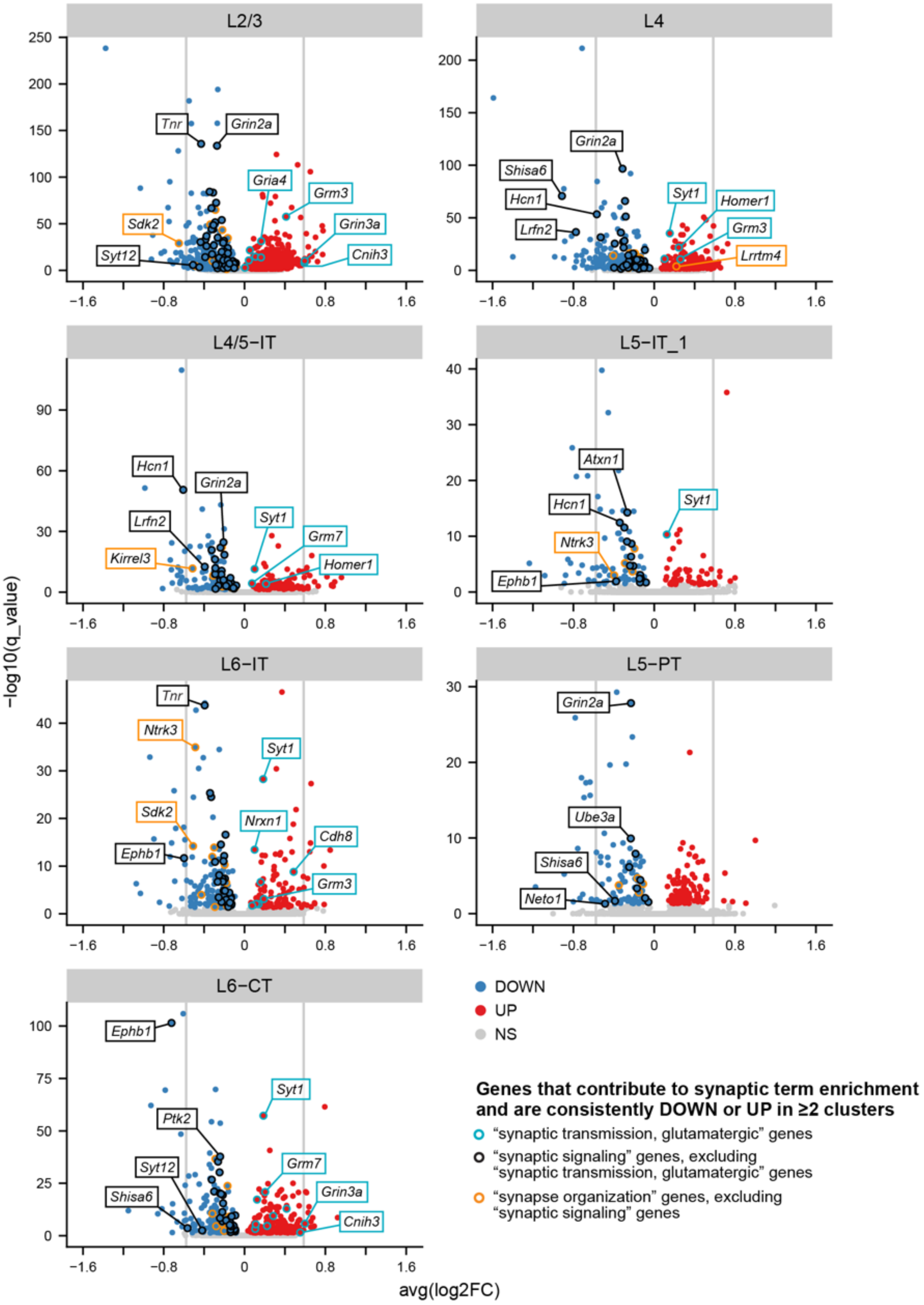

L2/3
L4
L4/5−IT
L5−IT_1
L6−IT
L5−PT
L6−CT
−log10(q_value)
avg(log2FC)
Tnr
Grin2a
Gria4
Grm3
Sdk2
Grin3a
Syt12
Cnih3
Shisa6
Hcn1
Lrfn2
Syt1
Homer1
Lrrtm4
Kirrel3
Grm7
Atxn1
Ntrk3
Ephb1
Nrxn1
Cdh8
Ube3a
Neto1
Ptk2
DOWN
UP
NS
Genes that contribute to synaptic term enrichment and are consistently DOWN or UP in ≥2 clusters
"synaptic transmission, glutamatergic" genes
"synaptic signaling" genes, excluding "synaptic transmission, glutamatergic" genes
"synapse organization" genes, excluding "synaptic signaling" genes

**Figure S24. Differential expression of consistently downregulated and upregulated synaptic term genes across excitatory neuron subtypes in the *Chd8*$^{+/-}$ juvenile cortex.** Volcano plots of the Monocle 3 differential gene expression results for excitatory neuronal clusters (Methods). Fill color for each gene indicates significant downregulation (DOWN; blue), significant upregulation (UP; red), or not significantly different (NS; gray). Outline color indicates differentially expressed genes (DEGs) that both contribute to the enrichment of synaptic functional terms identified by GO:BP term analysis ("synaptic transmission, glutamatergic," blue; "synaptic signaling," black; "synapse organization," orange) and were consistently downregulated or upregulated across ≥2 excitatory neuronal clusters. A subset of DEGs contributed to enrichment for multiple synaptic GO:BP terms. DEGs in both the "synaptic transmission, glutamatergic" and "synaptic signaling" term lists are color-coded as "synaptic transmission, glutamatergic" genes. DEGs in both the "synaptic signaling" and "synapse organization" lists are color-coded as "synaptic signaling" genes. For each cluster, the two most significantly downregulated or upregulated synaptic term genes are labeled, along with the two downregulated and upregulated synaptic term genes with effect sizes of the greatest magnitude. See also Figure 7, Figures S23 and S25, Table S30, and Tables S33-S34. Vertical gray lines = ±log2(1.5 fold-change); L = layer of excitatory cortical neuron; IT = intratelencephalic; PT = pyramidal tract; CT = corticothalamic.

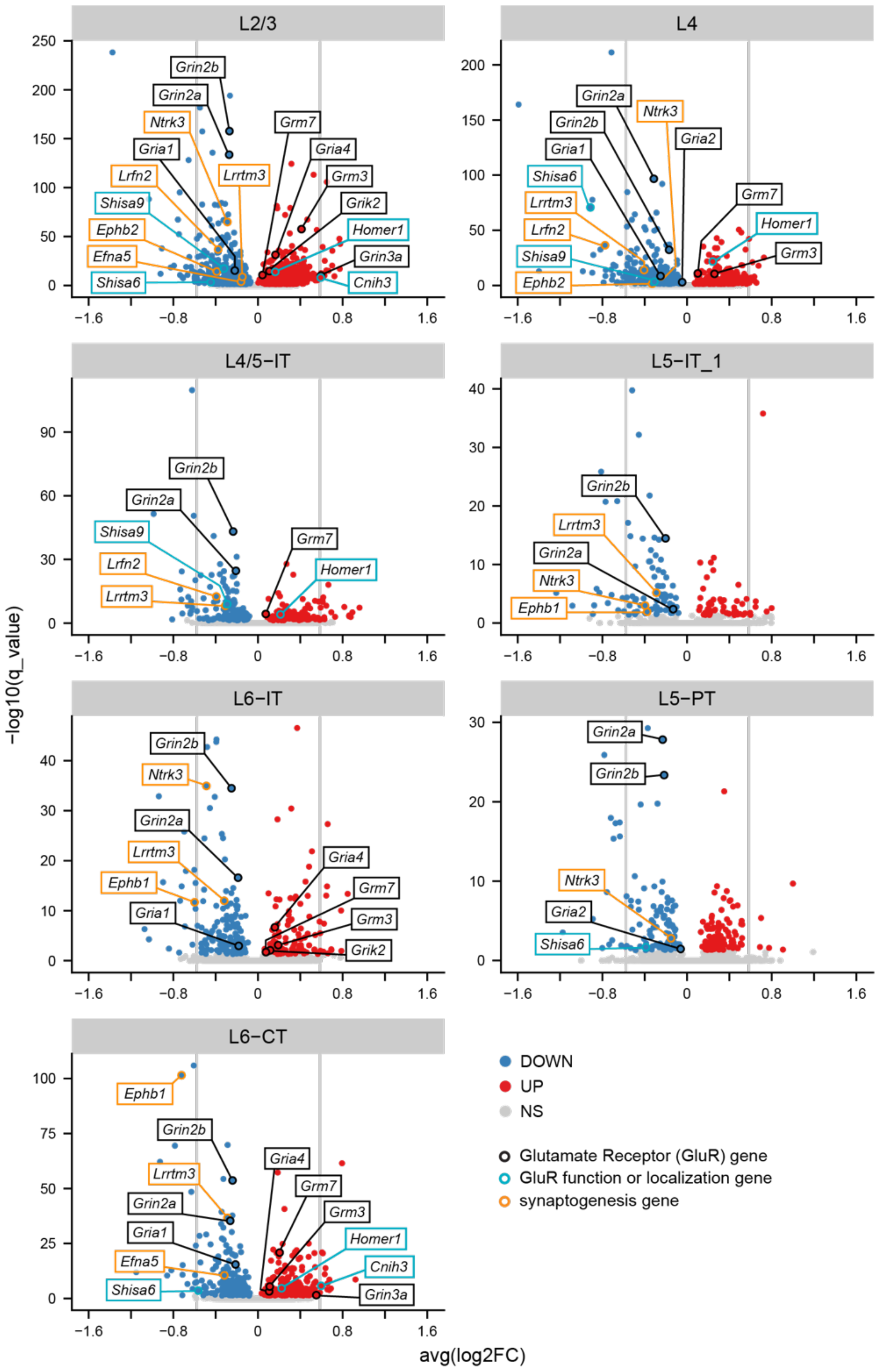
L2/3
L4
L4/5−IT
L5−IT_1
L6−IT
L5−PT
L6−CT
Grin2b
Grin2a
Ntrk3
Gria1
Lrfn2
Shisa9
Ephb2
Efna5
Shisa6
Lrrtm3
Grm7
Gria4
Grm3
Grik2
Homer1
Grin3a
Cnih3
Gria2
Ephb1
−log10(q_value)
avg(log2FC)
DOWN
UP
NS
Glutamate Receptor (GluR) gene
GluR function or localization gene
synaptogenesis gene

**Figure S25. Differential expression of consistently downregulated and upregulated genes encoding glutamate receptors (GluRs), regulators of GluRs, or regulators of synaptogenesis across excitatory neuron subtypes in the *Chd8*$^{+/-}$ juvenile cortex.** Volcano plots of the Monocle 3 differential gene expression results for excitatory neuronal clusters (Methods). Fill color for each gene indicates significant downregulation (DOWN; blue), significant upregulation (UP; red), or not significantly different (NS; gray). A subset of consistently downregulated or upregulated genes that encode glutamate receptors (GluRs), regulators of GluR localization and/or function, or synaptogenesis are outlined in black, light blue, or orange, respectively. These genes are a subset of those outlined in Figure S24. See also Figure 7, Figures S23-S24, Table S30, and Tables S33-S34. Vertical gray lines = ±log2(1.5 fold-change); L = layer of excitatory cortical neuron; IT = intratelencephalic; PT = pyramidal tract; CT = corticothalamic.
